# Engineering Biosensors to Enhance Monoterpene Indole Alkaloid Production in Yeast

**DOI:** 10.64898/2026.06.01.729246

**Authors:** Maxence Holtz, Simon D’Oelsnitz, Cecília Castellví Domingo, Niklas G. Madsen, Mars Yu Hong, Jonathan Asmund Arnesen, Aafke C. A. van Aalst, Samantha de Haan, Christopher K. Weingarten, Ditte H. Welner, Pamela A. Silver, Y. Jessie Zhang, Michael K. Jensen, Carlos G. Acevedo-Rocha

## Abstract

Monoterpene Indole Alkaloids (MIAs) are a diverse family of plant natural products with various medicinal applications. Although MIAs, such as vinblastine and reserpine, are clinically validated, sourcing of MIAs for clinical use or drug discovery from natural resources or via chemical synthesis is hampered due to their scarcity and chemical complexity. Refactoring MIA biosynthesis pathways in microbial cell factories could offer an alternative, more stable and potentially sustainable manufacturing route for alkaloid medicines and novel therapies. However, reaching commercially attractive titers, rates and yields remains challenging owing to the length and complexity of these metabolic pathways. One critical bottleneck is the low screening throughput and very high cost of the analytical methods used to quantify MIA for optimizing production. In this study, we evolved RamR, a promiscuous bacterial transcription factor to respond to five different MIAs, resulting in highly sensitive and selective sensor variants (EC_50_<10 μM). X-ray crystallography and computational modeling provided insight into the MIA binding of the evolved biosensor variants. The RamR biosensing platform was functionalized in yeast and subsequently applied in a cost-effective semi-throughput screening campaign of a 188-gene overexpression library to identify high-performing cell factory designs for strictosidine, the common precursor for all MIAs. The fluorescent biosensor signal correlated with HPLC quantification (r^2^ = 0.932) allowing identification of single metabolic engineering hits which when combined yielded a maximum titer of >220 mg/L strictosidine, 3-fold higher than the parental reference strain. This study demonstrates the development of selective biosensors for MIAs and the cost-effective identification of novel metabolic engineering hits for optimizing MIA production in microbial cell factories.

## Introduction

Plants have served as a source of highly potent therapeutic agents, forming the basis of numerous treatments for human diseases throughout history. Monoterpene indole alkaloids (MIAs) are an important family of sprecialized metabolites produced by *Gentianales* plants, with over 3,000 known compounds exhibiting a variety of bioactivities and medicinal uses [1]. Prominent examples include vincristine, vinblastine, and camptothecin analogs, which are essential in cancer treatment, while ajmaline and reserpine are utilized in cardiovascular therapies [1].

Supplying MIAs at scale for pharmaceutical applications is challenged by the low metabolite accumulation *in planta* and generally poor extraction yields, resulting in high prices and frequent shortages for such essential drugs sourced from plant material [2]. Due to the complex molecular architecture of these alkaloids, including multiple chiral centers, total chemical synthesis is not viable on a commercial scale [3]. There is therefore a strong interest in refactoring MIA production in heterologous hosts to decrease the cost of MIA-based therapies, discover novel clinical candidates, and improve the supply chain stability of MIAs.

*Saccharomyces cerevisiae* has emerged as the leading microbial platform for MIA biosynthesis, combining a well-established synthetic biology toolbox with eukaryotic cellular features that support post-translational modifications, organelle-based compartmentalization, and the functional expression of complex plant enzymes such as cytochrome P450 monoxygenases [4]. Across diverse plant lineages, the biosynthesis of MIAs initiates with the formation of strictosidine from tryptamine and geranyl diphosphate-derived secologanin; this key intermediate gives rise to the vast MIA family via diverse tailoring reactions (**Supp. Fig. 1**) [1,5]. Elucidation of plant MIA biosynthetic pathways has enabled both the development of a strictosidine-producing yeast platform and the stepwise reconstruction of downstream pathway branches in yeast leading to the production of diverse MIAs such as alstonine, serpentine, mitragynine, catharanthine and vindoline, the latter two serving as advanced pharmaceutical ingredients for the industrial semi-synthesis of vinblastine and vincristine [2,4,6–9]. In *S. cerevisiae*, reported MIA titers span several orders of magnitude, from >800 mg/L achieved recently for the central intermediate strictosidine [10], to ∼160 mg/L for catharanthine enabled by pathway scaffolding strategies [11], whereas documented titer for other downstream alkaloids, such as rauwolscine or mitragynine, remain <1 mg/L.

Strain engineering aimed at enhancing titer, rate, and yield to meet commercial benchmarks is a laborious and costly endeavor, frequently requiring years of sequential optimization efforts [12,13]. In addition, enzymatic bottlenecks and shunt products have been consistently observed in published yeast MIA pathway refactoring efforts, typically involving low activity, poor expression, or substrate promiscuity, reflecting the evolutionary constraints of secondary metabolism enzymes which are not optimized to support high-flux [14]. Similarly, optimization of MIA cell factories remains slow and costly due to reliance on low-throughput analytical methods, underscoring the urgent need for versatile, high-throughput and cost-effective tools to accelerate MIA enzyme and metabolic engineering [12].

In this study, we evolved the highly promiscuous allosteric transcription factor (aTF) RamR from *Salmonella typhimurium* to respond to five different MIAs of industrial and medicinal interest using an established directed evolution workflow [15]. After functionalization in yeast, these biosensors were applied as high-throughput screening tools to optimize strictosidine-producing cell factories. This work establishes the first biosensor-enabled platform for MIA pathway engineering in yeast, providing a scalable framework to accelerate strain development toward commercially viable production platforms.

## Results

### RamR as a scaffold to engineer MIA biosensors

Aiming for broad applicability, we selected five therapeutically and structurally diverse MIAs as target ligands to develop biosensors for: strictosidine (STR, central precursor to all MIAs), rauwolscine (RAU, alpha-2 adrenergic receptor antagonist), catharanthine (CAT, precursor of cancer therapeutic vinblastine), mitragynine (MIT, pain management, addiction, and depression) and quinine (QUI, antimalaria and flavoring agent of tonic water) (**Fig. 1A**). STR, RAU, MIT and CAT and their derivatives have important pharmacological applications, fully elucidated biosynthetic pathways and have been produced in yeast before [2,7,9]. QUI was selected to provide a screening tool to assist ongoing pathway discovery efforts [16–18].

**Figure 1:**
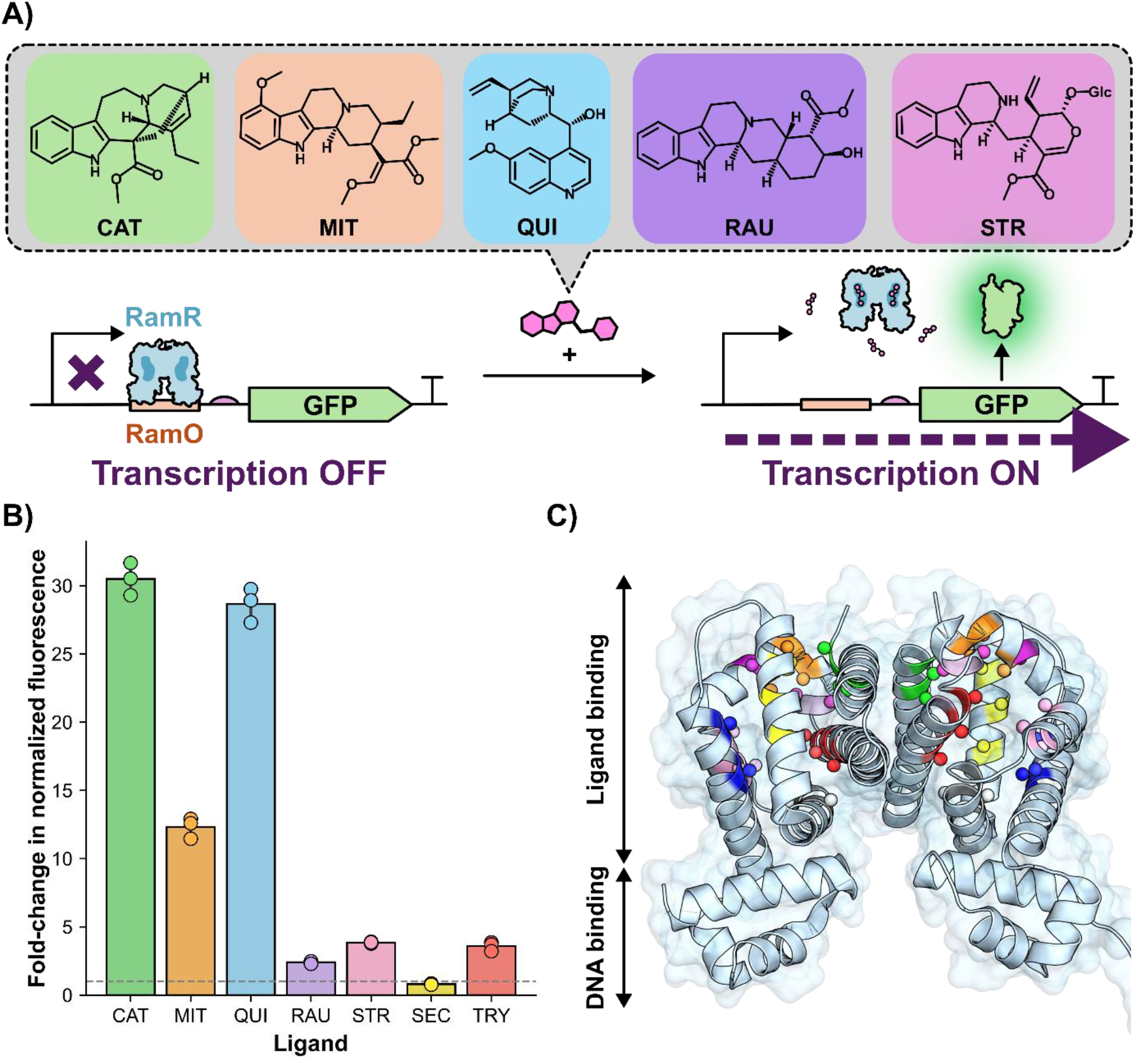
RamR as a scaffold for MIA biosensing. **A)** Illustration of the RamR biosensing mechanism with structures of the MIA ligands included in this study. In the absence of ligand, RamR associates with the RamO operator, blocking transcription. Ligand binding induces a conformational change that disrupts DNA binding, allowing reporter gene transcription to proceed. **B)**Fluorescence response fold-change of WT RamR with the five target MIAs supplemented at 100 µM compared to sensor background signal (DMSO 1%). Measurements for each condition represent the average of three biological replicates. Error bars represent S.D.+/−the mean. **C)** Overall structure of WT RamR (PDB 3VVX) with colored residues targeted for site-saturation mutagenesis in the ligand binding pocket. Full library information is available in **Supp. Fig. 2**.

RamR is a TetR-family transcriptional repressor from *S. typhimurium* that naturally controls the expression of multidrug efflux pump. It possesses a large hydrophobic binding pocket with flexible aromatic residues and its promiscuity and evolvability has been harnessed to develop highly sensitive and selective biosensors for plant benzylisoquinoline alkaloids (BIAs), *Amaryllidaceae* alkaloids (AAs), and chiral amines [15,19,20]. To evaluate the potential of RamR as a scaffold for engineering MIA-responsive biosensors (**Fig. 1B**), we examined the response of the wild-type (WT) protein to the five selected ligands in *Escherichia coli* using the previously described biosensing plasmids pReg-RamR and pramr-GFP [15]. RamR exhibited a ligand-dependent activation, with normalized GFP outputs spanning a 2.2-to 29.2-fold increase upon induction at 100 µM (**Fig. 1C**). To further characterize RamR specificity for MIA pathway intermediates, we tested MIA pathway precursors secologanin and tryptamine. Secologanin showed no detectable activation, whereas tryptamine triggered a 3.6-fold GFP induction when supplemented at 1 mM, underscoring the necessity of counterselection for tryptamine sensitivity during biosensor engineering.

### Evolution of RamR towards MIAs

The initial responsiveness of RamR toward MIAs was promising; however, further optimization of its sensitivity and selectivity was essential for both enzyme and metabolic engineering. We implemented the SELIS (Seamless Enrichment of Ligand Inducible Sensors) directed evolution workflow [15] to evolve RamR biosensors for MIAs. This method combines two stages: first, a selection step, in which only variants capable of promoter repression support cell growth, and second, a fluorescence-based screen to identify those that activate in the presence of the target molecule. As part of this workflow, nine site-saturation mutagenesis (SSM) libraries were generated using NNS codons to diversify clusters of three neighboring residues in RamR’s ligand-binding pocket (**Fig. 1C, Supp. Fig. 2**). Five of them came from previous BIA biosensor engineering work [15]. The other four were designed by combining molecular docking of target ligands, tunnel analysis with CAVER [21], and residue mutability scores obtained with Hotspot Wizard [22]. Each SSM library encoded approximately 32,000 unique genotypes (32×32×32), equivalent to 8,000 unique amino acid combinations (20×20×20), and these were pooled prior to selection and screening, enabling evaluation of up to ∼295,000 distinct variants per round of evolution. Additionally, each SELIS round incorporated an error-prone PCR (epPCR) library targeting the full coding sequence, introducing an average of two mutations per gene.

We completed two rounds of evolution for MIT, RAU, and CAT, and one round for QUI and STR (**Fig. 2A-E, Table 1**). All campaigns included 1 mM tryptamine counterselection to eliminate off-target responsiveness. A relatively highly-sensitive QUI1 variant (EC_50_ = 2.5 µM) was obtained after just one round and three mutations. STR evolution was restricted to a single round due to limited compound supply, as it is not commercially available. The evolved sensors QUI1, CAT2, MIT2, and RAU2 exhibited an EC_50_ <10 µM. For STR1, EC_50_ determination was not possible due to incomplete signal saturation. Nevertheless, a near-linear response was observed between 25 and 100 µM.

**Figure 2:**
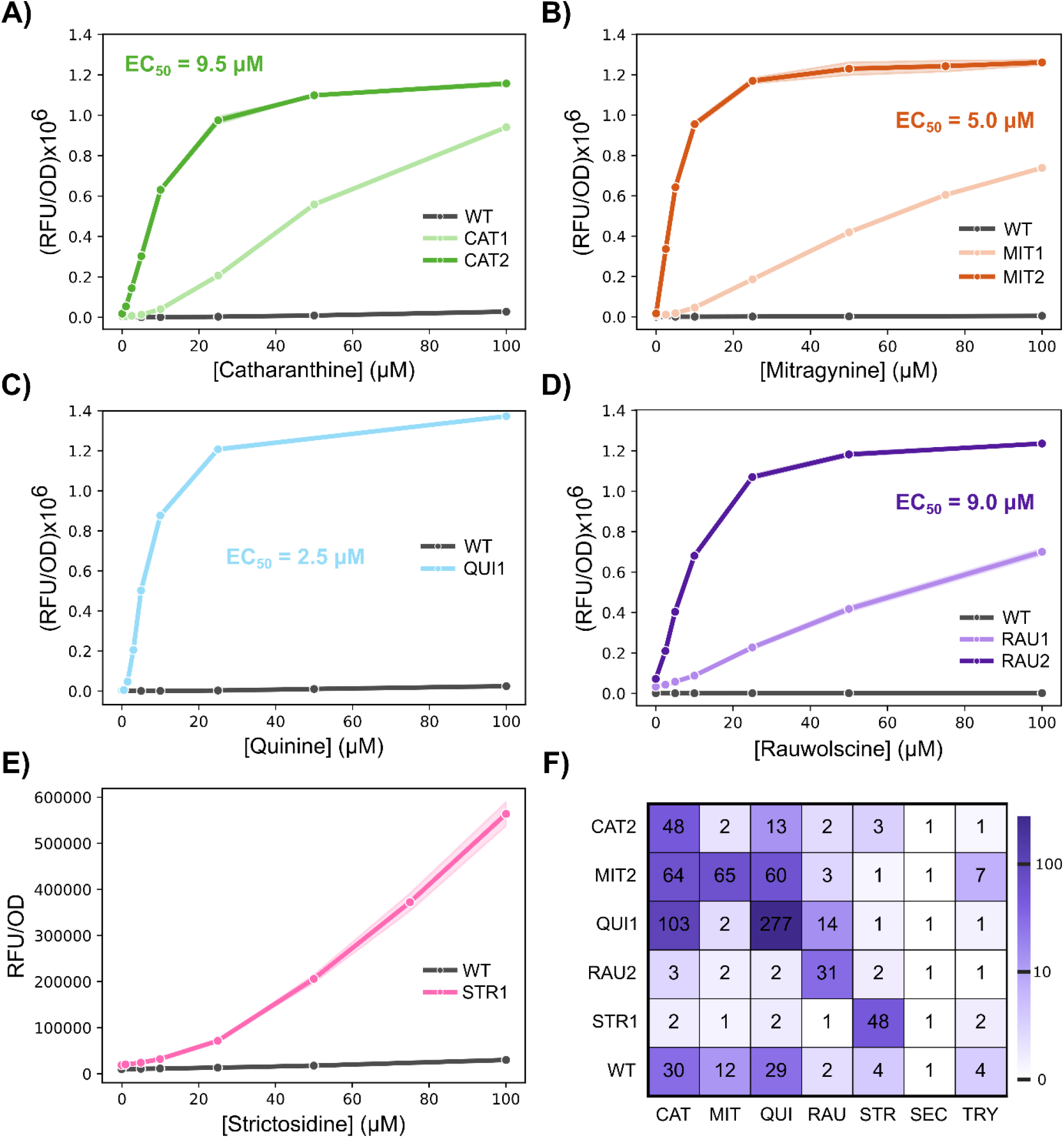
RamR directed evolution campaign for MIA biosensing. **A-E)** Dose-response curves for the different generations of RamR variants for the five different MIAs. Measurements for each condition represent the average of three biological replicates. Shaded regions around trend-line represent S.D.+/−the mean. **F)** Specificity matrix of the final evolved sensors. Fold-change response is shown for all MIAs and their pathway precursors (SEC, secologanin; TRY, tryptamine) for the CAT2, MIT2, QUI1, RAU2, STR1 and WT sensors. The indicated MIAs were supplemented at 100 μM in all conditions except for TRY that was added at 1 mM to reflect fermentation conditions. Measurements for each condition represent the average of three biological replicates.

**Table 1.** Mutations identified in improved RamR MIA biosensors.

| Variant | Mutations | Library fixed |
| --- | --- | --- |
| CAT1 | T85C I88C Y92A | Yellow |
| CAT2 | M143R | Black |
| MIT1 | K63I L66M M70V | Blue |
| MIT2 | M184T L188I T189C | Green |
| QUI1 | K63R L66H M70C | Blue |
| RAU1 | T85N I88A Y92V | Yellow |
| RAU2 | F142M M143S | Lilac |
| STR1 | L133V C134P S137P | Purple |

To assess specificity, we profiled each evolved biosensor against all five target ligands (**Fig. 2F**). The RAU2 and STR1 biosensors exhibited high specificity for their cognate ligand showing no significant response for any of the other MIAs or pathway precursors. Importantly, for these two sensors the operational range is within the order of magnitude of what can be produced in yeast (Bradley et al., 2024; Holtz, Rago, et al., 2024). On the other hand, while CAT2 responded primarily to catharanthine, it showed minor cross-reactivity with QUI. MIT2 responded similarly to CAT, MIT, and QUI, and QUI1 exhibited dual responsiveness to QUI and CAT. Importantly, in practical metabolic engineering scenarios, these MIAs would likely not coexist in the same host cell, as each would be produced in separate pathway contexts. Testing individual biosynthetic intermediates would provide a finer specificity map, but this remains limited by the lack of commercial availability of many of these compounds. Notably, the promiscuous sensors identified here may serve as promising scaffolds for future biosensor engineering efforts targeting related MIAs. The absence of shared mutations among these variants underscores the presence of multiple distinct evolutionary paths toward sensitive RamR-based MIA recognition.

Additionally, RAU2 was evaluated against several yohimbane and heteroyohimbane MIAs: tetrahydroalstonine (THA), yohimbine (YOH), corynanthine (COR), and ajmalicine (AJM), which are known side products of the *Rauvolfia tetraphyla* yohimbine synthase (RteYOS) when expressed in yeast for rauwolscine biosynthesis (Bradley et al., 2024). Remarkably, RAU2 exhibited high enantiospecificity: YOH, COR, and AJM did not yield a significant response, and only THA induced a modest signal at 50 µM of 4.8-fold induction (**Supp. Fig. 3**). This high selectivity was achieved without explicit counterselection during evolution, highlighting RAU2’s potential as a tool to guide future engineering of RteYOS toward improved regio- and enantioselectivity. Such efforts could help reduce shunt product formation in yeast which is a key bottleneck in MIA pathway optimization [7].

### Structure-function analysis of evolved RamR biosensors

To elucidate the structure-function relationships in the engineered RamR biosensors, we integrated crystallographic and computational strategies. We resolved high-resolution crystal structures of CAT2-apo (2.4 Å, PDB ID: 36CD) and QUI1-holo (1.8 Å, PDB ID: 36BZ). For MIT2, STR1 and RAU2, we generated structural models using computational approaches (Boltz-2 co-folding and AutoDock Vina docking). STR1 required modeling due to limited ligand availability, while MIT2 and RAU2 resisted co-crystallization despite extensive efforts.

Backbone responses to evolution differed markedly from those observed in the WT crystal structure (PDB 3VVX): CAT2 shows substantial apo-state remodelling (>4.5 Å dimer C-α RMSD) while QUI1 preserves the wild-type backbone (<2 Å), achieving recognition through sidechain rearrangement alone. Mutations cluster predominantly in the ligand-binding domain helices α4, α5, α8, α9, and α11, leaving other secondary structures untouched (**Fig. 3A, Supp. Fig. 4**). A more detailed analysis of the mutational effects in each variant is provided in the **Supp. Text**.

**Figure 3:**
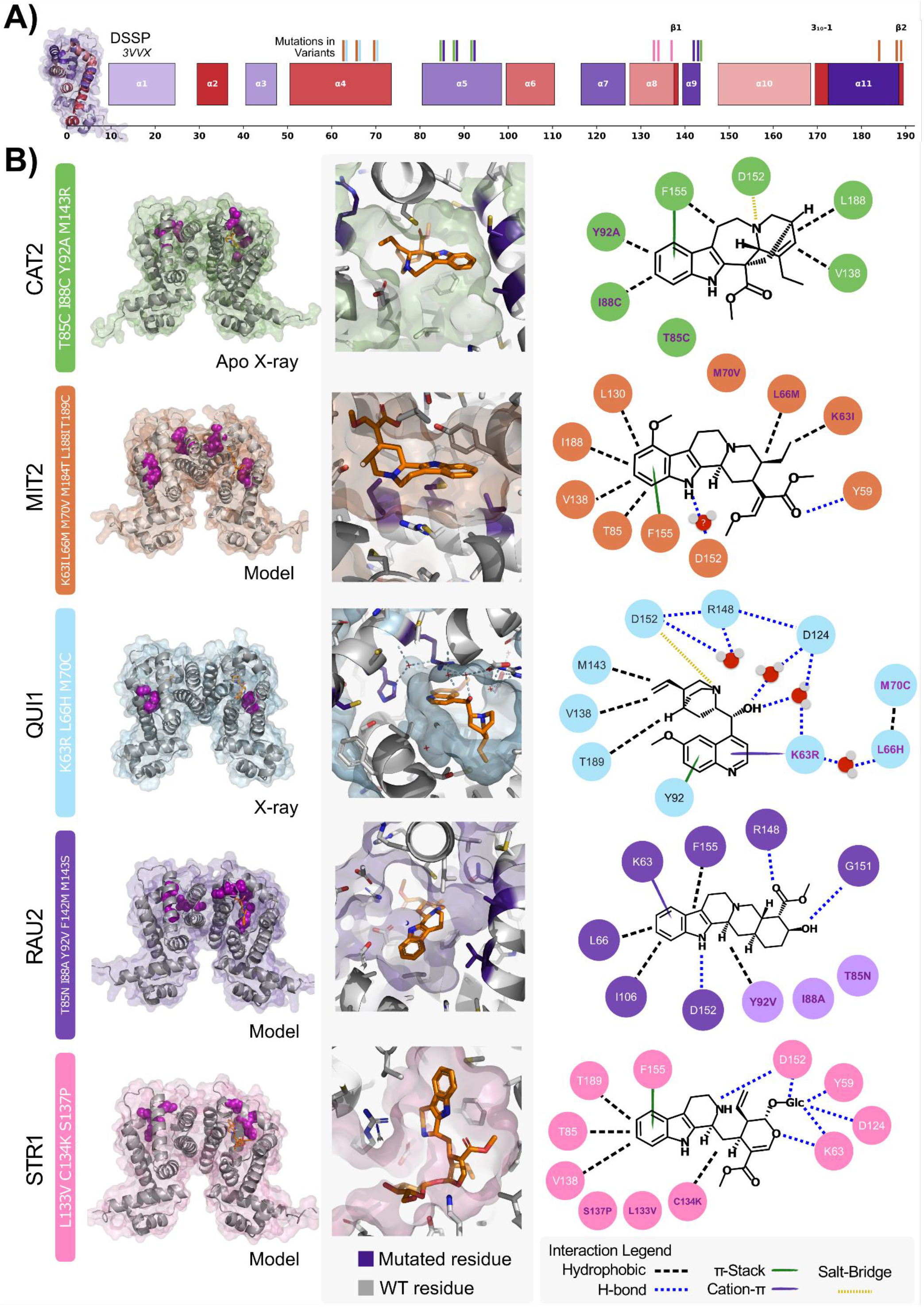
Structure-function analysis of evolved RamR variants. **A**) Annotated secondary structure elements showing the location of accumulated mutations in *α* -helix 4,5,8,9,11. The secondary structure elements *α*_1−3_ form the DNA binding domain. **B)** Panel showing the overall structure (X-ray or modelled) of each final MIA biosensor with mutations accumulated highlighted in purple and the ligand pose highlighted in orange (left). The middle panel shows the zoomed-in ligand binding pose with mutations highlighted in purple. The right-most panel shows the immediate interaction with the ligand, color-coded by interaction type (see legend). For CAT2 a high-resolution apo-structure was obtained, in which by cross-referencing co-folding (Boltz-2) and docking (Vina) poses we estimated an approximate binding pose. For MIT2, a modelled pose is used for the analysis (co-folding (Boltz-2) and docking (Vina)). For QUI1 a high-resolution holo structure was obtained, showing strong hydrogen bonding networks mediated by structural water. For both RAU2 and STR1 cross-referencing co-folding (Boltz-2) and docking (Vina) poses was used to estimate an approximate binding pose. All residues are indexed in a gapless and M0-Q193 frame, corresponding to the wild-type CDS in 0-based index.

Despite the structural diversity of MIA ligands, binding modes recurrently exploit a shared set of generalist residues (**Fig. 3B**). D152 is present in every variant but switches roles by context: a direct salt bridge to CAT, a hydrogen bond to the tertiary amines of STR and RAU, and a water-bridged contact to QUI via R148 and D124 (**Fig. 3B, Supp. Fig. 5-11**). F155 acts as a near-universal π-partner across QUI, MIT, RAU, and STR, complemented by Y92 as a secondary aromatic surface. K63 (or its evolved substitutions) acts as a cation-π donor: but is replaced by isoleucine in MIT2 to relieve a steric clash (**Supp. Fig. 6**), and upgraded to arginine in QUI1, where the guanidinium is conformationally pre-organized by adjacent C70/H66 packing and a structural water that locks the productive rotamer (**Supp. Fig. 7**). In every pose, the hydrophilic face (D152, R148, K63(R), ordered waters) is opposed by a hydrophobic face composed of V138, L130, L188, T189, M143, and F155, producing an amphiphilic pocket complementary to the amphiphilic MIA ligands themselves.

The serendipitous enantioselectivity of RAU2 observed previously (**Supp. Fig. 3**) is rationalized by a single structural feature. Superposing the tested (hetero)yohimbane MIAs (THA, YOH, COR and AJM) onto the modelled RAU2 pose identifies V138 and the D152 backbone in helix α10 as the principal steric gates: only RAU and THA are accommodated without clashes, consistent with the moderate fold-induction observed for THA (**Supp. Fig. 9-10**). For the C-20 and C-16 epimers, inversion at a single stereocenter places the C-17 hydroxyl or methyl ester directly into these gating residues. Enantioselectivity is therefore a consequence of a tightly fitted pocket toward the rich stereocenter regions opposite the planar indole moiety of the yohimbane analogs.

Finally, STR1 shows an adaptation to glycoside-modified substrates: the C134K/L133V/S137P cluster on helix α8 reshapes the pocket through helix repacking, β-bridge perturbation, and a new lysine reaching toward the sugar hydroxyls (**Supp. Fig 11**). This indicates that STR1 and the broader RamR scaffold are plausible starting points for evolving biosensors against other high-value glycoside-modified compounds.

### RamR functionalization in yeast

Following the development of selective RamR-based MIA biosensors in *E. coli*, we aimed to adapt RamR for use in *S. cerevisiae* as a platform for MIA cell factory screening (**Fig. 4A**). Previous studies have demonstrated that numerous prokaryotic aTFs can be re-engineered to function in yeast [23]. A key feature of this process is the design of a synthetic reporter promoter that enables functional biosensor output in the yeast cell context. For repressor-type aTFs, a common design approach involves placing the aTF’s DNA-binding sequence (operator) immediately downstream of the TATA-box of a strong constitutive yeast promoter. This configuration allows repression through steric hindrance of RNA polymerase recruitment upon aTF binding.

**Figure 4:**
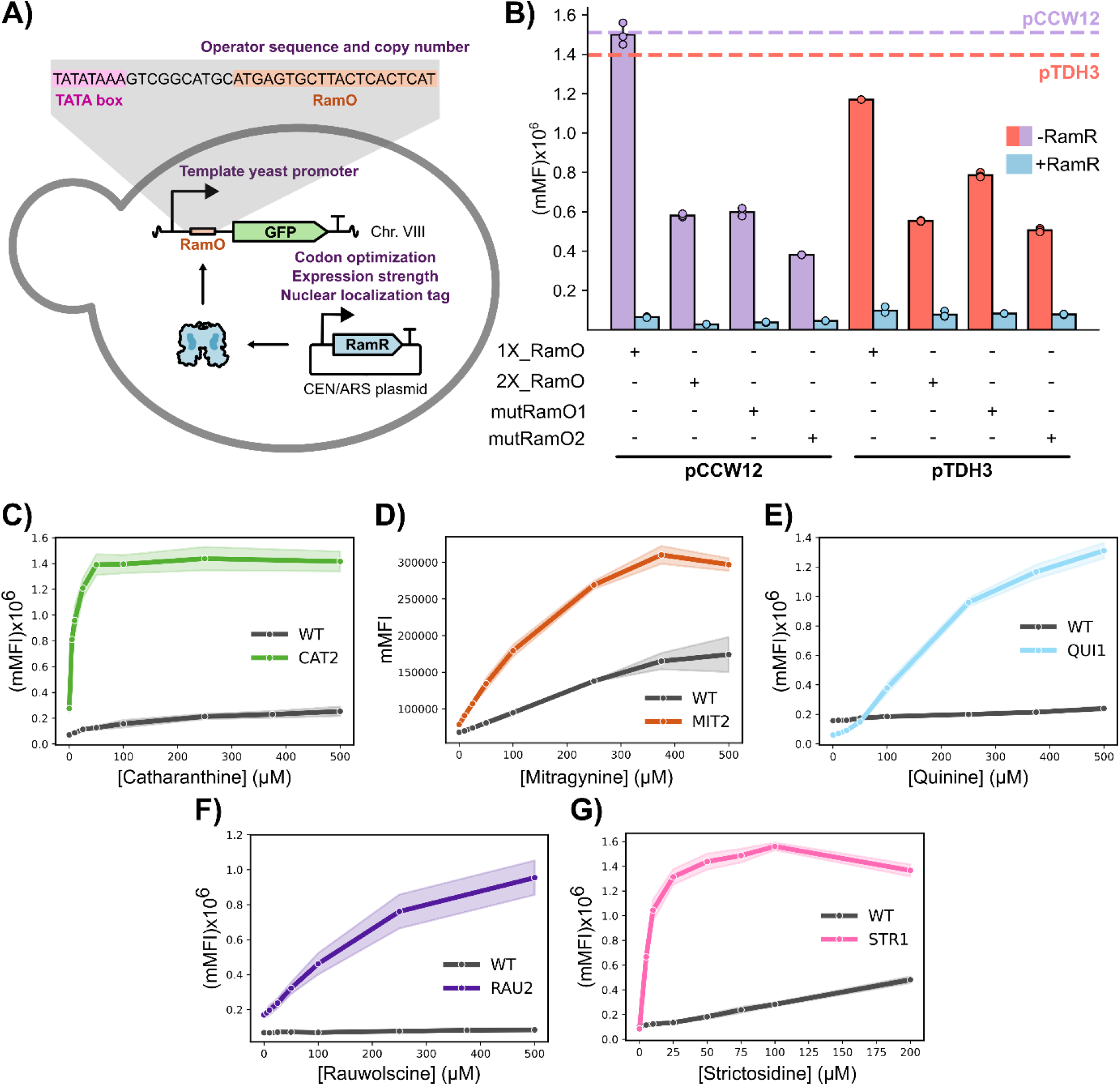
A platform for RamR biosensing in yeast. **A)** Functionalization strategy of RamR in *S. cerevisiae*. Chimeric reporter promoters were engineered by inserting the RamR operator sequence downstream of the TATA box of constitutive yeast promoters. These constructs were cloned upstream of a yeGFP reporter and integrated into the genome, while RamR was expressed from a centromeric plasmid. Optimization strategies for the yeast RamR platform are indicated in purple on the figure. **B)** RamR repression assay for the different reporter promoter variants. Fluorescence levels were determined for each strain after 6 h in the presence or absence of RamR, and the activity of the corresponding template promoters under the same conditions is indicated by dashed lines. Measurements for each condition represent the average of three biological replicates. Error bars represent S.D.+/−the mean.

We selected pTDH3 and pCCW12, two of the strongest constitutive promoters in *S. cerevisiae*, as templates, since both have been previously employed to functionalize TetR-family aTFs in yeast [24,25,25]. Following operator placement strategies from this prior work, we inserted various RamR operator (RamO) sequences into the promoters: one or two copies of the native RamO (designated 1X_RamO and 2X_RamO), as well as two engineered RamO variants (mutRamO1 and mutRamO2) designed as perfect palindromes and shown in *E. coli* to exhibit increased DNA-binding affinity (unpublished data). This yielded eight chimeric promoters, each cloned upstream of yeast-enhanced green fluorescent protein (yeGFP) and integrated into the yeast genome. Non codon optimized RamR was constitutively expressed from pCCW12 on a low-copy CEN/ARS plasmid. We quantified the median yeGFP fluorescence intensity (mMFI) in the presence and absence of RamR to assess repression efficiency. Strong repression of yeGFP was observed for all reporter promoters upon RamR expression, with the pCCW12_1X_RamO variant (strain yCCD12) producing the highest repression, corresponding to a 22.8-fold reduction in fluorescence (**Fig. 4B**). Remarkably, the 1X_RamO configuration consistently outperformed other operator versions in both promoter contexts, which may be attributable to its short sequence (20 bp) and the correspondingly small decrease in basal transcription activity relative to the wild-type promoter (99 %). We next evaluated whether codon optimization would enhance repression in yeast, but found that the optimized sequence yielded slightly reduced repression (**Supp. Fig. 13**). This result supports continued use of the native sequence, which also simplifies cloning by enabling direct transfer of RamR mutants from *E. coli* constructs into yeast.

The final MIA-responsive RamR mutants selected from the *E. coli* screen (CAT2, MIT2, QUI1, RAU2, and STR1) were cloned into the yeast expression plasmid and introduced into the strain containing the optimized reporter construct. Dose-response curves were generated by inducing these biosensor strains with varying concentrations of their respective ligands and measuring fluorescence intensity using flow cytometry (**Fig. 4C-G**). Induction of yeGFP expression with increasing ligand concentration was observed for all sensors, demonstrating that RamR biosensors are fully functional in yeast. The engineered variants retain superior affinity relative to the WT RamR in yeast for their cognate MIA ligand. The apparent sensitivity of the MIT2, QUI1, and RAU2 sensors was lower in yeast compared to *E. coli*. Conversely, the STR1 sensor appeared more sensitive in yeast than in *E. coli*, while CAT2 had a comparable sensitivity in both organisms. Based on the strong pH-dependent differences in biosensor response (**Supp. Fig 14**), we hypothesize that transport processes, including membrane permeation and nuclear access of the ligand, are limiting factors.

Consistently with the results observed in *E. coli*, some of the sensor mutants like QUI1 and CAT2 exhibited a lower background signal than WT RamR indicating a higher DNA-binding affinity in the uninduced state. We reasoned that the biosensor’s transfer function could be altered by the level of repressor expression. By varying the expression strength of these sensors, we found that the transfer function of the biosensor could be modulated, shifting its dynamic and operational range (**Supp. Fig. 15**). The extent of these changes correlated with the DNA-binding affinity of each variant. This approach could be exploited in the future to optimize sensor characteristics for specific strain screening applications.

For metabolic engineering applications, we evaluated genome integration of the RamR expression cassette to improve genetic stability by eliminating plasmid loss and copy-number variability during long-term cultivations. Genome integrated constructs produced more homogeneous fluorescence distributions in flow cytometry, increasing sensor resolution (**Supp. Fig. 16**). Incorporation of a C-terminal SV40 nuclear localization sequence (NLS) further reduced background signal and promoted functional accumulation of RamR in the nucleus (**Supp. Fig. 17**).

Overall, our results establish that RamR mutants evolved in *E. coli* can be functionalized in yeast to create a functional, tunable biosensing platform, with transfer functions adjustable through repressor expression level and nuclear targeting.

### Strictosidine sensing in yeast production strains

Since our last report of engineering *de novo* strictosidine production in yeast [9], strain MIA-CZ-1 was further engineered by swapping the geraniol 8-hydroxylase from *Catharanthus roseus* (CrG8H) with homologs from that enzyme that have been shown to support higher catalytic activity in yeast [26,27] (**Supp. Fig. 18**). Out of the variants tested only TeG8H from *Tabernaemontana elegans* outperformed CrG8H leading to 69 % increase in strictosidine titer reaching 45.9 ± 8.5 mg/L in strain yMHO130. Additionally, switching the carbon source from 2 % glucose to 2 % trehalose with 10 % glycerol also increased production of strictosidine by 44 % reaching 66.2 ± 0.8 mg/L in 3xSC media base (see **Methods**). When grown in YPTreGly media strain yMHO130 produced 74.1 ± 0.3 mg/L strictosidine (**Supp. Fig. 19**).

To evaluate the STR1 sensor as a screening tool for strictosidine production in yeast cell factories, we integrated the STR1-NLS RamR variant together with the optimized reporter construct into the genome of strain yMHO130, generating strain yMHO145. As a negative control, the penultimate enzyme in the strictosidine biosynthetic pathway, CrSLS, was deleted in yMHO145, yielding strain yMHO147, which is incapable of producing strictosidine. Both strains were cultivated in YPTreGly under production conditions, and biosensor fluorescence was measured at 8, 24, 48, and 72 hours (**Supp. Fig. 20**). Additionally, strain yCCD23, expressing only the reporter construct without the repressor, was included to assess the transcriptional capacity of the pCCW12_1X_RamO promoter over time. The largest fluorescence difference between the producing strain (yMHO145) and the non-producing control (yMHO147) was observed at 72 h, with yMHO145 exhibiting a 62 % higher GFP signal. However, this increase was substantially lower than expected considering the order of magnitude of STR titer produced by yMHO145 and dynamic range of the STR1-NLS sensor. To further investigate this discrepancy, we supplemented both yMHO145 and the STR1-NLS non-producer biosensor strain yMHO94 with 100 µM STR. After 6 h, yMHO94 displayed a 20.5-fold induction of yeGFP expression, whereas only a 1.7-fold induction was observed in yMHO145 (**Supp. Fig. 21**). Notably, yMHO145 is a highly engineered strain, harboring 25 overexpressed genes and deletion of 7 native yeast genes. These results suggest that the substantial metabolic burden associated with this extensive engineering impairs the performance of the RamR-based biosensor, leading to reduced sensing capacity in the strictosidine-producing strain. In these conditions, the dynamic output range of the biosensor in the production strain is insufficient to reliably discriminate between enzyme or strain variants.

We first explored an alternative strategy in which the STR1 expressed in *E. coli* was used to detect STR in supernatants from yeast production strains. Multiple conditions were tested, including different supernatant-to-medium ratios, yeast production media, and the presence or absence of buffering. Despite these efforts, STR could not be sensed, and *E. coli* growth was affected (data not shown). This outcome contrasts with previous success using *E. coli* RamR-based sensors to detect sesquiterpenes in yeast supernatants (unpublished data), indicating that the incompatibility likely arises from features specific to the MIA pathway and its metabolites. Based on these observations, we transitioned to a yeast-based sensing approach, expressing the STR1-NLS sensor in a wild-type strain (yMHO94) to enable detection of STR in supernatants derived from production strains (**Fig. 5A**). When inducing yMHO94 with supernatant from the STR producer yMHO130 in buffered YPD at pH 7 with a 1:20 volume ratio, the biosensor signal was >8-fold higher than when using broth from the non-producers MIA-B0 (WT parental strain) or yMHO212 (yMHO130 ΔCrSLS) (**Supp. Fig. 22**).

**Figure 5:**
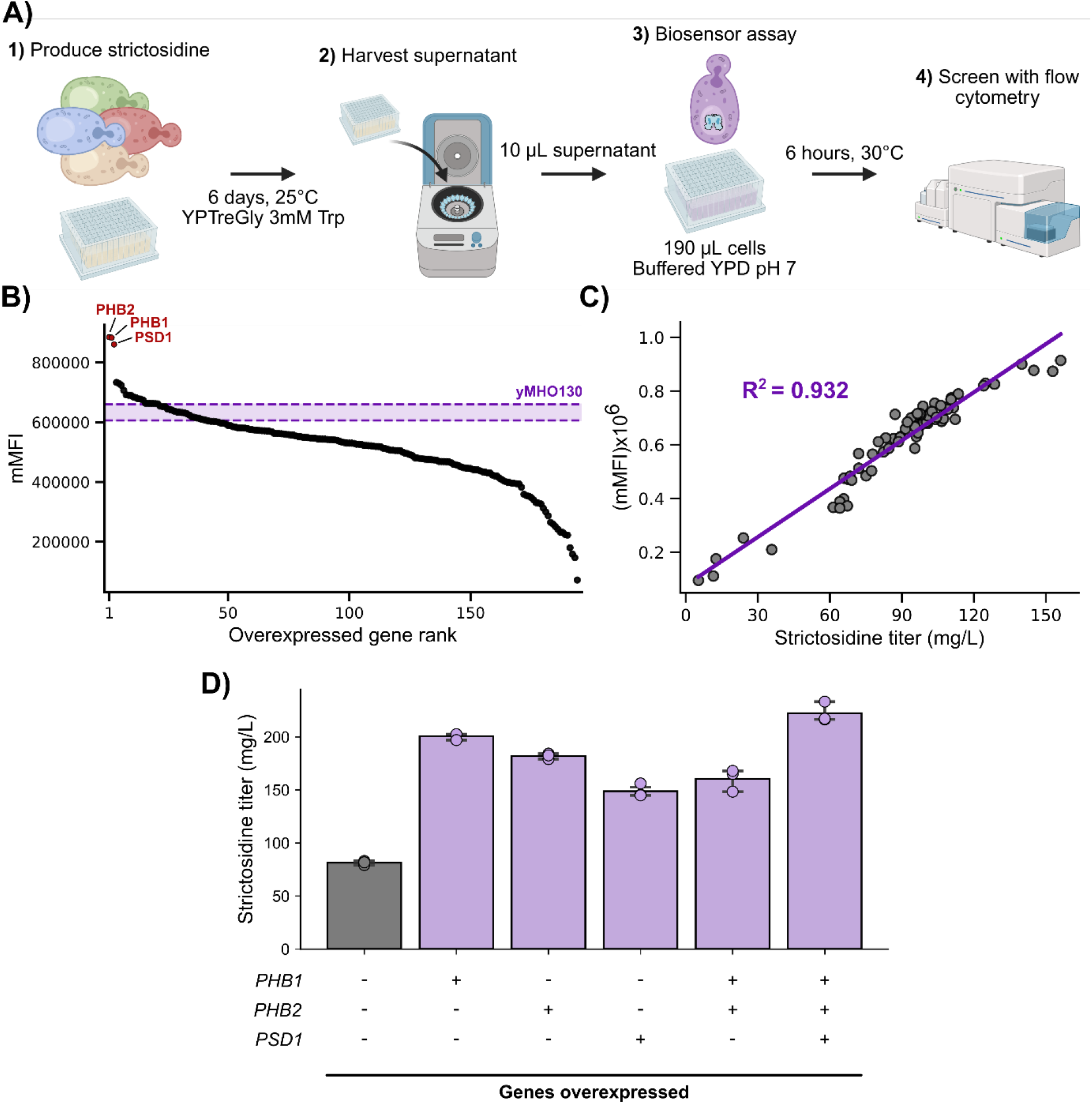
Application of the STR1 sensor for yeast-derived strictosidine biosensing. **A)** Workflow for medium-throughput screening of strictosidine production in engineered yeast cells. **B)** Ranked fluorescence intensity of yMHO94 induced with supernatant from the gene overexpression library integrated in yMHO130. Measurements for each overexpressed gene represent the average of four biological replicates. The control parental strain yMHO130 was grown in each plate and the 95% confidence interval from the fluorescence measured from these samples is shown in purple (n = 30). **C)** Correlation of HPLC-based quantification of strictosidine and yeGFP fluorescence from yMHO94 measured with flow cytometry (mMFI) in a subset of the gene overexpression library integrated in the yMHO130. Data were fitted with a simple linear regression model. For flow cytometry each data point consists of technical triplicates of 30,000 events. **D)** Impact of individual and combined overexpression hits identified from biosensor screen on strictosidine production titer in strain yMHO130 (grey). Measurements for each condition represent the average of three biological replicates. Error bars represent S.D.+/−the mean.

To further demonstrate that such an assay could be used as a semi-throughput screen of STR yeast cell factories, we built a single gene overexpression library in yMHO130 containing 188 candidates (**Supp. Table 5**). These candidates included the STR pathway enzymes, MIA transporters, as well as proteins from *S. cerevisiae* and some of their homologs from *Yarrowia lipolytica* that have been shown to improve production of terpenes and cytochrome P450 functionality in literature. Among others, examples of such proteins include chaperones, cofactor regeneration, and lipid biosynthesis enzymes. These strains each overexpressing one gene from the library were cultivated to produce STR and the resulting supernatant was assayed using yMHO94. A subset of this library (67 samples) was also ran with HPLC to quantify STR titers. The biosensor assay provided a linear correlation between sensor fluorescence and titer (r^2^ = 0.932) demonstrating that it could be used to screen STR strain libraries with high confidence (**Fig. 5B**). Screening the whole library (**Fig. 5C**) enabled identification of three mains hits from *S. cerevisiae*: *PHB1* (YGR132C), *PHB2* (YGR231C) and *PSD1* (YNL169C) which significantly improved STR production. *PHB1, PHB2* and *PSD1* overexpression respectively led STR titers of 200.6 ± 2.5 mg/L (2.7-fold increase), 182.0 ± 2.1 mg/L (2.4-fold increase) and 147.0 ± 6.1 mg/L (2.0-fold increase) (**Fig. 5D**). PHB1/2 are the two subunits of the mitochondrial prohibitin complex, that has chaperone activity, and regulates mitophagy [28]. PSD1 is a phosphatidylserine decarboxylase also located on the mitochondrial inner membrane that plays a role in lipid biosynthesis and mitochondrial fusion and morphology [29]. Although individual overexpression of *PHB1* and *PHB2* improved production, their combined overexpression resulted in a reduced titer compared to each single-gene overexpression (160.3 ± 8.6 mg/L). However, simultaneous overexpression of *PHB1, PHB2*, and *PSD1* resulted in the highest STR production titer observed in this study with 222.2 ± 2.7 mg/L, representing a 3-fold increase in titer over the parental strain. This highlights an example of application of the biosensors developed in this work to identify strain variants with improved MIA production capabilities.

## Discussion

In this study, we used the SELIS directed evolution workflow to evolve RamR to recognize five structurally diverse MIAs. The biosensors developed in this work were highly sensitive (EC_50_ < 10 µM for all but STR1) and exhibited various selectivity profiles with some highly specific variants (RAU2, STR1 and CAT2) and some more promiscuous generalist MIA sensors (MIT2, QUI1) that could be used as starting point to evolve sensors for other MIAs. This work further highlights the versatility of RamR as a generalist aTF, making it a prime candidate for the development of biosensors for bulky, hydrophobic and aromatic compounds, especially alkaloids. The remarkable evolvability of RamR is substantiated by a recent deep mutational scan which investigates the effects of substitutions on DNA binding [30]. It indicates that the mutational landscape for DNA binding and the landscape for ligand recognition overlap substantially at the positions we evolved (**Supp. Fig. 12**). This concordance supports a picture of RamR as a genuinely plastic allosteric scaffold, where ligand-pocket remodelling and DNA-binding competence can be co-optimized rather than traded off. To our knowledge, our work constitutes a seminal report of biosensor development for the highly pharmaceutically relevant MIA family, with the exception of a previous TtgR-based quinine biosensor [31].

While RamR has been evolved to recognize a wide variety of ligands [15,19,20], so far the applications of such biosensors have been limited to *E. coli*. Metabolic engineering efforts for production of plant alkaloids including MIAs, BIAs and AAs would usually be carried in yeast as these pathways contain many ER-anchored cytochrome P450s and require the eukaryotic cell compartmentalization for specific enzyme activities [2,32,33]. By leveraging extensive literature on the functionalization of bacterial transcription factors in yeast [23,34–36], we engineered chimeric constitutive yeast promoters that are repressible by RamR, achieving a dynamic range exceeding 22-fold. The engineered MIA biosensors were functional in yeast and displayed sensitivity profiles that differed from those observed in *E. coli*, with the STR1 sensor exhibiting increased sensitivity, CAT2 remained unaffected, while QUI1, MIT2, and RAU2 showed reduced responsiveness. These differences may arise from variations in small molecule-specific transport across cellular membranes or from host-specific differences in the biosensor context as it has been reported previously [25,36].

Refactoring the engineered STR1 biosensor into STR cell factories enabled *in vivo* sensing of production, but reduced the dynamic output range by >10-fold relative to a reference host without refactored MIA biosynthesis. This attenuation likely reflects the extensive metabolic and cellular engineering of the MIA production strains. Future work should determine whether biosensor performance can be restored through redesign of the sensing circuit, for example implementing signal-amplifier architectures based on activator-driven transcriptional cascade modules [37]. Alternatively, growth-coupled biosensing strategies or selection-based outputs (e.g. antibiotic resistance) may provide a more robust performance under high metabolic load in engineered production strains [38,39].

To mitigate that issue, we designed an assay to detect STR in supernatant from production strains using a separate biosensor strain. Such approach of separating production and sensor strains (prokaryotic or eukaryotic for each type) has been applied previously to sense production of fatty acids [40], D-glucaric acid [41], triacetic acid lactone, naringenin and L-DOPA [42]. Using the STR1-NLS sensor expressed in a WT yeast background enabled robust STR biosensing with a wide dynamic range of GFP signal (>20-fold) and a good linear correlation between metabolite titer and sensor output.

We used this assay to screen the production performance of a single gene overexpression library containing 188 candidates. The top hits *PHB1/2* and *PSD1* led to >2-fold increase in STR titer. Combining these hits further resulted in >220 mg/L STR, a 3-fold improvement over the parental strain. These genes are all related to mitochondrial function and phosphatidylethanolamine (PE) metabolism, which were not previously known to be important in MIA production in yeast cell factories. PSD1 is responsible for the production of PE in mitochondria that is then distributed to other organelles, including the ER [43]. Increasing PE content in membrane is known to increase membrane fluidity and curvature [44], potentially supporting protein insertion, folding and complex formation of heterologous cytochrome P450s and their redox partners [45,46]. The production increase observed upon *PSD1* overexpression may act synergistically with INO2-driven ER expansion in the yMHO130 strain, yielding an enlarged ER that is also optimized for P450 catalysis through increased PE content. The prohibitin complex (PBH1/2) is involved in stabilizing the mitochondrial inner membrane and its proteins, acting as a scaffold that maintains membrane integrity and lipid microdomains [47,48]. This stabilization promotes efficient PE synthesis and trafficking to the ER [48] which could relate to P450 catalysis as described earlier. Additionally, PHB1/2 support mitochondrial function, helping maintain redox balance and cofactor availability, which could also boost pathway flux [49]. To our knowledge, there are no prior reports describing the overexpression of mitochondrial prohibitins (*PHB1* or *PHB2*) or *PSD1* as deliberate metabolic engineering strategies to increase heterologous product titers in *S. cerevisiae*. While *PHB1* has been shown to improve cytochrome P450-dependent production when co-expressed with the plant membrane steroid binding protein AtMSBP1 in yeast [50], such a scaffold is not present in our strains, and the effects of *PHB1* overexpression alone on production have not been previously reported.

Overall, our biosensor-based screening approach enabled the identification of non-obvious engineering targets that enhance MIA production in yeast. This study provides versatile MIA biosensors for metabolic engineering and pathway discovery efforts, directly supporting the development of microbial platforms for stable and scalable production of essential MIA medicines and novel therapies.

## Conclusions

In summary, this work establishes the first biosensor toolkit for high-throughput screening of MIA production in yeast cell factories, overcoming the throughput limitations of conventional analytical chemistry. Coupling engineered RamR-based biosensors with strictosidine production screening enabled the identification of previously unexplored engineering targets linked to mitochondrial function and phosphatidylethanolamine metabolism. Together with rational metabolic engineering and cultivation condition optimization, this biosensor-guided workflow resulted in an overall ∼8-fold increase in strictosidine titer over the course of this work, ultimately exceeding 220 mg/L. Overall, this study establishes a biosensor-enabled engineering framework that could help bring yeast-based MIA production closer to industrially scalable and commercially viable biomanufacturing.

## Materials and Methods

### Strains, plasmids and media

Routine cloning and directed evolution experiments were conducted in *Escherichia coli* DH10B (New England Biolabs), which additionally served as the host for functional characterization of all biosensor constructs. Recombinant protein production of the RamR variants was performed in *E. coli* BL21(DE3) (New England Biolabs). Bacterial cultures used for cloning procedures, fluorescence measurements, orthogonality testing, and evolutionary screening were grown in LB Miller medium (tryptone 10 g/L, yeast extract 5 g/L, NaCl 10 g/L). Solid media consisted of LB agar containing 1.5% (w/v) agar and the respective antibiotics was used during cloning and evolution workflows.

Plasmids described in this work were constructed using NEBuilder HiFi assembly (New England Biolabs) or using the Yeast MoClo (modular cloning) toolkit as described in the original publication [51]. When relevant, antibiotics were used for selection at routine concentrations for *E. coli*: 100 μg.ml^−1^ ampicillin, 50 μg.ml^−1^ kanamycin or 25 μg.ml^−1^ chloramphenicol. Synthetic gene fragments and primers were ordered from IDT (Integrated DNA Technologies). The full list of synthetic gene fragments and plasmids used in this study are respectively available in **Supp. Table 1** and **Supp. Table 2**.

All *S. cerevisiae* strains described in this work are based on CEN.PK2-1C (full list in **Supp. Table 3**). For routine propagation and transformation purposes, yeast strains were grown at 30 °C in Yeast Peptone Dextrose medium (YPD; yeast extract 10 g/L, peptone 20 g/L and glucose 20 g/L). Biosensor characterization was performed in Synthetic Complete medium at pH 7.0 (SC; yeast nitrogen base 6.7 g/L, yeast synthetic drop-out medium supplement without histidine 1.92 g/L, histidine 0.072 g/L and glucose 20 g/L). Yeast genome engineering was carried using the EasyClone-MarkerFree system [52]. Integrative plasmids were linearized by digestion with NotI (New England Biolabs) prior to transformation. Yeast cells were transformed using the conventional lithium acetate chemical transformation procedure [53].

### Chemicals

Ligands used in this study were: mitragynine (Biosynth Ltd, FM26024), quinine (Sigma-Aldrich, 22620), rauwolscine (Sigma-Aldrich, TA9H97BAEB7C), catharanthine (Sigma-Aldrich, PHL80380) and strictosidine (Phytoconsult, discontinued). All ligands had a purity of at least 97 % and stocks were prepared by dissolving the ligand in pure DMSO at a concentration of 25 mM.

### RamR library design and construction

The WT RamR structure (PDB 3VVX) was used to dock the ligands of interest using Autodock Vina [54], analyze tunnels with CAVER (Jurcik et al., 2018), compute residue mutability scores with Hotspot Wizard (Sumbalova et al., 2018). Five SSM libraries came from previous RamR directed evolution work [15]. The other four were designed by selecting residues in close proximity from the docked ligands, present in the protein tunnel bottlenecks and showing good mutability score (**Supp. Fig. 2**). Each final library contained 3 neighbor residues that were randomized using NNS codons using Q5 polymerase (New England Biolabs). Mutagenic primers for library construction are available in **Supp. Table 4**. Error-prone PCR (epPCR) was carried out using the GeneMorph II mutagenesis kit (Agilent), according to the manufacturer’s instructions aiming for aiming for 0-4.5 mutations/kb in the entire RamR CDS. Libraries were assembled into pReg-RamR using NEBuilder HiFi assembly (New England Biolabs) following the manufacturer’s instructions and electroporated in *E. coli* DH10B bearing pSELIS-RamR. Transformation efficiency always exceeded 10^6^ for each round of selection, indicating several fold coverage of the library. Transformed cells were grown in LB medium overnight at 37 °C with ampicilin and chloramphenicol.

### RamR directed evolution in *E. coli*

The SELIS directed evolution workflow was carried as described previously [15]. 20 µL of pooled library liquid cultures was used to inoculate 5 mL of LB containing ampicillin, chloramphenicol, 100 μg.ml^−1^ zeocin (Thermo Fisher, R25001) and 1 mM tryptamine for counterselection. This selection culture was grown for approximately 7 h at 37 °C until reaching an optical density of OD_600_ = 0.5. Following a 1:2000 dilution in fresh LB from this culture, 100 µL were plated on LB agar plates containing ampicillin, chloramphenicol and the target MIA at different concentrations below the EC_50_ of the parental sensor. Plates were incubated overnight at 37 °C. The next day, the brightest colonies were picked and grown overnight in LB with the appropriate antibiotics in a 96-deep-well plate at 37 °C. Cells harboring pSELIS and pReg plasmids encoding the parental RamR variant was also inoculated from a cryostock.

On the subsequent day, 4 µL from each overnight culture was transferred into two individual wells of a fresh 96-deep-well plate, each containing 176 µL of LB medium with appropriate antibiotics. In parallel, four additional were inoculated in the same way with the parental RamR variant strain. Typically, the upper half of the plate comprised 46 distinct clones, while the corresponding duplicates were positioned directly below them in the lower half. After 2 h of growth at 37 °C and 250 r.p.m., the top half of the 96-well plate was induced with 20 μl LB medium with 1 % v/v DMSO, whereas the bottom half of the plate was induced with 100 μl LB medium containing the target MIA ligand at desired concentration (1 % v/v final DMSO concentration). Cells were incubated at 37 °C with 250 r.p.m. shaking for 4 extra hours. The plate was then centrifugated (3500 × g, 4 °C, 10 min) and the pellets were resuspended in 200 µL PBS. One hundred microliters of the cell resuspension for each condition was transferred to a 96-well microtiter plate, from which the GFP fluorescence (excitation, 485 nm; emission, 509 nm; gain 70) and absorbance (600 nm) were measured using a BioTek Synergy H1 plate reader (Agilent). Fluorescence was normalized by the OD_600_ and the hits showing improved sensitivity compared to the parental RamR variant were selected for sequencing and further dose-response characterization.

Dose response of RamR variants were carried out in the same way as the on/off assay described in the previous paragraph. Tested variants were induced with varying MIA ligand concentration after a 2 h growth period keeping the final DMSO content at 1 % for all the assays. After 4 h of growth with the ligands, the normalized fluorescence signal of the sensor was determined.

### Protein purification

The RamR variants were cloned into pET28a(+) vectors and transformed into BL21 DE3 strains for expression. 1 L cultures (LB with 50ug/mL Kanamycin) were inoculated 1:1000 with an overnight preculture and grown at 37°C to an OD_600_ of ∼0.4. IPTG was added to a concentration of 0.1mM for induction and incubated at 16°C for 20 h. The cultures were pelleted (20 min at 4000 g, 4°C) and resuspended in 40 mL lysis buffer (300 mM NaCl, 50 mM Na_2_HPO_4_ pH 8.0, 30 mM imidazole, 15 mM β-mercaptoethanol). In the case of the QUI1 variant, both lysis, elution and storage buffer were supplemented with 1 mM quinine. Cells were lysed by sonication (60 % amplitude, 5 s OFF and 1 s ON, 150x cycles, 2.5min on time). The resulting lysate was spun down (45 min at 15,000 g, 4°C) and the pellet was discarded.

The supernatant was loaded onto a 5 mL HisTrap column, washed with lysis buffer (10 CV), and eluted in elution buffer (30mL). Elution buffer had the same composition as the lysis buffer except the imidazole concentration was 250 mM. The fractions were analyzed by SDS-PAGE. Clean elution fractions were gathered and 1 M DTT was added to final concentration of 3mM. ∼200 U of TEV protease was added to the solution and dialyzed against dialysis buffer overnight at 4°C

His-tag cleavage by TEV was confirmed by SDS-PAGE. The protein was concentrated down to ∼5mL in an Amicon Ultra Centrifugal Filter (10kDa MWCO, Millipore) and stored in 200 mM NaCl, 20 mM Tris-HCl pH 8.0, 3 mM DTT. The sample was gel filtered using a HiLoad 16/600 Superdex 75 pg size exclusion column (Cytiva) equilibrated with the storage buffer, and fractions from the expected peak were confirmed for purity by SDS-PAGE.

For the QUI1 variant complexed with quinine, clean fractions were pooled and concentrated to ∼15 mg/mL. The resulting sample was flash frozen in liquid nitrogen and stored at -80°C.

For the MIT2 variant complexed with mitragynine, clean fractions were pooled and concentrated to ∼1 mg/mL. 50mM mitragynine solution in anhydrous DMSO was added to the concentrated sample to a final concentration of 0.5 mM MIT (10 X molar excess to protein) and incubated overnight. The protein was further concentrated to ∼10 mg/mL, flash frozen in liquid nitrogen and stored at -80°C.

For the CAT2 variant complexed with catharanthine, clean fractions were pooled and concentrated to ∼1 mg/mL. The solution was mixed with an equal volume of 200 mM NaCl, 20 mM Tris-HCl pH 8.0, 200 mM sodium cacodylate pH 8.0, and 3 mM DTT. Then 50 mM catharanthine in DMSO was added to this solution to a final concentration of 0.5 mM CAT (10 X molar excess to protein) and incubated overnight. The protein was further concentrated to ∼5 mg/mL, flash frozen in liquid nitrogen and stored at -80°C.

### X-ray crystallography

To identify crystallization conditions of RamR variants, the purified protein samples were directly used in sparse matrix screening using Phoenix Robotic System (Art Robinson). Initial crystal hits generally appear after incubating the screening plates at 4°C for 1-3 days. Single crystal formation was further optimized by manually setting sitting-drop vapor diffusion experiments, varying pH and precipitant concentrations. Under similar workflow, QUI1-QUI complex formed clusters of cubic crystals in the 2:1 mixture of protein solution with 0.1M sodium citrate (pH 5.5), 0.2 M potassium iodide, and 30 % (w/v) PEG4000. CAT2-CAT complex formed cubic crystals in a 2:1 mixture of protein solution with 5 % PEG1000, 0.1 M sodium phosphate-citrate (pH 5.2), and 20 % ethanol. Individual single crystals were harvested and flash-frozen in liquid nitrogen after brief incubation with a reservoir solution supplemented with 30 % (v/v) glycerol. X-ray diffraction data were collected at beamline NE-CAT 24ID-E (Illinois, US) for QUI1 and at ALS beamline 5.0.3 (California, US) for CAT2. X-ray diffraction data were processed through automated data processing pipeline, xia2 [55]. Phenix was used for phasing by molecular replacement with the published RamR wildtype model (PDB code: 3VVX) used as the search model [56]. The molecular replacement solution was iteratively built and refined using Coot and Phenix refine package [57].The quality of the final refined structures was evaluated by MolProbity [58]. The final statistics for data collection and structure determination are shown in **Supp. Table 6**.

### Structural modelling of RamR

Canonical helix and strand assignments for wild-type RamR were obtained by running DSSP [59] on the apo WT crystal structure (PDB ID: 3VVX). The resulting assignments were used as the reference nomenclature throughout this work, with helices numbered α1-α11 and all residues indexed in a gapless and M0-Q193 frame, corresponding to the wild-type CDS in 0-based index, for consistency with respect to previously published work [15].

Protein-ligand co-folding was performed with Boltz-2 [60] via the Rowan computational platform (Rowan Scientific Corporation, 2025). Each co-folding run used 5 seeds with diffusion guidance, energy minimization, and ligand ranking by confidence score. MSA generation was enabled (use_msa_server: true) along with physically-grounded potentials (use_potentials: true). Predicted complexes were filtered for stereochemical and physicochemical plausibility using PoseBusters [61], and only valid poses were retained for analysis.

Molecular docking was performed with AutoDock Vina [54] via the Rowan platform, using the Vinardo scoring function [62] and an exhaustiveness of 8. Apo receptor structures used as docking targets were taken either from the experimental crystal structures (CAT2) or from Boltz-2 co-folded models (RAU2, STR1). Search boxes of 10 × 10 × 10 Å were centered on the geometric mean of the binding-pocket residues, and pose refinement was enabled (do_pose_refinement: true).

For variants lacking an experimental holo structure (MIT, CAT2, RAU2, STR1) candidate binding poses were obtained by cross-referencing Boltz-2 co-folded ensembles against Vina poses. The poses showing agreement between the two methods, satisfying PoseBusters validity, and consistent with the accumulated mutations were retained for structural interpretation.

Low-energy conformer ensembles for RAU and its structural analogs (yohimbine, corynanthine, ajmalicine, tetrahydroalstonine) were generated using CREST [63] via the Rowan platform. Conformer search was performed with the iMTD-GC metadynamics protocol using GFNn-xTB [64] for the metadynamics sampling, followed by single-point refinement at the g-xTB [65] level of theory. The lowest-energy conformer of each analog was used for superposition onto the modelled RAU2 holo pose (**Supp. Fig 9-10**).

Direct intermolecular interactions: hydrogen bonds, salt bridges, π-stacking, cation-π, hydrophobic contacts, and water bridges were identified using the Protein-Ligand Interaction Profiler (PLIP) [66] on both experimental holo structures and modelled complexes. Default detection thresholds were used throughout on the PLIP webserver.

### Yeast biosensor characterization

Yeast strains carrying the RamR biosensing platform were grown overnight in SC-URA (plasmid based RamR expression) or in YPD (genome integrated RamR) at 30 °C and 250 r.p.m. The following day, the precultures were diluted 1:25 in fresh SC-URA or SC media depending on whether the strain expressed RamR on a plasmid or not. Unless otherwise specified, all biosensor characterization experiments were carried out at pH 7.0. From these cell suspensions, 180 µL was transferred per well in a 96-deep-well plate. The plate was incubated for 2 h at 30 °C and 250 r.p.m before inducing the biosensors with the MIA ligand. Similarly to the biosensor characterization in *E. coli*, 20 µL of MIA ligands in SC media was used as inducer bringing the DMSO concentration to 1 % v/v for any assay. Following additional 6 h of incubation at 30 °C and 250 r.p.m, 50 µL of cells were mixed with 100 µL of PBS in a microtiter plate and fluorescence was analyzed using a NovoCyte Quanteon (Agilent) flow cytometer. For each condition, three biological replicates were analyzed with a threshold of 20,000 events per replicate. The cells were gated for singlets in the exponential phase and the median fluorescence intensity (MFI) for the GFP signal (FITC-H channel) was determined using FlowJo v10.8 Software (BD Life Sciences).

### Strictosidine production in yeast

Strictosidine production strains were inoculated from a plate in 1 mL of YPD in 15 mL preculture tubes and incubated overnight at 30 °C and 250 r.p.m shaking. The next day 50 μL of preculture was used to inoculate 250 μL of production media in a 96-deep well plate. Unless stated otherwise, the production media was YPTreGly (yeast extract 10 g/L, peptone 20 g/L, trehalose 20 g/L and glycerol 100 g/L). All cultivations were run as biological triplicates originating from different colonies. Plates were incubated at 25 °C for 144 h with shaking at 300 r.p.m. Following 144 h, 100 μL of each sample was filtered through a filter plate (PALL, AcroPrep Advance, 0.2-μm Supor membrane for medium/water) by centrifugation at 2200 g for 1.5 min. Samples were stored at −20 °C prior to injection or biosensor assay.

### Strictosidine biosensing from yeast supernatant

Biosensing of strictosidine was carried using strain yMHO94 which was grown overnight in YPD at 30 °C and 250 r.p.m. The following day, the preculture was diluted 1:25 in fresh YPD media buffered at pH 7.0 (potassium phosphate buffer, 100 mM). 190 µL of this cell suspension was added into each well of a 96-deep-well plate which was incubated at 30 °C and 250 r.p.m for 2 h. After this time, the biosensing strain was induced with 10 µL of filtered supernatant from strictosidine production strains (1:20 dilution). Following additional 6 h of incubation at 30 °C and 250 r.p.m, 50 µL of cells were mixed with 100 µL of PBS in a microtiter plate and fluorescence was analyzed using a NovoCyte Quanteon (Agilent) flow cytometer. Flow cytometry data was analyzed the same way as for the yeast biosensor characterization section.

### Overexpression library construction in yMHO130

Each gene from the overexpression library was amplified from genomic DNA of *S. cerevisiae, Y. lipolytica*, or from synthetic gene templates, depending on gene origin (see **Supp. Table 5**). Coding sequences (CDSs) were amplified using Platinum SuperFi II DNA Polymerase (Thermo Fisher) with 10 ng of template DNA. Primers were designed to include 30 bp overhangs homologous to the pTDH3 promoter (5′) and tADH1 terminator (3′).

These overhangs enabled *in vivo* assembly of each CDS with two additional PCR fragments: (i) an UP fragment containing a 500 bp upstream region of site VIII-1 and the pTDH3 promoter, and (ii) a DOWN fragment containing a 500 bp downstream region of site VIII-1 and the tADH1 terminator. This assembly allowed targeted genomic integration of each overexpression construct at the VIII-1 locus using the corresponding gRNA plasmid (pCFB9340) [67].

Transformations were performed in a 96-well plate format. Each well contained 500 ng of gRNA plasmid, 6 µL of pooled, unpurified UP and DOWN PCR fragments, and 10 µL of unpurified CDS PCR product. Following transformation, cells were plated and colonies were picked using QTrays in combination with a QPix 460 colony picking system (Molecular Devices). For each gene in the library, four independent colonies were selected and screened for strictosidine production and biosensingSupp. Fig..

### High-performance liquid chromatography analysis

Strictosidine containing samples were run on a Zorbax Eclipse Plus C18 reverse-phased column (internal diameter 4.6 mm, particle size 3.5 µm, pore size 95 A, length 100 mm, Agilent). The mobile phase consisted of Mili-Q water with 0.1 % formic acid (solvent A) and 100 % acetonitrile (solvent B). The solvent composition was initially A = 98.0 % and B = 2.0 % for 2 min. A linear gradient was run until A = 90.0 % and B = 10.0 % at 5.0 min. A second gradient was run until A = 85.0 % and B = 15.0 % at 8.0 min. At 8.2 min, A was set to 2.0 %, and B to 98.0 %. This condition was kept constant until 9.5 min, after which the initial composition was retrieved (A = 98.0 %, B = 2.0 moa%) and remained unchanged until the end of the run (11.5 min). The column oven temperature was set to 30 °C and the flow rate was set to 1 mL/min with 10 µL of sample injection. Quantification was based on the specific peak area (RT = 6 min) at a detection wavelength of 280 nm.

## Supporting information

Holtz et al, 2026 Supplementary Information

## Acknowledgements

This work was funded by the Novo Nordisk Foundation Copenhagen Bioscience Ph.D. Program grant No. NNF22SA0078231 (awarded to MH) and Novo Nordisk Foundation grant No. NNF20CC0035580. NGM was supported by the Danish Data Science Academy, which is funded by the Novo Nordisk Foundation (grant No. NNF21SA0069429) and VILLUM FONDEN (grant No. 40516). ACAA was funded by the Rubicon research programme which is financed by the Dutch research council (NWO) (file number 019.231EN.007). JAA received funding from the Novo Nordisk Foundation grant No. NNF24OC0095360. The structural work was supported by the National Institutes of Health grant R35GM148356 (Y.J.Z.).

