## Supplementary material for "Engineering Biosensors to Enhance Monoterpene Indole Alkaloid Production in Yeast": Holtz et al, 2026 Supplementary Information

### Contents of Supplementary Information


**Supplementary Figure 1.** Core MIA biosynthesis pathway.

**Supplementary Figure 2.** Library design rationale for RamR directed evolution and residue mapping on WT RamR (PDB 3VVX).

**Supplementary Figure 3.** Response of the RAU2 biosensor induced by different yohimbane and heteroyohimbane MIAs in *E. coli*.

**Supplementary Figure 4.** RamR secondary structure annotation to label helix numbers.

**Supplementary Figure 5.** CAT2 apo variant structure with CAT.

**Supplementary Figure 6.** MIT2 holo modelling to determine residues driving ligand binding.

**Supplementary Figure 7.** QUI1 holo structure implicating structural water in binding.

**Supplementary Figure 8.** RAU2 holo modelling to determine residues driving ligand binding.

**Supplementary Figure 9.** RAU2 holo modelling to determine residues driving enantioselectivity.

**Supplementary Figure 10.** Further RAU2 holo modelling to determine residues driving enantioselectivity

**Supplementary Figure 11.** STR1 modelled holo structure.

**Supplementary Figure 12.** Cross-referencing Growth-based Quantitative Sequencing (GROQ-seq) data with RamR mutations obtained in this work.

**Supplementary Figure 13.** Comparison of the repression efficiency of codon-optimized RamR for *S. cerevisiae* and the native non-codon optimized sequence in yeast.

**Supplementary Figure 14.** Effect of pH on RamR  MIA biosensing in yeast.

**Supplementary Figure 15.** Effect of RamR expression strength on biosensor transfer function for CAT2 (**A**), QUI1 (**B**) and RAU2 (**C**).

**Supplementary Figure 16.** Modulation of RamR biosensing by genomic integration and nuclear localization.

**Supplementary Figure 17.** Comparison of RamR MIA biosensor dose–response in yeast (**A**) and background signal in the absence of ligand (**B**) for plasmid-based versus genome-integrated RamR circuits, with or without an SV40 NLS tag.

**Supplementary Figure 18.** Geraniol hydroxylase (G8H) screen in MIA-CZ-1 for strictosidine production.

**Supplementary Figure 19.** Comparison of production media for STR production by yMHO130.

**Supplementary Figure 20.** Strictosidine biosensing in STR production strain yMHO145 and non-producer CrSLS knockout yMHO147 over time.

**Supplementary Figure 21.** Comparison of genome integrated STR1-NLS biosensor signal without ligand (green) and induced with 100 µM STR (orange) in a WT background yeast strain (yMHO94) and in a strictosidine producer (yMHO144).

**Supplementary Figure 22.** Strictosidine biosensing with yMHO94 (STR1-NLS sensor) induced with supernatant from different yeast strains (1:20 volume ratio).

**Supplementary Table 1.** Sequences of synthetic DNA fragments used in this study.

**Supplementary Table 2.** Plasmids cloned and used in this study.

**Supplementary Table 3.** Yeast strains used in this study

**Supplementary Table 4.** Primers used for RamR library construction.

**Supplementary Table 5.** Overview of genes included in the overexpression library tested in yMHO130 to demonstrate an application of the STR1 sensor.

**Supplementary Table 6.** X-ray crystallography data collection and refinement statistics.

**Supplementary Text**

#### Supplementary Figures

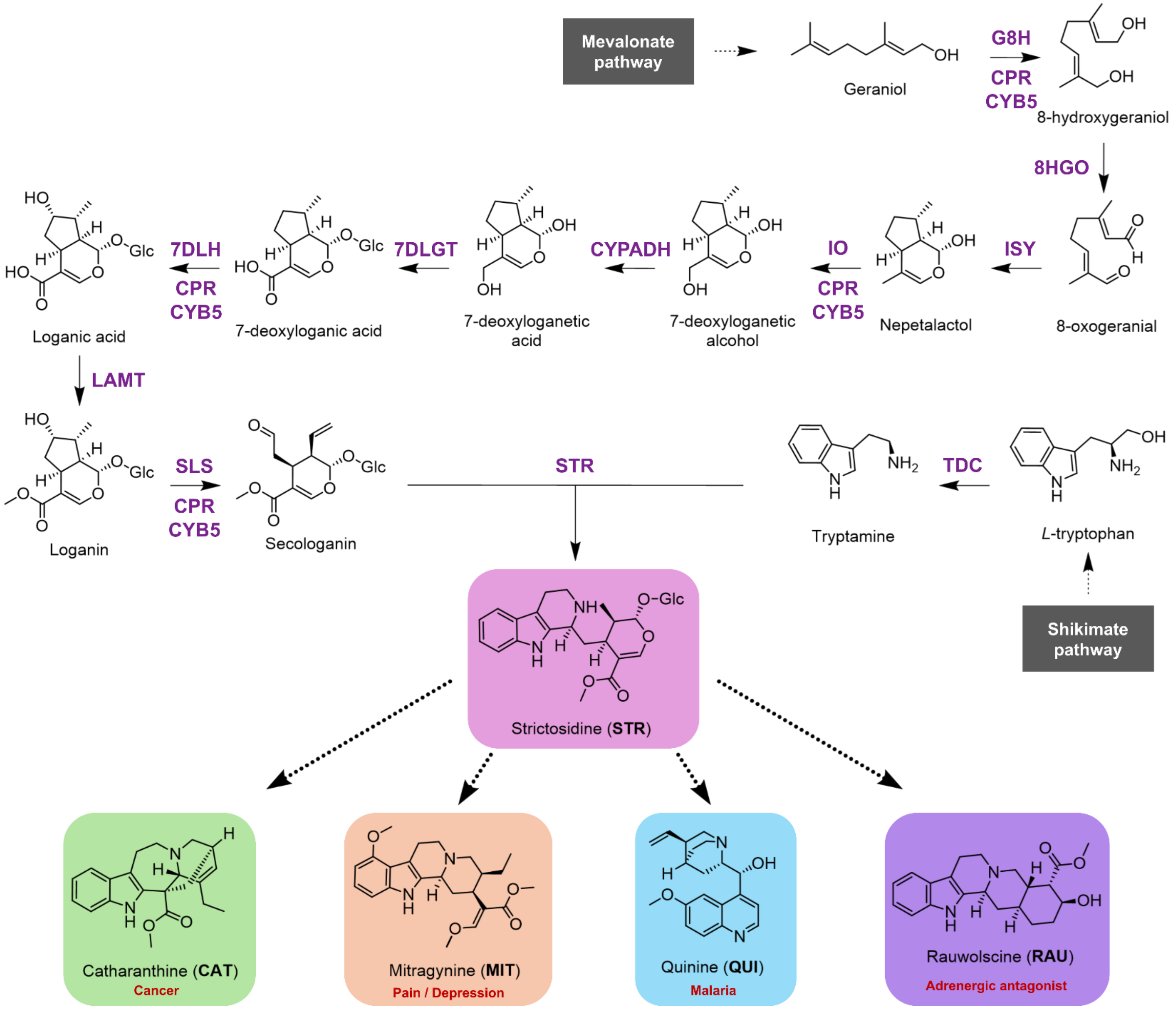


**Supplementary Figure 1.** Core MIA biosynthesis pathway. The central MIA precursor strictosidine gives rise to a wide diversity of >3000 MIA structures through the action of tailoring enzymes. Abbreviations are as follows: IPP, isopentenyl pyrophosphate; DMAPP, dimethylallyl pyrophosphate; GPPS, GPP synthase; FPSN144W, FPP synthase N144W variant; CPR, NADPH-cytochrome P450 reductase; CYB5, cytochrome b5; GES, geraniol synthase; G8H, geraniol 8-hydroxylase; 8HGO, 8-hydroxygeraniol oxidoreductase; ISY, iridoid synthase; IO, iridoid oxidase; CYPADH, alcohol dehydrogenase 2; 7DLGT, 7-deoxyloganetic acid glucosyl transferase; 7DLH, 7-deoxyloganic acid hydroxylase; LAMT, loganic acid O-methyltransferase; TDC, tryptophan decarboxylase; SLS, secologanin synthase; STR, strictosidine synthase.

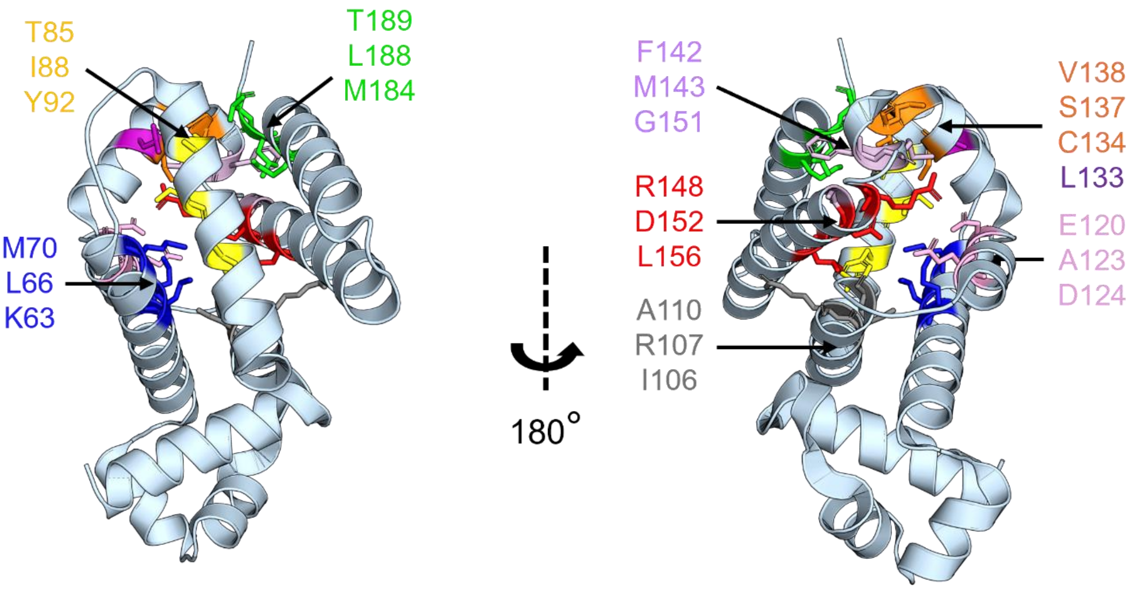


| Library color | Residues targeted | Design rationale | Reference |
|---|---|---|---|
| **BLUE** | K63 L66 M70 | Previously used to successfully engineer BIA biosensors. | (d'Oelsnitz et al., 2022) |
| **YELLOW** | T85 I88 Y92 | Previously used to successfully engineer BIA biosensors. | (d'Oelsnitz et al., 2022) |
| **PINK** | E120 A123 D124 | Previously used to successfully engineer BIA biosensors. | (d'Oelsnitz et al., 2022) |
| **PURPLE** | L133 C134 S137 | Previously used to successfully engineer BIA biosensors. | (d'Oelsnitz et al., 2022) |
| **RED** | R148 D152 L156 | Previously used to successfully engineer BIA biosensors. | (d'Oelsnitz et al., 2022) |
| **SILVER** | I106 R107 A110 | Proximity to docked MIAs. Good mutability score from Hotspot Wizard. | This study. |
| **ORANGE** | C134 S137 V138 | Proximity to docked MIAs. Good mutability score from Hotspot Wizard. V138 was a hotspot in previous epPCR evolution campaigns (unpublished data). C134 is a bottleneck residue in the main RamR tunnel computed by CAVER. | This study. |
| **LILAC** | F142 M143 G151 | Proximity to docked MIAs. Good mutability score from Hotspot Wizard. M143 is a bottleneck residue in the main RamR tunnel computed by CAVER. | This study. |
| **GREEN** | M184 L188 T189 | Proximity to docked MIAs. Good mutability score from Hotspot Wizard. M184 and L188 came out as hits in previous epPCR evolution campaigns (unpublished data). | This study. |
| **BLACK** | Error-prone PCR (random) | Find non-obvious mutations. | This study. |

**Supplementary Figure 2.** Library design rationale for RamR directed evolution and residue mapping on WT RamR (PDB 3VVX). Residues were numbered based on that reference structure and to match previous work on RamR engineering (this experimental structure misses the first Met residue).

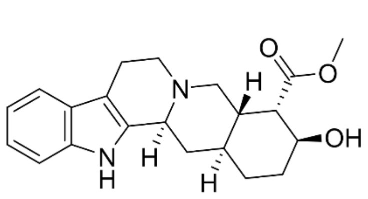

Rauwolscine (**RAU**)

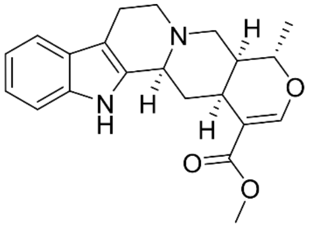

Tetrahydroalstonine (**THA**)

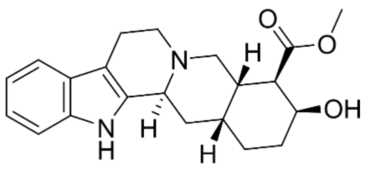

Yohimbine (**YOH**)

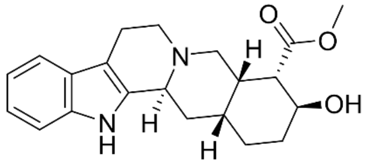

Corynanthine (**COR**)

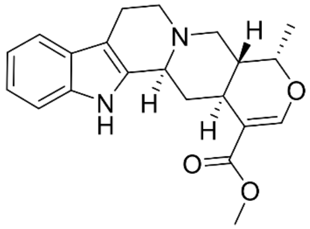

Ajmalicine (**AJM**)

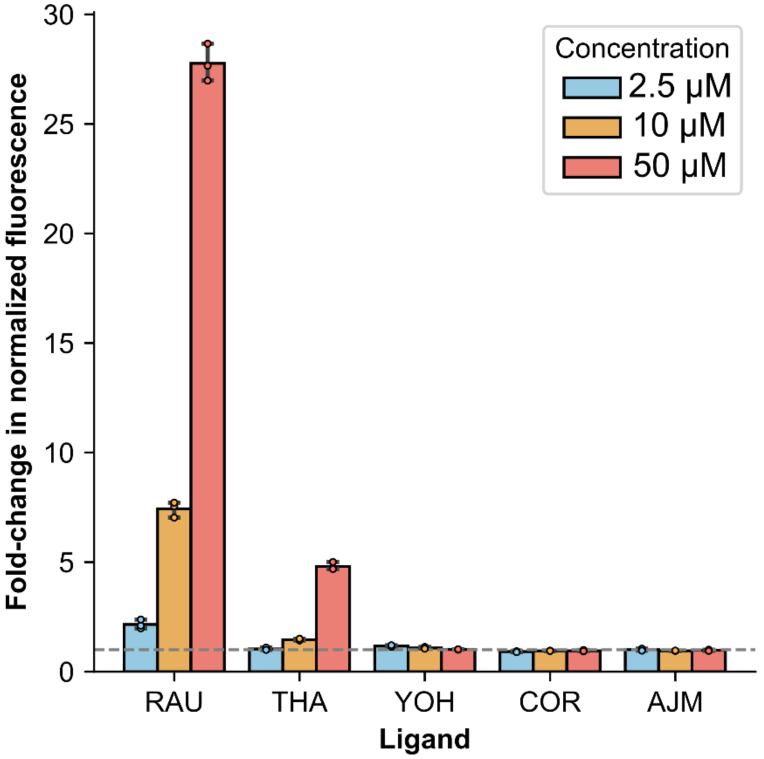


**Supplementary Figure 3.** Response of the RAU2 biosensor induced by different yohimbane and heteroyohimbane MIAs in *E. coli*. Fluorescence was normalized by OD at 600 nm and fold change was calculated by dividing by the normalized fluorescence value for a control without ligand (DMSO 1%). Measurements for each condition represent the average of three biological replicates. Error bars represent S.D.+/−the mean.

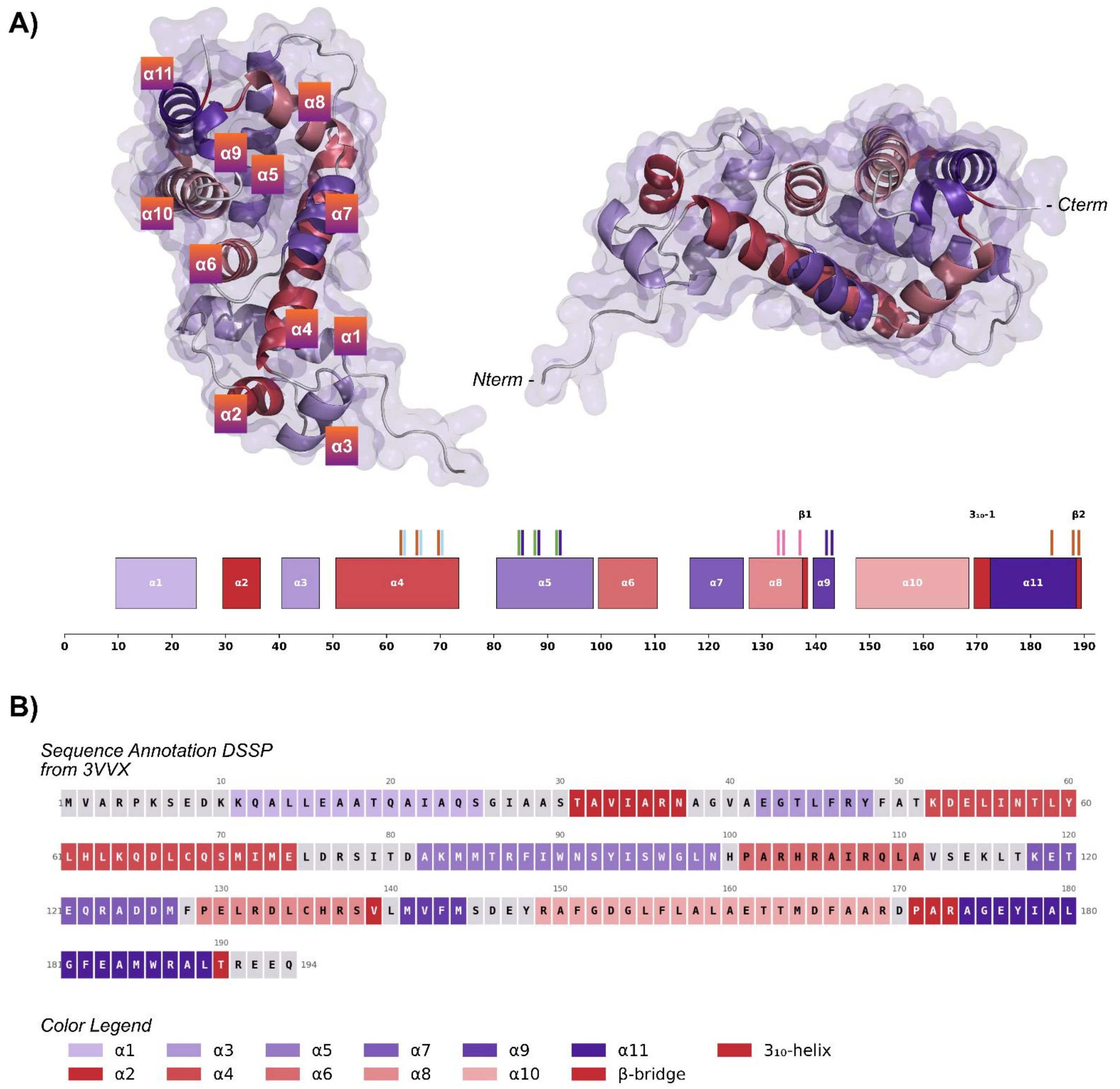


**Supplementary Figure 4**. RamR Secondary Structure Annotation to label Helix Numbers. **A)** DSSP annotated secondary structure overlayed on the apo wild-type RamR (PDB ID: 3VVX) given canonical notation to the secondary structure helices. Notably there exists a single-pair anti-parallel β-sheet annotation between T189-V138 as well as a short 3_10 helix segment. **B)** DSSP annotation overlaid on the wild-type RamR sequence. Grey indicates either loop or coil DSSP codes.

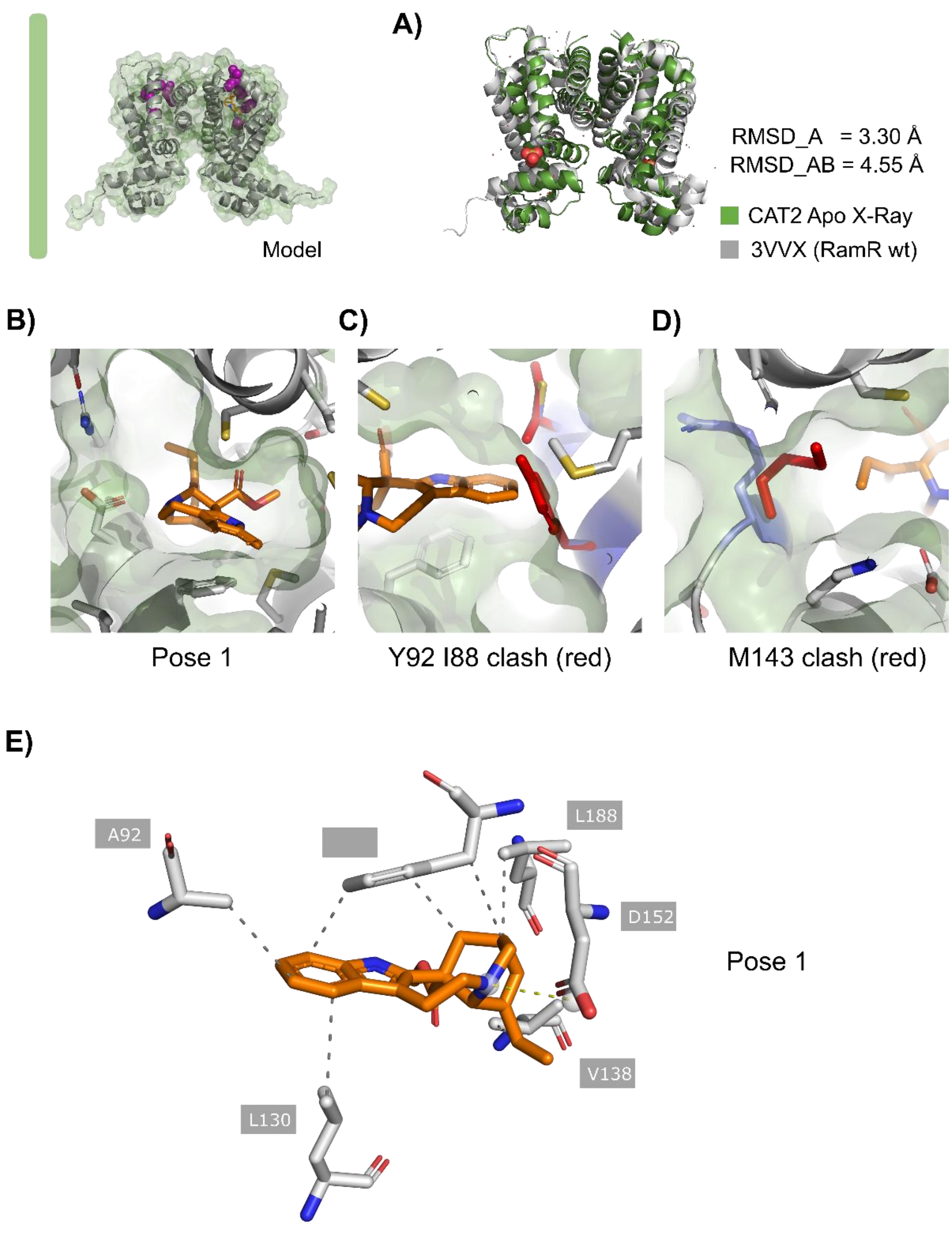


**Supplementary Figure 5**. CAT2 Apo Variant Structure with CAT. **A)** Comparison of C-alpha RMSD between wild-type RamR (3VVX) and CAT2-apo shows that when comparing both chains simultaneously a large conformation change (>4.5 Å) has been induced by mutations alone. Aligning only chain A, rather than the homodimer, still results in a large deviation from the wild-type RamR. **B)** Pose 1, obtained from cross-referencing co-folding (Boltz-2) and docking (Vina) poses, indicates moderate shape complementarity. **C)** Illustrating the wild-type residues as positions 88, 92 (I,Y) in red, reveals their steric incompatibility with the binding pose of CAT, indicating the role of the mutations I88C Y92A in CAT2. **D)** Illustration of the wild-type residue M143 (red) shows a vdW steric clash with CAT, illustrating the function of M143R. The arginine swings out toward the solvent exposed surface, leaving space for the ligand. **E)** Protein-Ligand Interaction Profiler (PLIP) shows key immediate contacts to CAT. A hydrophobic sandwich is formed by L130 and F155, while D152 forms a salt bridge with CAT.

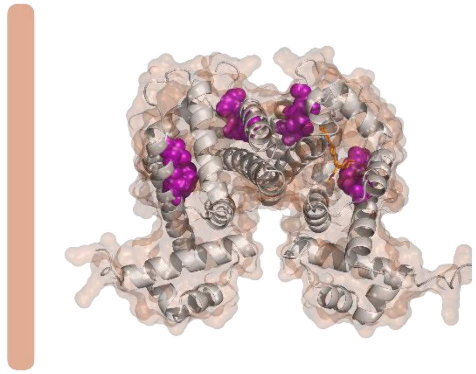

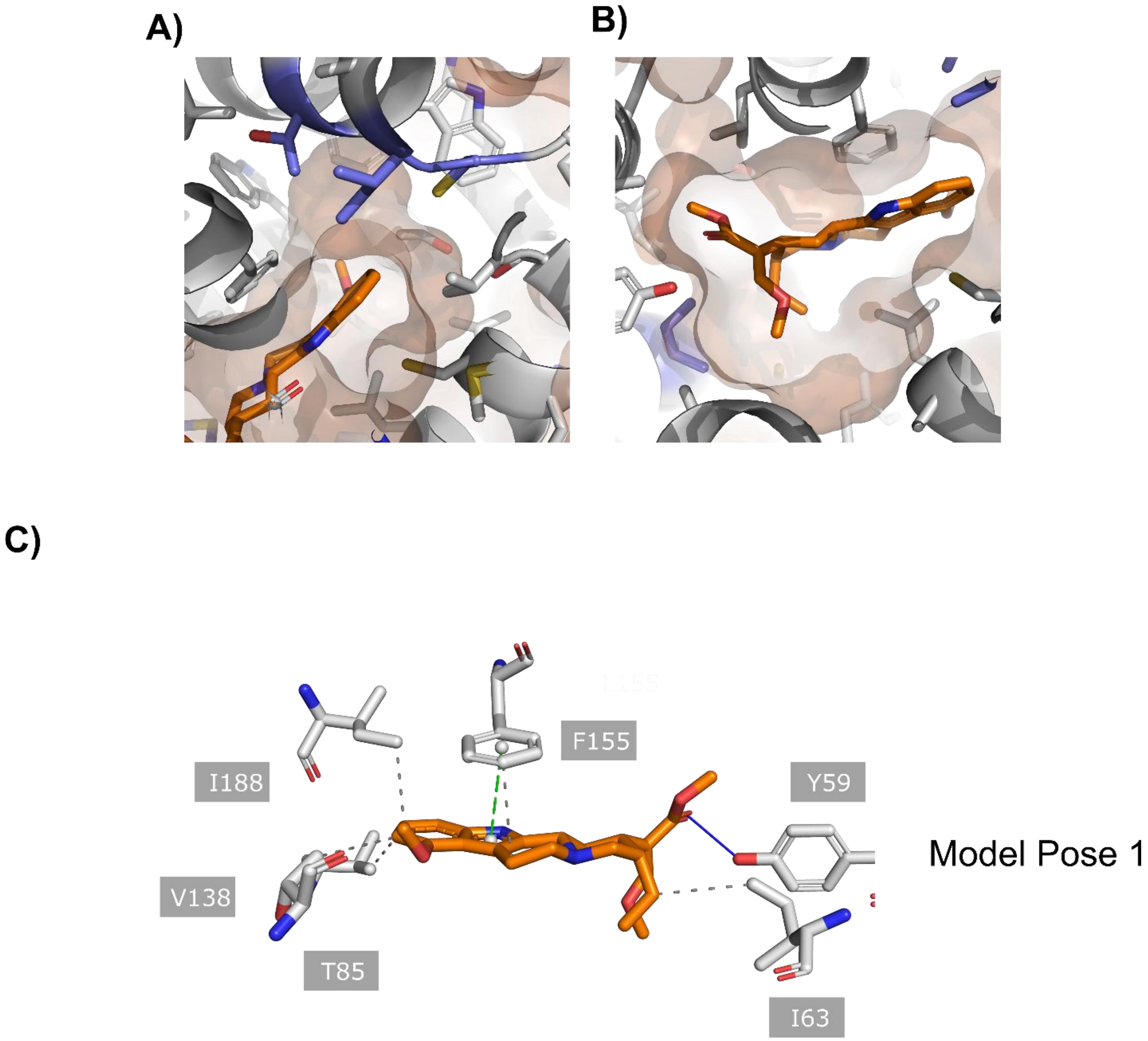
A)
B)
C)
I188
F155
Y59
V138
T85
I63
Model Pose 1

**Supplementary Figure 6.** MIT2 Modelled Holo Structure. **A)** Modelled pose of MIT obtained shows stronger shape complementarity to binding pocket and MIT is adjacent to the mutated clade: K63I L66M M70V. **B)** Rotated view of the pose in A). **C)** PLIP diagram of the modelled alternative pose indicates the possible role of Y59 as a H-bond donor to the carbonyl of MIT, F155 as a (π-π-bond) and various other immediate hydrophobic interactions.

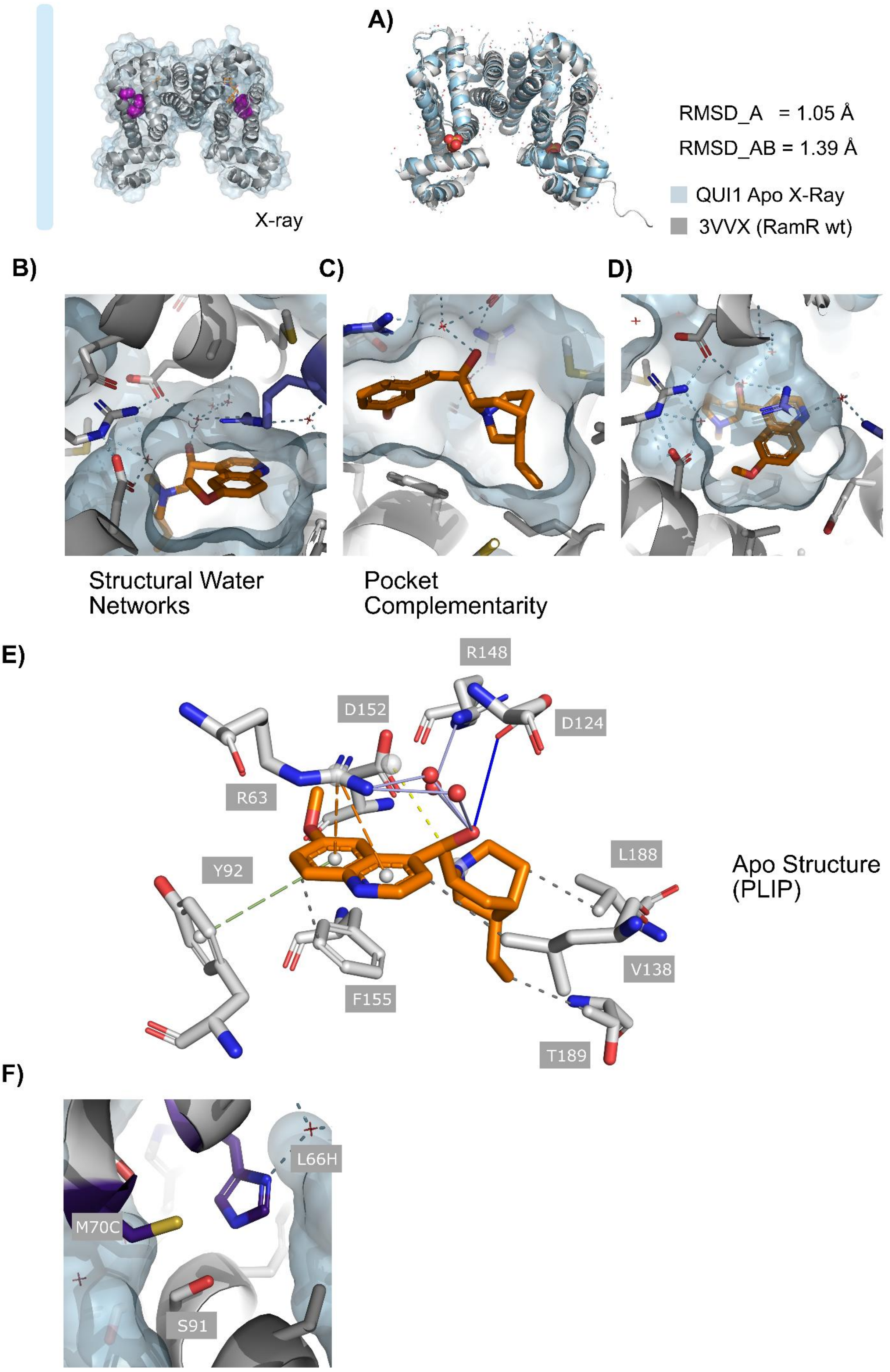

A)
X-ray
RMSD_A = 1.05 Å
RMSD_AB = 1.39 Å
QUI1 Apo X-Ray
3VVX (RamR wt)
B)
C)
D)
Structural Water Networks
Pocket Complementarity
E)
R148
D152
D124
R63
Y92
L188
V138
F155
T189
Apo Structure (PLIP)
F)
L66H
M70C
S91

**Supplementary Figure 7**. QUI1 Holo Structure Implicating Structural Water in Binding. **A)** Comparison of C-alpha RMSD between wild-type RamR (3VVX) and the holo QUI1 structure shows a low deviation (<2Å) both at the monomer and homodimer complex level. **B)** The key K63R (blue) contributes to a resolved hydrogen bonding network mediated by structural water. Notably the hydrogen-bonding network is opposed by hydrophobic residues on the other side, forming an amphiphilic pocked for QUI. **C)** Strong shape complementarity to QUI along with the many immediate interactions explain the low-micromolar affinity for QUI. **D)** Alternative view of the hydrogen bonding network mediated by structural water. **E)** PLIP diagram highlighting the key immediate residues. Notably, F155 and Y92 form π-interaction with QUI, while R63 forms a cation-π bond while mediating an indirect hydrogen bond with the hydroxyl of QUI.

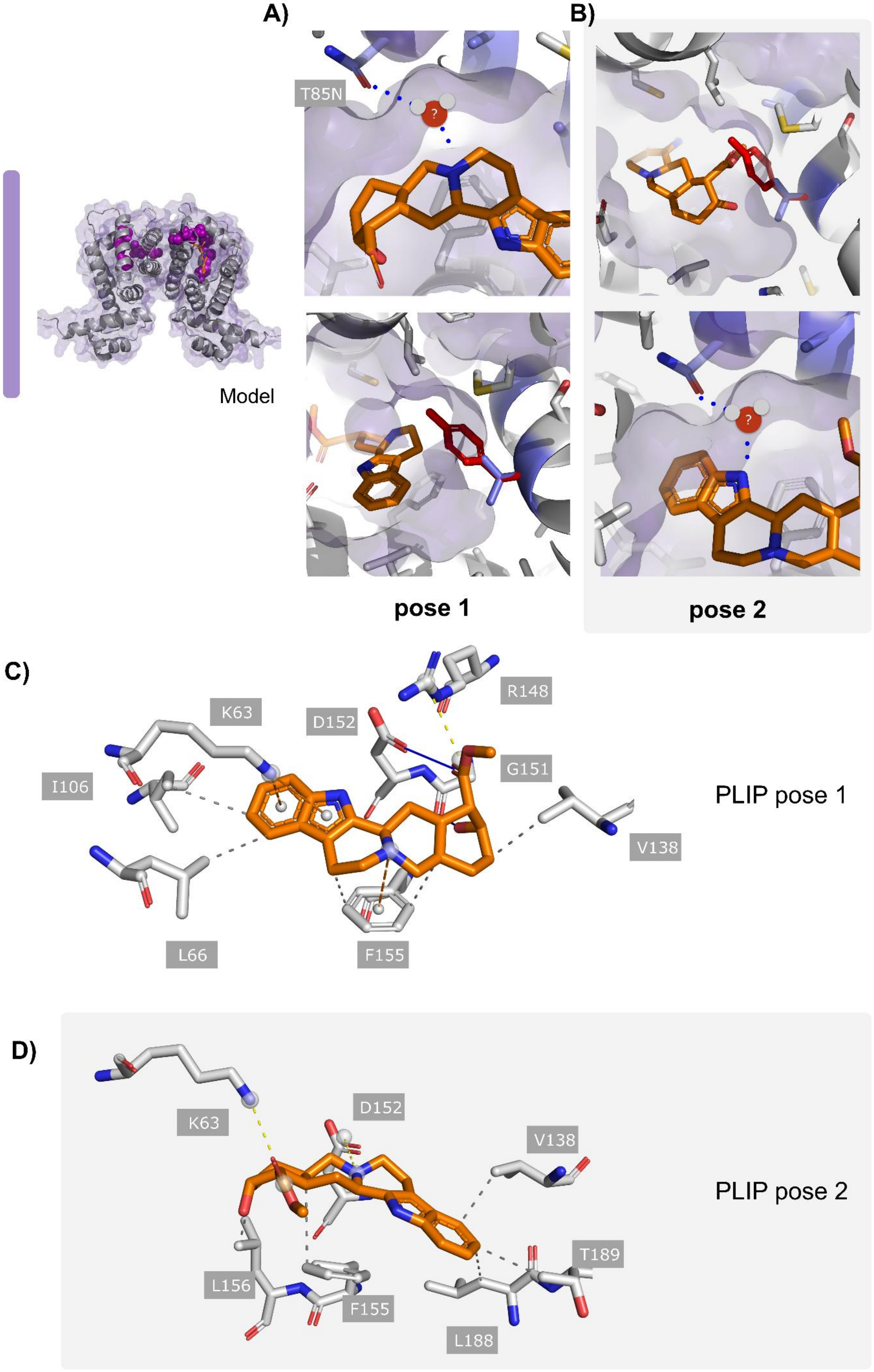

A)
B)
T85N
?
Model
pose 1
pose 2
C)
K63
D152
R148
G151
I106
V138
L66
F155
PLIP pose 1
D)
K63
D152
V138
T189
L156
F155
L188
PLIP pose 2

**Supplementary Figure 8.** RAU2 Holo Modelling to Determine Residues Driving Ligand Binding. Cross-referencing co-folding (Boltz-2, **(A)**) and docking (Vina **(B)**) poses indicated two viable poses which both correlate with the introduced mutations (RAU2: T85N I88A Y92V F142M M143S). T85N may mediate a water mediated hydrogen bond to either the indoloquinolizidine nitrogen (tertiary amine, in pose 1) or the pyrrole-type nitrogen in indole (pose 2). The Y92V substitution prevents a steric clash which would clash with either pose 1 and 2, which correlated with the low fold-induction of RAU in wild-type RamR (**Fig. 1B**) C) PLIP diagram of pose 1: K63 shows a cation-π interaction with RAU, while D152 and R148 mediate a hydrophilic interaction with the carboxylic acid of RAU (H-bond and/or salt-bridge). **D)** PLIP diagram of pose 2 showing a hydrophilic K62-carboxylic acid interaction, F155 as a cannonical π-interaction partner while D152 could form a hydrophilic interaction with the tertiary amine in RAU.

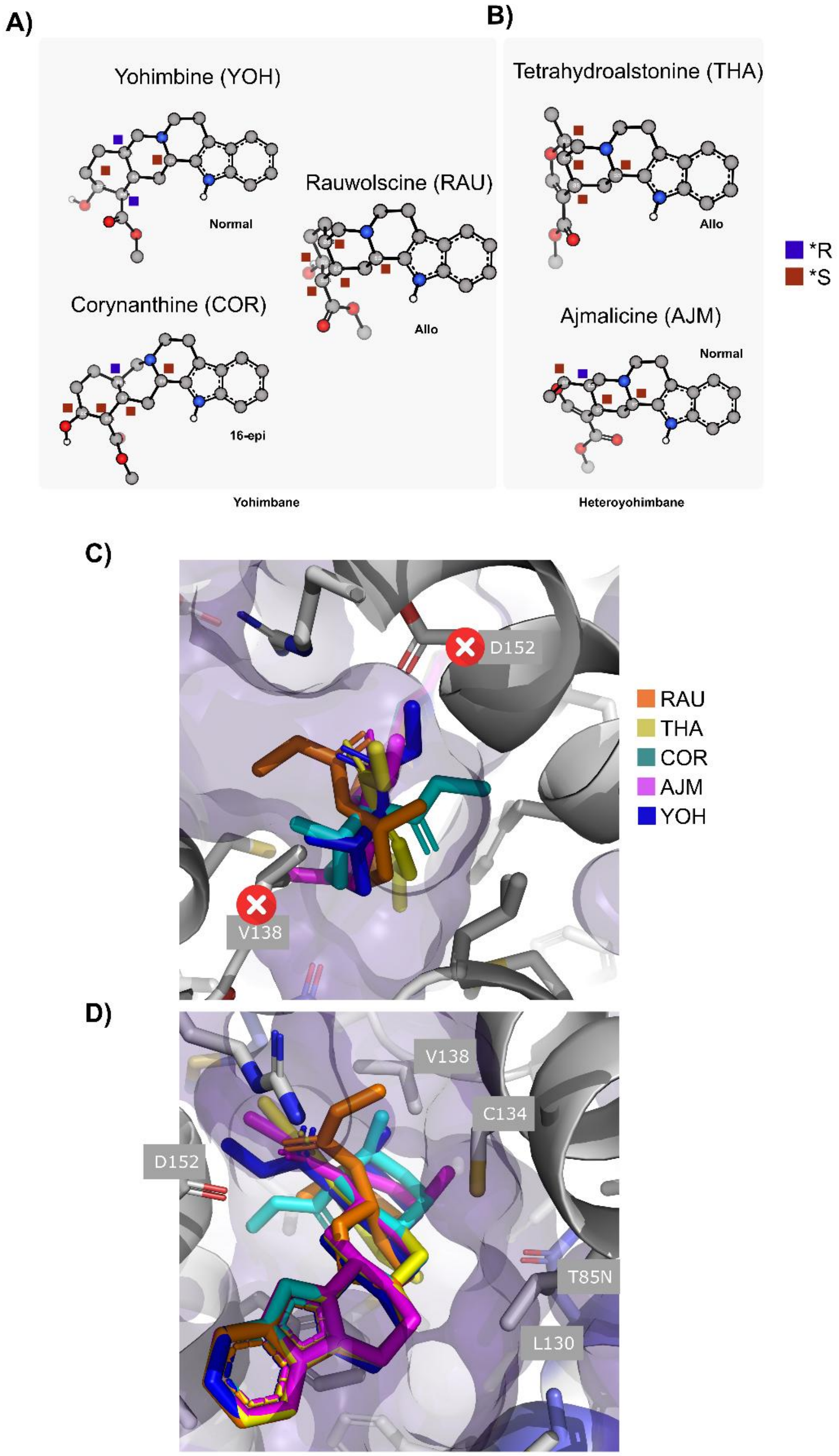


**Supplementary Figure 9.** RAU2 Holo Modelling to Determine Residues Driving Enantioselectivity. **A)** Structural Comparison of RAU and chemically related analogs in their dominant conformer: rauwolscine (RAU), yohimbine (YOH), and corynanthine (COR) share the pentacyclic yohimbane skeleton and differ only in the configuration at single stereocenters. RAU is the C-20 epimer of YOH (allo vs. normal D/E ring fusion), and COR is the C-16 epimer of YOH. **B)** Ajmalicine (AJM) and tetrahydroalstonine (THA) are heteroyohimbanes in which the saturated E-ring is replaced by a vinyl-ether. **C)** RAU2 modelled holo structure with analogs superimposed and aligned on the conserved indole skeleton. This implicates V138 and D152 (backbone) as steric sites which may explain the selectivity of RAU2. Both RAU and THA, which are colored orange and yellow are the only sterically compatible poses. This corresponds with the moderate fold-induction by THA experimentally. **D)** COR, AJM, and YOH maintain vdW clashes with the two aforementioned residues, while L130, T85N, and C134 may be further investigated for their role in discriminating the RAU analogs.

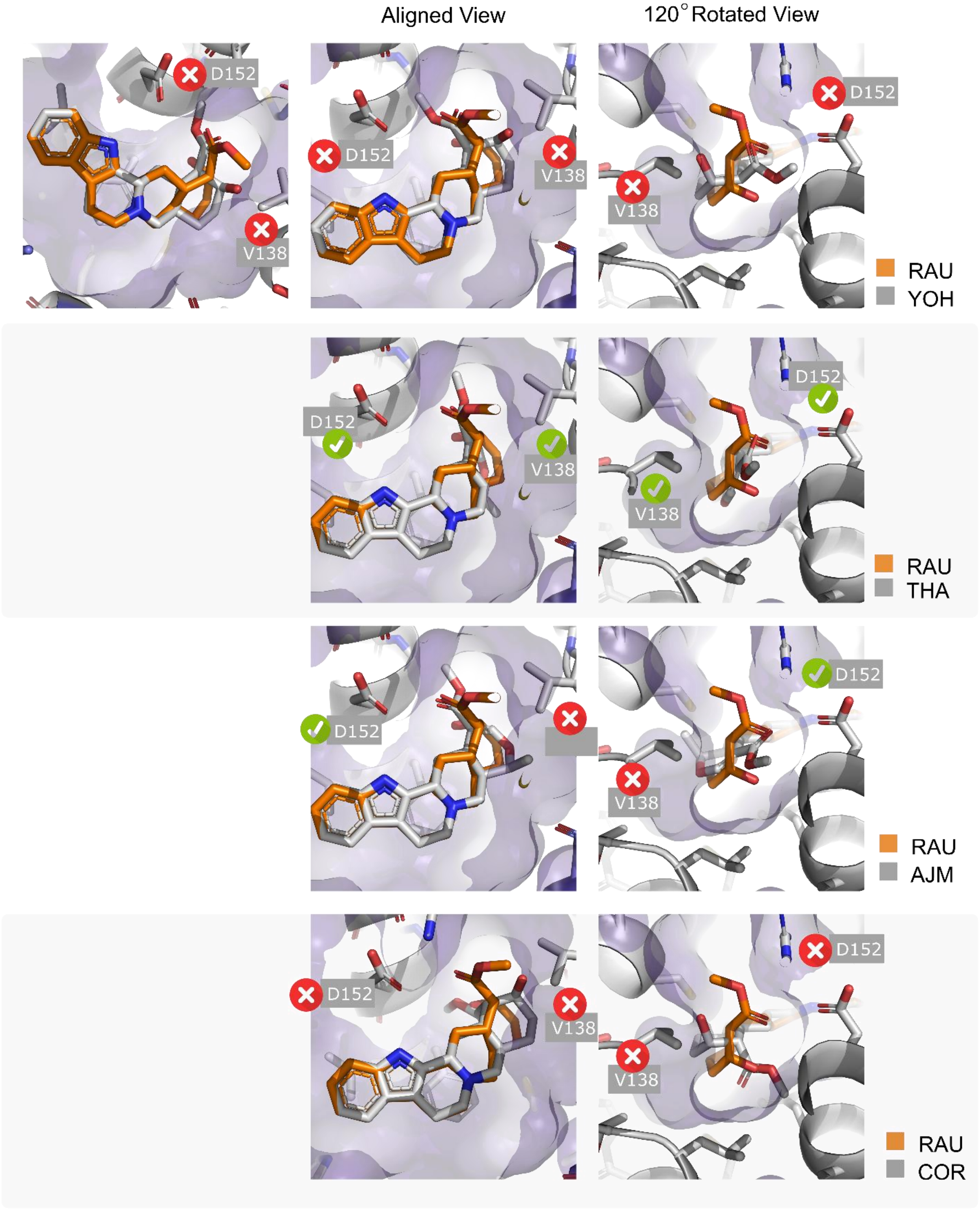


**Supplementary Figure 10.** Further RAU2 Holo Modelling to Determine Residues Driving Enantioselectivity. To clearly visualize the individual steric effects arising from isomers of RAU in RAU2, each are vizualized separately (from **Supp. Fig. 9**). RAU is always shown in orange and the alternative ligand in grey. Steric clashes from the grey ligand are shown with red crosses.

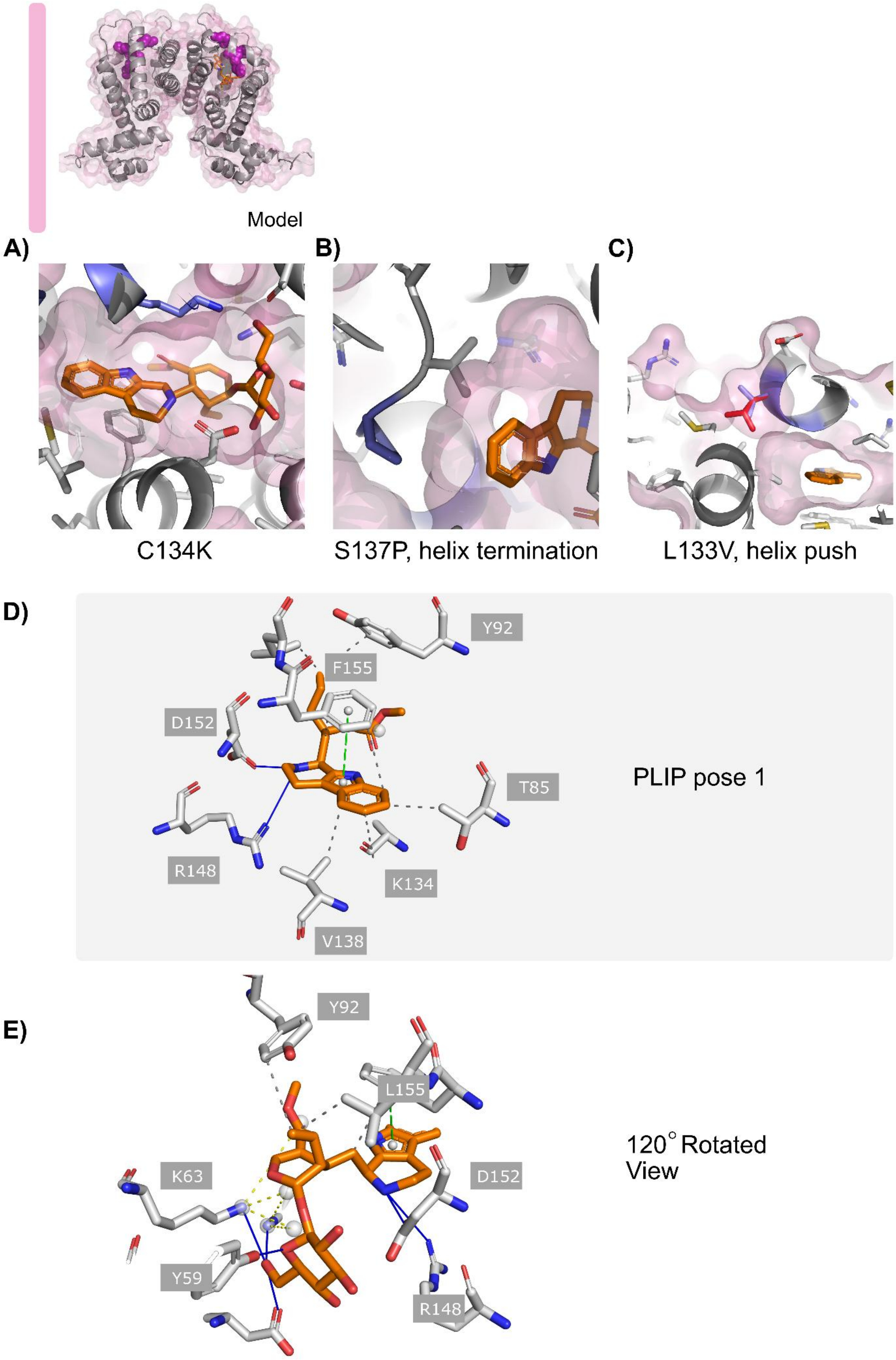

Model
A)
B)
C)
C134K
S137P, helix termination
L133V, helix push
D)
Y92
F155
D152
T85
R148
K134
V138
PLIP pose 1
E)
Y92
L155
K63
D152
Y59
R148
120° Rotated View

**Supplementary Figure 11.** STR1 Modelled Holo Structure. **A)** Cross-referencing co-folding (Boltz-2) and docking (Vina) poses indicates one viable pose for STR. This may explain the role of C134K, as the flexible and long sidechain reaches toward the hydroxyl of the glucose moiety while forming hydrophobic contacts with the indole substructure of STR. **B)** S137P enables the premature termination of helix α8 which affects both the distal β-bridge (V138, **Supp. Fig. 4**) and may push V138 toward STR forming a more complimentary hydrophobic pocket. **C)** The enlargement by L133V (also helix α8), may push the helix inward, forming a tighter binding pocket. **D)** PLIP diagram of modelled STR pose in STR1 shows both D152 and R148 may form hydrogen bonds with the tertiary amine, while V138, K134, T85, F155 and Y92 form a complementary hydrophobic pocket toward the indole substructure. **E)** Rotated view of D) enables visualizing residues that interact with the glucose moiety of STR.

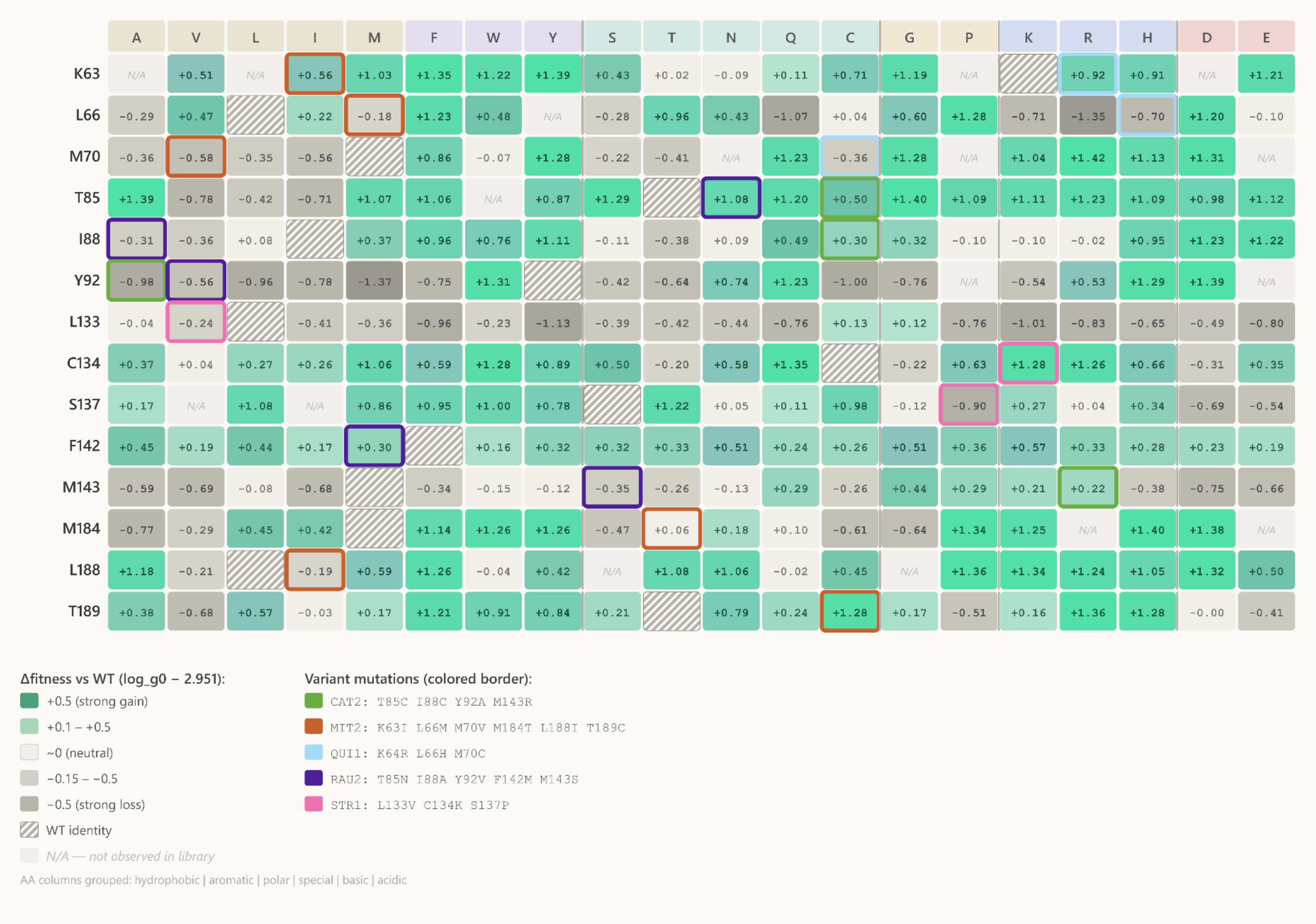


**Supplementary Figure 12.** Cross-referencing Growth-based Quantitative Sequencing (GROQ-seq) data with RamR mutations obtained in this work. Heatmap showing the change in uninduced transcription (Δlog_g0) relative to wild-type mean (2.951) for single amino acid substitution variants at the 14 residue positions targeted across the five engineered RamR MIA sensors. Teal indicates decreased uninduced transcription (stronger repression, higher DNA binding activity) and grey indicates increased uninduced transcription (weaker repression, lower DNA binding activity). Data from the NIST GROQ-seq RamR dataset (32,593 variants) (Spinner et al., 2026).

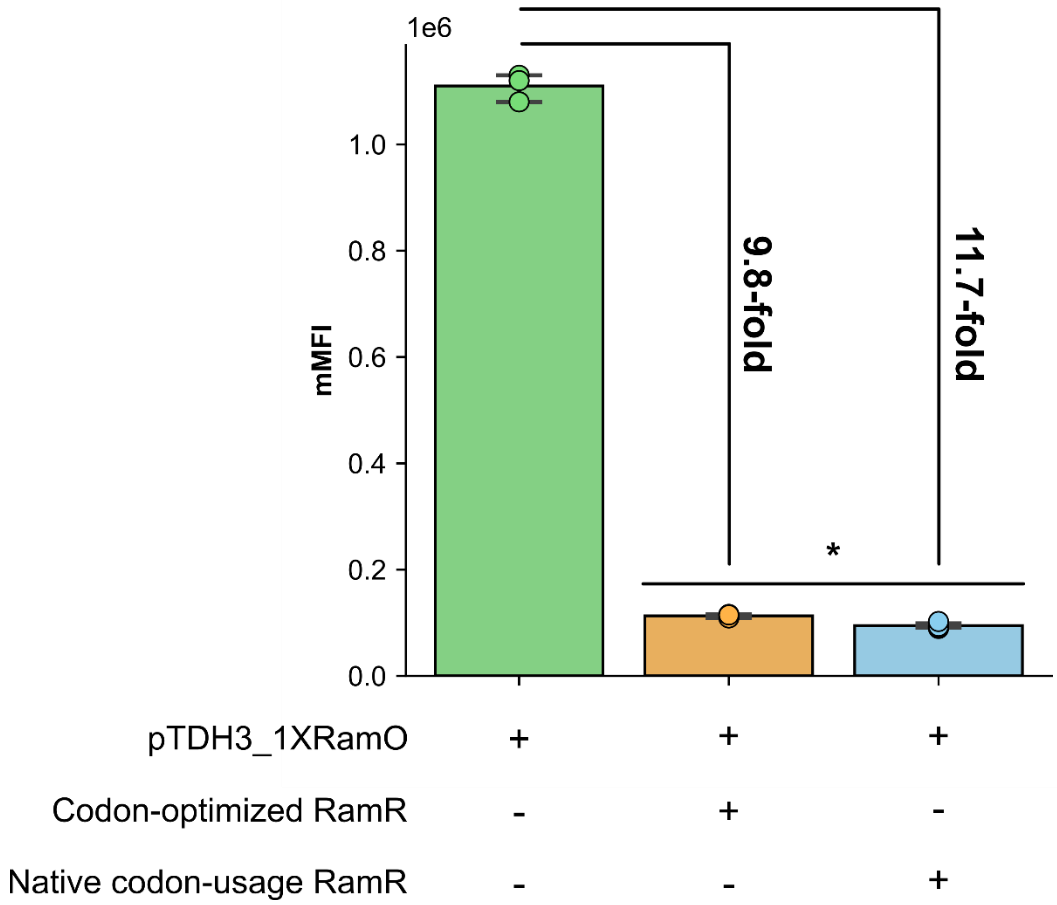


**Supplementary Figure 13.** Comparison of the repression efficiency of codon-optimized RamR for *S. cerevisiae* and the native non-codon optimized sequence in yeast. The reporter promoter used is pTDH3_1XRamO and data was acquired with flow cytometry after 6h. Measurements for each condition represent the average of three biological replicates. Error bars represent S.D.+/−the mean. Statistical significance was determined using an unpaired t-test. * p = 0.013.

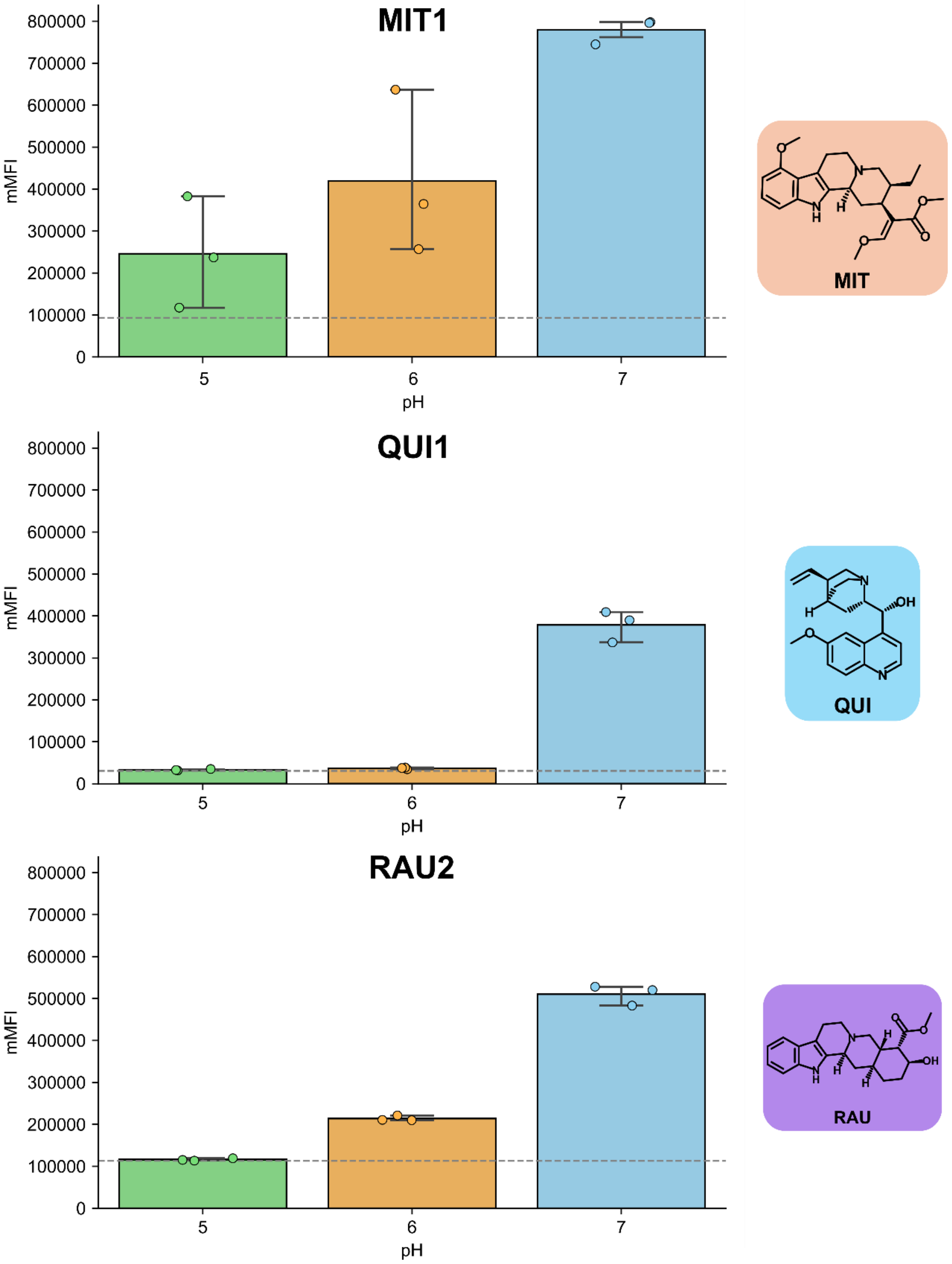


**Supplementary Figure 14.** Effect of pH on RamR  MIA biosensing in yeast. MIA ligands were supplemented at 100 µM (QUI1 and RAU) or 250 µM for MIT. Dashed grey line indicates the background signal mean value (no ligand). Measurements for each condition represent the average of three biological replicates. Error bars represent S.D.+/−the mean.

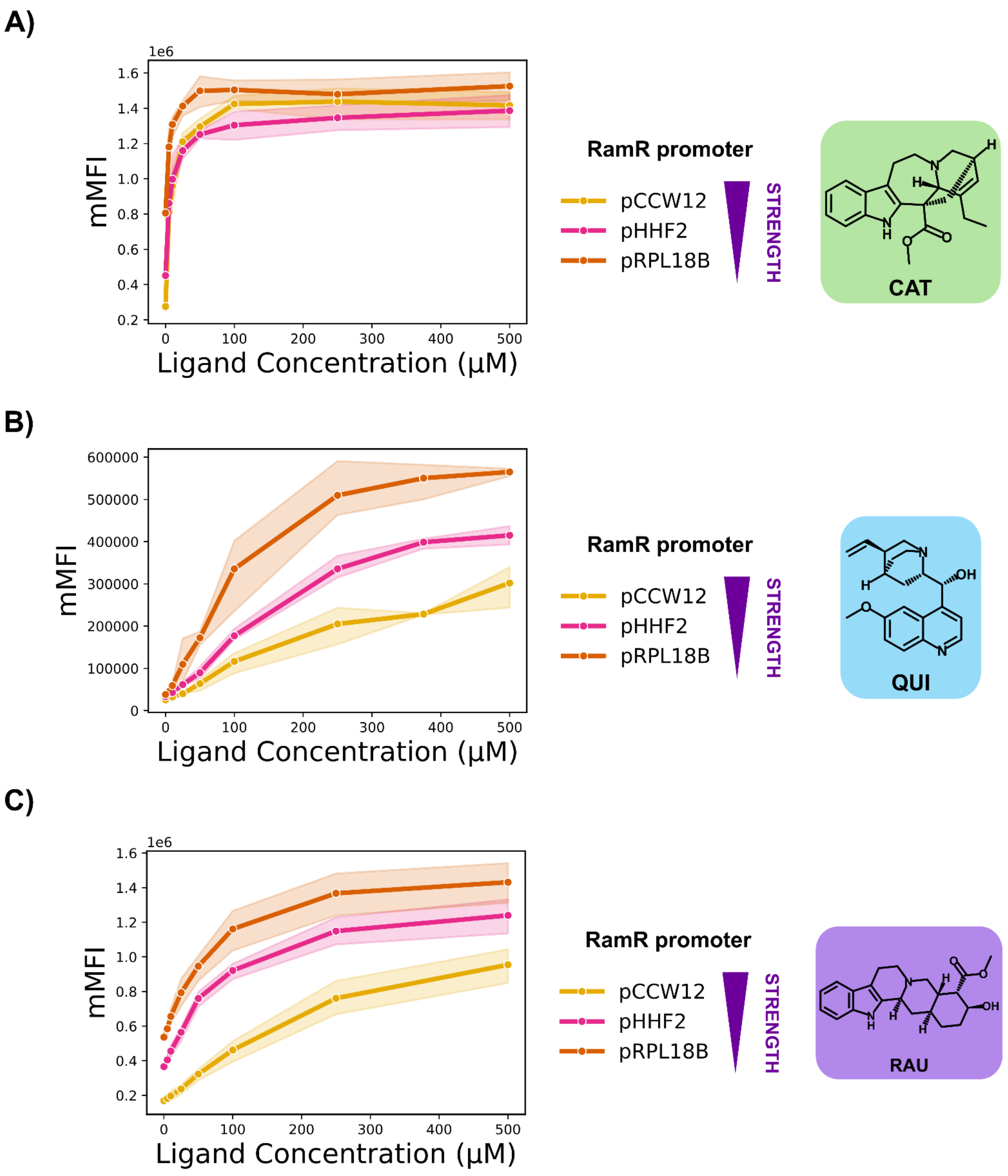


**Supplementary Figure 15.** Effect of RamR expression strength on biosensor transfer function for CAT2 (**A**), QUI1 (**B**) and RAU2 (**C**). RamR variants were expressed from CEN-ARS plasmids. Measurements for each condition represent the average of three biological replicates. Error bars represent S.D.+/−the mean.

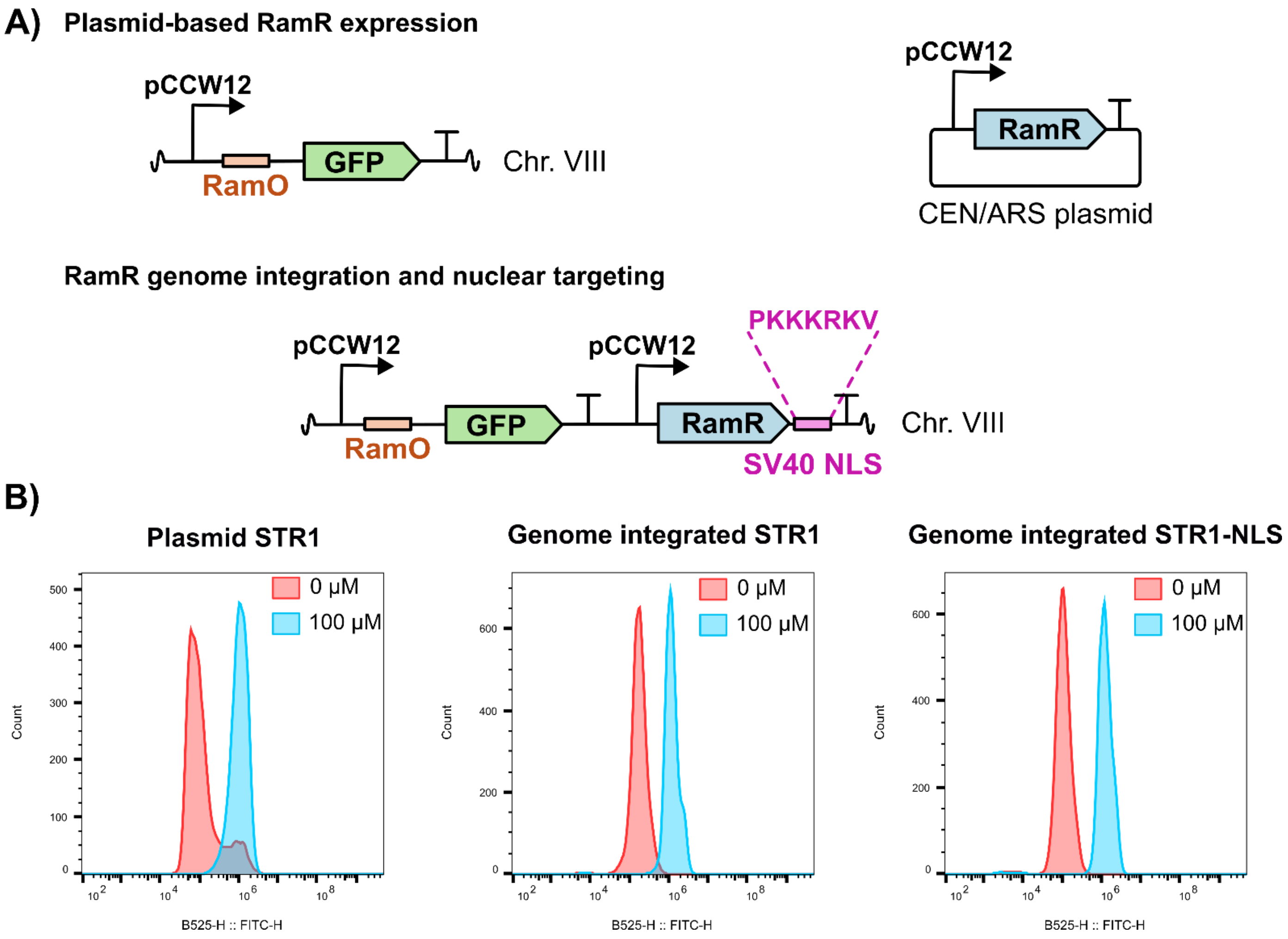


**Supplementary Figure 16.** Modulation of RamR biosensing by genomic integration and nuclear localization. **A)** Schematic representation of plasmid versus genome-integrated RamR circuits. **B)** Comparison of representative flow cytometry histograms for the STR1-sensor in the uninduced (red) and induced with 100 μM of STR (blue) states for plasmid-based or genome integrated RamR circuits.

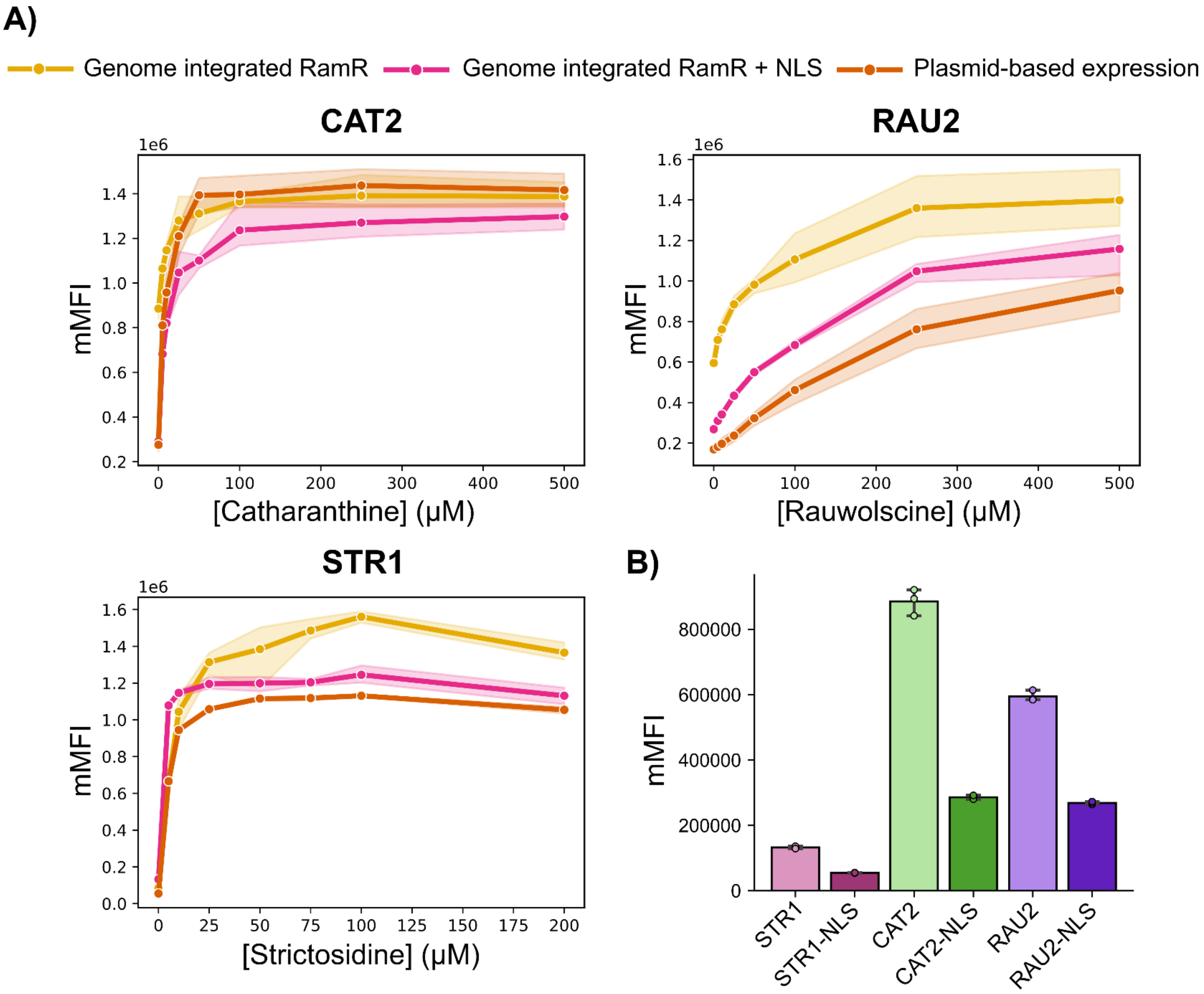


**Supplementary Figure 17.** Comparison of RamR MIA biosensor dose–response in yeast (**A**) and background signal in the absence of ligand (**B**) for plasmid-based versus genome-integrated RamR circuits, with or without an SV40 NLS tag. Measurements for each condition represent the average of three biological replicates. Error bars represent S.D.+/−the mean

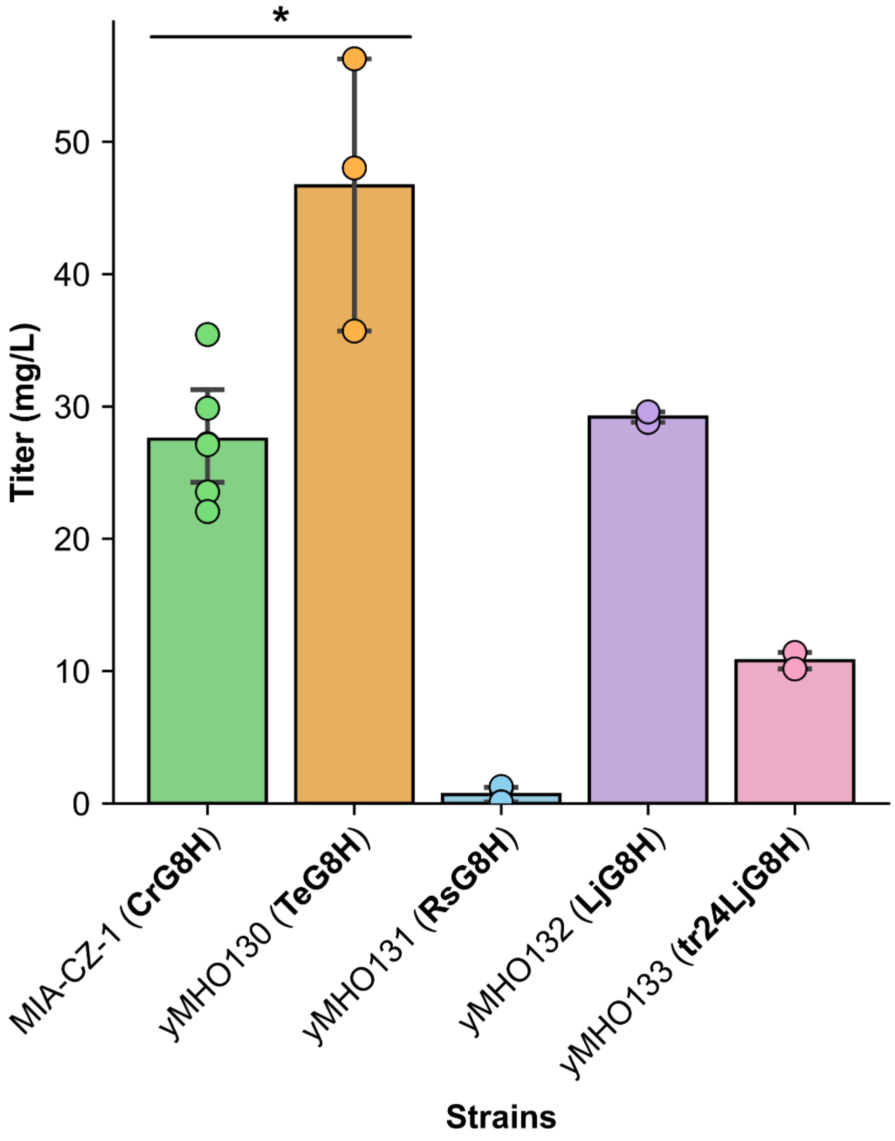


**Supplementary Figure 18.** Geraniol hydroxylase (G8H) screen in MIA-CZ-1 for strictosidine production. Strains were grown in 3xSC 2 % glucose 3mM Trp 1% Pbov at 25°C for six days. Abbreviations are: Cr, *Catharantus roseus*; Te, *Tabernaemontana elegans*; Rs, *Rauvolfia serpentina*; Lj, Lonicera *japonica*. Measurements for each condition represent the average of at least three biological replicates. Error bars represent S.D.+/−the mean. Statistical significance was determined using an unpaired t-test. * $p = 0.0055$.

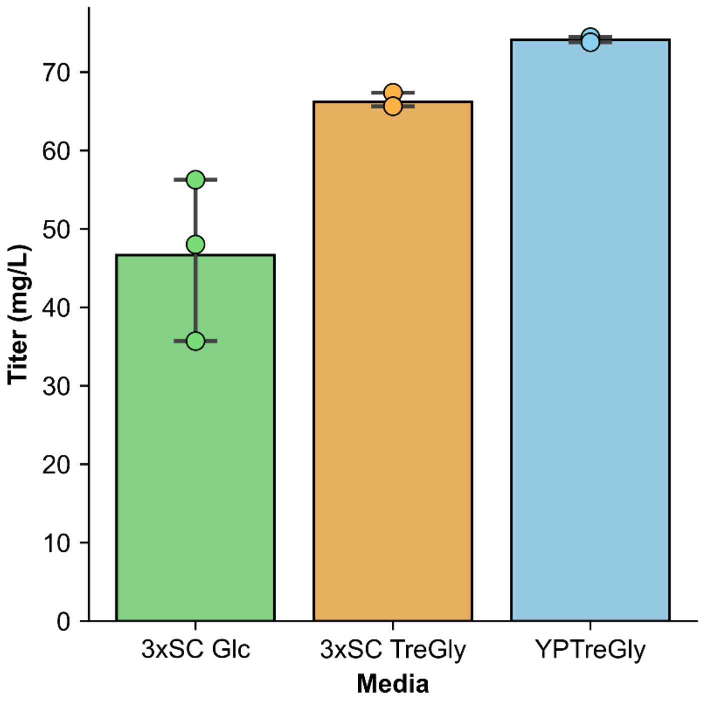


**Supplementary Figure 19.** Comparison of production media for STR production by yMHO130. 3xSC is up concentrated yeast synthetic complete media with 1% bovine peptone and 3 mM tryptophan. Glucose (Glc) and trehalose (Tre) were supplemented at 2 % w/v. Glycerol (Gly) was supplemented at 10 % w/v. YP refers to yeast extract and peptone media base and was also supplemented with 3 mM tryptophan. Strains were grown for 6 days at 25 °C. Measurements for each condition represent the average of at least three biological replicates. Error bars represent S.D.+/−the mean.

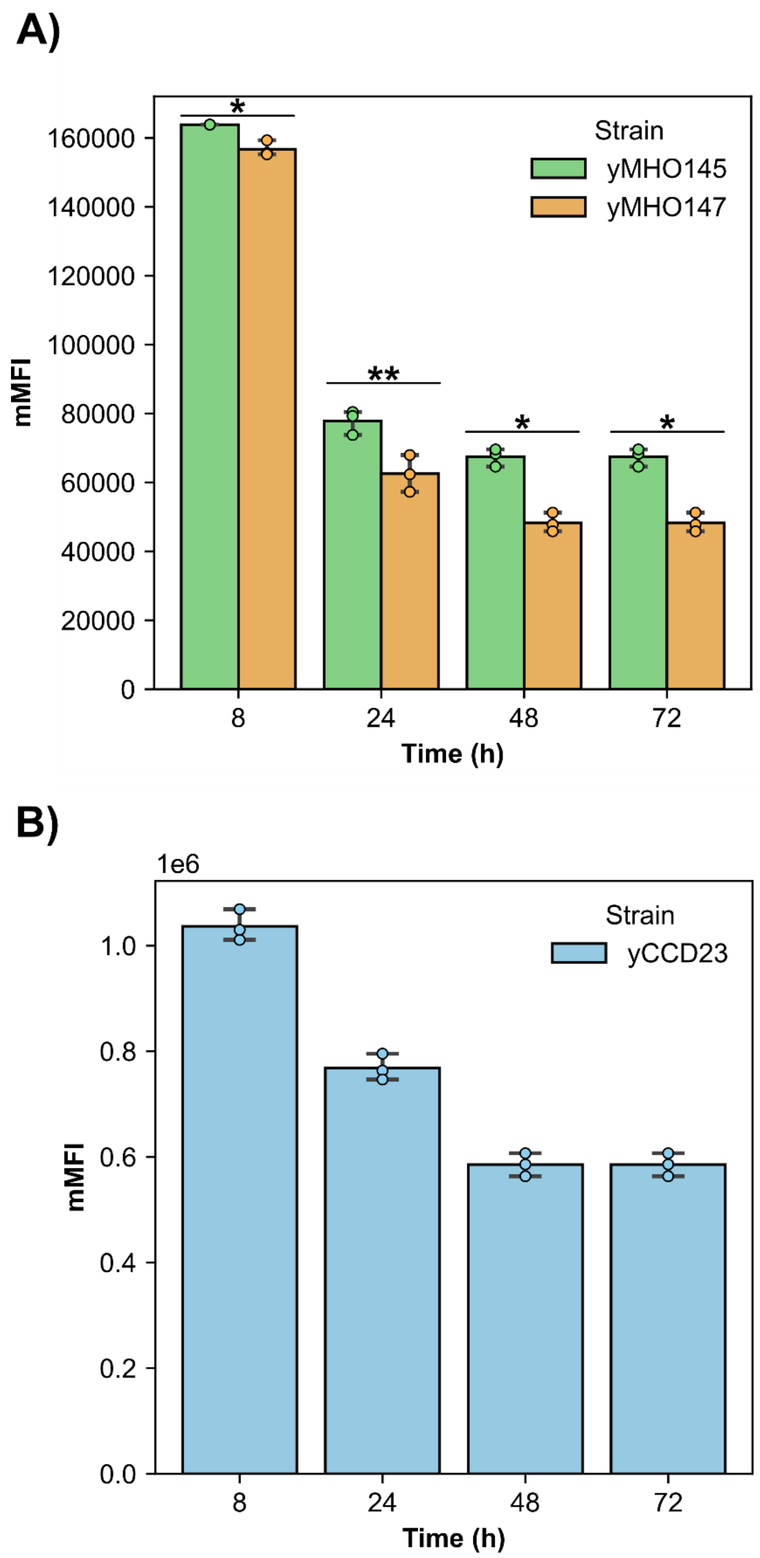


**Supplementary Figure 20. A**) Strictosidine biosensing in STR production strain yMHO145 and non-producer CrSLS knockout yMHO147 over time. **B)** Basal transcriptional activity of RamR reporter promoter pCCW12_1XRamO (strain yCCD23) over time without repressor expressed. Strains were grown in YPTreGly at 25°C. Measurements for each condition represent the average of at least three biological replicates. Error bars represent S.D.+/−the mean. Statistical significance was determined using an unpaired t-test. * $p < 0.01$, ** $p < 0.05$.

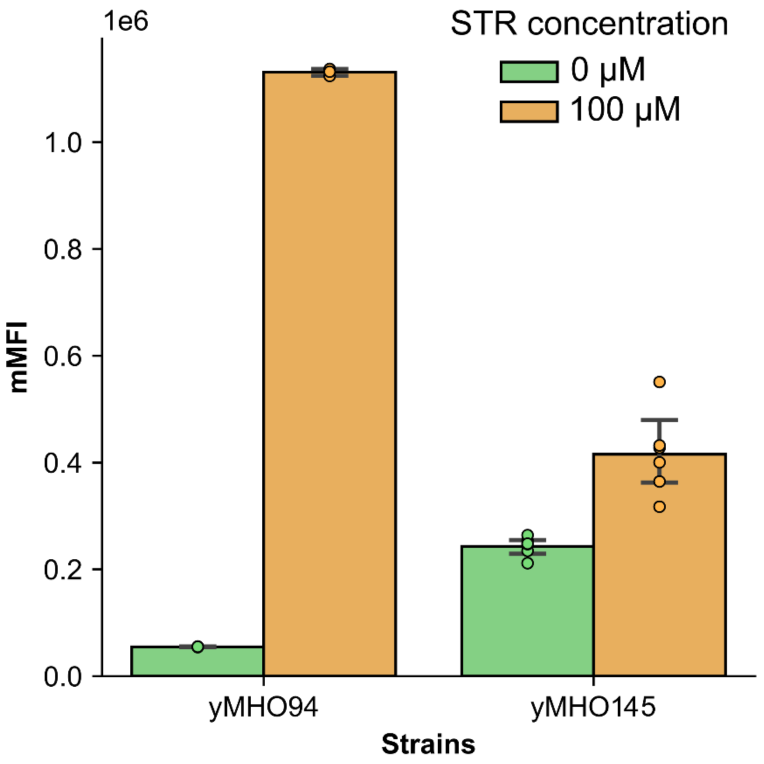


**Supplementary Figure 21.** Comparison of genome integrated STR1-NLS biosensor signal without ligand (green) and induced with 100 μM STR (orange) in a WT background yeast strain (yMHO94) and in a strictosidine producer (yMHO145). Strains were grown in SC 2% glucose pH 7 media at 30°C and fluorescence was measured with flow cytometry after 6 h. Measurements for each condition represent the average of at least three biological replicates. Error bars represent S.D.+/−the mean.

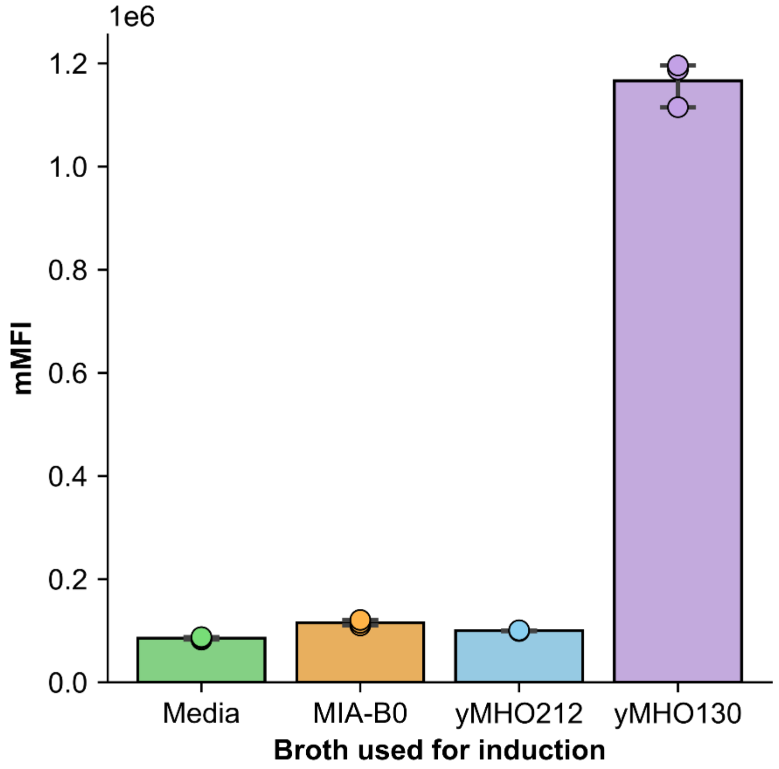


**Supplementary Figure 22.** Strictosidine biosensing with yMHO94 (STR1-NLS sensor) induced with supernatant from different yeast strains (1:20 volume ratio). Supernatants used were fresh YPTreGly production media (green), MIA-B0 (WT parental yeast strain, orange), yMHO212 (yMHO130 ΔCrSLS, interrupted MIA pathway, blue) and yMHO130 (STR producer, purple). Biosensing was carried in buffered YPD pH 7 media and fluorescence was measured with flow cytometry after 6 h. Measurements for each condition represent the average of three biological replicates. Error bars represent S.D.+/−the mean

### Supplementary Tables

**Supplementary Table 1. Sequences of synthetic DNA fragments used in this study.** For coding sequences, start codons are highlighted in bold and stop codons are underlined. For promoter sequences the RamO operator sequence is shown in red.

| Name | Type | Reference | Nucleotide sequence |
|---|---|---|---|
| RamR_nat | CDS from *Salmonella typhimurium.* | (d'Oelsnitz et al., 2022) | **ATG**GTTGCTCGCCCAAAGTCTGAGGACAAAAAGCAGGCATTGCTTG<br>AAGCGGCAACTCAAGCCATCGCGCAATCAGGCATTGCCGCTAGTAC<br>CGCTGTAATTGCACGCAATGCGGGAGTTGCGGAAGGGACGTTGTTC<br>CGCTATTTCGCAACGAAAGATGAGTTGATCAACACCCTTTACTTACA<br>TTTGAAACAGGACCTGTGCCAATCAATGATCATGGAATTGGATCGTT<br>CTATTACTGACGCTAAGATGATGACCCGTTTTATCTGGAACAGTTATA<br>TTAGCTGGGGATTGAACCACCCAGCTCGCCATCGTGCCATTCGTCA<br>GTTGGCGGTTTCTGAAAAGTTGACGAAGGAAACCGAACAACGCGCG<br>GATGATATGTTCCCGGAGTTACGCGACTTGTGCCACCGTAGTGTTCT<br>TATGGTGTTTATGTCCGACGAGTACCGCGCCTTCGGCGACGGGTTG<br>TTCTTGGCGCTTGCTGAGACGACTATGGATTTCGCTGCGCGCGACC<br>CGGCTCGCGCTGGTGAGTACATTGCGTTGGGCTTCGAGGCTATGTG<br>GCGCGCACTTACGCGCGAAGAGCAGTAA |
| RamR_Sc_opt | CDS from *Salmonella typhimurium* codon optimized for *S. cerevisiae*. | This study. | **ATG**GTCGCCAGACCAAAGTCTGAAGACAAGAAGCAAGCTTTGTTGG<br>AAGCTGCTACTCAAGCTATCGCTCAATCTGGTATCGCCGCTTCCACT<br>GCTGTCATTGCTAGAAACGCCGGTGTTGCTGAAGGTACATTATTCAG<br>ATACTTCGCTACCAAAGATGAATTGATCAACACTTTGTACTTGCATTT<br>GAAGCAAGATTTATGTCAATCCATGATTATGGAATTGGACAGATCCA<br>TTACTGATGCTAAGATGATGAATAGATTCGCTTGGAACTCTGTTATTT<br>CTTGGGGTTTGAATCACCCAGCTAGACACAGAGCTATCAGACAATTG<br>GCTGTTTCTGAGAAGTTGACCAAGGAAACTGAACAAAGAGCTGATGA<br>CATGTTCCCAGAACTACGTGACTTGTGTCACCGTTCTGTTTTGATGG<br>TTTTCATGTCAGACGAATACAGAGCTTTCGGTGATGGTTTGTTTTTG<br>GCTTTAGCTGAAACCACCATGGACTTTGCTGCCAGAGATCCAGCCC<br>GTGCTGGTGAATATATTGCCTTGGGTTTCGAAGCTATGTGGAGAGCA<br>CTGACCAGAGAAGAACAATGA |
| SV40 NLS | C-ter nuclear localization tag derived from Simian Virus 40 large T-antigen (PKKKRKV). | (Li et al., 2015) | CCAAAGAAGAAGAGAAAGGTA |
| TeG8H | CDS from *Tabernaemontana elegans* codon optimized for *S. cerevisiae*. | (Y. Zhang et al., 2023) | **ATG**GACTATCTAACATTGGTTTTGGGTTTGTTGTTTGCGCTCACTTTC<br>TACCAAGGTTTGTCTTACTTGTCTAGAAAATCTAAGAAGTTGCCACCT<br>GGTCCAGCTCCACTCCCATTCATTGGTAACTTGCATTTGTTGGGTGA<br>CCAACCACACAAGTCTTTAGCCAAGTTGTCCAAGAAGCACGGTCCG<br>TTGATGTCTTTGAAGTTGGGTCACATTACTACTGTTGTTATCTCCAGC<br>TCCGGTATGGCCAAAGAAGTCTTGCAAAAGCAAGACTTGGCTTTTTC<br>CTCGCGTTCTGTTCCAGATGCTTTGCACGCTCACAACCAATTCCAAT<br>TCTCAGTCGTCTGGTTGCCAGTTGCCTCTAGATGGAGATCCTTGAGA<br>AAGATTTTGAACTCCAACATTTTCTCTGGTAACAGATTGGACGCTAAT<br>CAACACCTAAGATGTCGTAAGGTCCAAGAATTAATTGCTTACTGTAG<br>AAAGAACTGTCAAACTGGTGAAGCTGTCGACGTTGGTCGTGCTGCT<br>TTCAGAACCTCCTTAAATTTATTGTCTAACACTATCTTCTCCAAGGAC<br>TTGACCGATCCATACTCTGACTCCGCTAAGGAATTCAAGGATTTAGT<br>CTGGAACATCATGGTCGAAGCCGGCAAACCAAACTTGGTCGATTTCT<br>TCCCAGTCCTTGAAAAGGTTGACCCACAAGGTATCAGAAAAAGAATG<br>ACCGTTCACTTCGGTAAGGTCTTGAAGCTATTCAACGGTCTAATCAA<br>CGAAAGATTGGAACAAAAGCGATTGCAATCTGGTAACAACGACGTGT |

| | | | |
|---|---|---|---|
| | | | TAGATGTTTTGTTGACCACCTCTGAAGAATCCCCAGAAGAAATCGAC<br>AGAACTCATATCGAGAGAATGTGTTTGGACTTGTTCGTTGCCGGTAC<br>CGACACCACCAGCTCCACTTTGGAATGGGCTATGGCTGAAATGTTG<br>AAGAACCCAGACAAGATGCAAAAGACTCAAGCTGAAATTGCTGAAGT<br>TATTGGTAAGGGTAAGGCCATTGAAGAAGCTGATGTCGCTAGATTGC<br>CATACTTGAGATGCGTTGTCAAGGAAACCTTGAGAATTCACCCACCA<br>GTTCCATTCTTGATCCCAAGAAAAGTTGACCAAGATGTTGAAGTCTG<br>TGGTTACAAGGTCCCAAAGGGTTCTCAAGTTTTAGTAAATGCTTGGG<br>CTATTGGTAGAGATGGTACTGTTTGGGAAGACCCATTAGCTTTCAAG<br>CCAGAAAGATTCTTTGAATCTGAAATCGATATCAGAGGTAGAGACTT<br>CGAAGTCATTCCATTTGGTGCTGGTAGAAGAATCTGTCCAGGCTTGC<br>CTTTGGCCGCTCGTATGGTTCCTGTTATGTTAGGTTCTCTATTGAACT<br>CTTTCAACTGGAAGTTGGAAGGTGGTATCGCCCCAAAGGATTTGGA<br>CATGGATGAAAAATTCGGTATCACTTTGCAAAAGGCTCATCCATTGC<br>GGGCTGTTCCATCTCCATTGTAA |
| RsG8H | CDS from *Rauvolfia serpentina* codon optimized for *S. cerevisiae*. | (Y. Zhang et al., 2023) | **ATG**GACTATTTGACCATTACCCTAGGTTTGTTGTTCGCCTTGACTTTC<br>TACCAAGGTTTATCTTACTTGTCTCGTAGATCCAAGAAGTTGCCACC<br>AGGCCCAGCTCCATTGCCTATCATCGGTAACTTGCATATGTTGGGTG<br>ACCAACCACACAAGTCTTTGGCTAAATTGTCCAAGAAGCACGGTCCA<br>ATTATGTCTCTAAAGTTGGGTCAAATCACCACCGTTGTTGTTAGTTCA<br>TCCGGTATGGCCAAGGAAGTCTTGCAAAAGCATGACTTGGCTTTCTC<br>CTCCCGTTCCATCCCAAACGCCTTGCACGCCCACAACCAATACCAAT<br>TCTCTGTTGTCTGGTTGCCAGTCGCTTCCAGATGGAGATCTTTGAGA<br>AAGACCTTGAACTCCAACATTTTCTCTGGTAACAGATTGGATGCTAA<br>CCAACACCTGAGATCTAGAAAGGTCGAAGAATTGATTGCTTACTGTC<br>GTAAGAACTGTCAAACTGGTGAAGCCGTCGATGTTGGTAGAGCTGC<br>TTTCAGAACCTCTTTGAACTTGTTATCTAACTTGATTTTTTCTAAGGAT<br>TTGACAGATCCATACTCTGACTCTGCTAAGGAATTCAAAGACTTGGT<br>TTGGAACATCATGGTTGAAGCTGGTAAGCCAAATTTAGTTGACTACT<br>TCCCAGTCTTGGAAAAGGTCGACCCACAAGGAATCAGACGTAGAAT<br>GACTGTTCACTTCGGTAAGGTTTTGAAGTTATTCGATGGTTTGGTTAA<br>CGAAAGATTGGAACAAAGAAGATCCCGTGGTGGTAAAAACGATGTA<br>CTCGATGTCTTATTGACCAACTCTGAAGAAAACCCAGAAGAAATTGA<br>CAGAACTCACATCGAAAGAATGTGTTTGGACTTATTCGTCGCTGGTA<br>CTGACACTACTTCCAGCACGCTAGAATGGGCTATGGCTGAAATGTTG<br>AAGAACCCAGACAAGATGAAGAAGACTCAAGCAGAATTAGCTGAAG<br>TTATCGGTAGAGGTAAAGCTATCGAAGAATCTGATTTGCCAAGATTA<br>CCATACCTAAGATGCGTCGTCAAGGAAACCTTGAGAATTCACCCACC<br>AGTTCCATTCTTGATCCCTAGAAGAGTTGAACAAGATGTTGAGGTTT<br>GTGGTTACAAGGTTCCTAAGGGTAGTCAAGTTTTGGTGAACGCTTGG<br>ACTATCGGTCGTGACGAAGCTGTCTGGGAAGACGCTTTGGCCTTTA<br>AGCCAGAAAGATTCTTAGAATCCGAACTGGATATCAGAGGTAGAGAC<br>TTCGAACTCATTCCATTTGGTGCTGGTCGTAGAATCTGTCCAGGTCT<br>TCCGTTGGCTTTGAGAACCGTCCCATTGATGGTCGGTTCCTTGTTGA<br>ATTCTTTCAATTGGAAGCTGGAAGGTGGTATTACTCCAAAGGATTTG<br>GACATGGAAGAAAAGTTCGGTATCACTTTGCAAAAAGCCCATCCATT<br>AAGAGCTGTTCCAATTCCATTTTGA |
| LjG8H | CDS from *Lonicera japonica* codon optimized for *S. cerevisiae*. | (Wang et al., 2023) | **ATG**GATTTCTTTACCATCGCCTTGTCTTTGTTGTTTGCTTTTACCTTG<br>TTCAAGGCTATCATCTCTTTAACCAAGTCTGGTAAGAACAAGAACTT<br>GCCACCAGGTCCAGCTCCATTGCCAATTATTGGTTCCTTGCACAAGT<br>TGGGTGACCAACCACACCAAAGTCTAGCCAAGTTGGCTAAGATCTA<br>CGGTCCAATCATGTCTTTGAAGTTAGGTAGAATCACCACCGTCGTTA<br>TTAGCTCTTCTGATGCTGCTAAGGAAGTCTTGCAAAAGAAGGATCTC<br>GCTTTCTCTTCTAGACACGTTCCAGACGCTTTGCACGCTCACAACCA<br>ATTCAAGTTCAGTGTTGTTTGGTTACCAGTCGCTCCACAATGGAGAT<br>CCTTGAGGAAGATTCTGAACTCCAACATCTTCTCTGGTAACAGATTG<br>GATGCTAACCAACACTTAAGATGTAGAAAGGTCGAAGAACTAATCAC<br>TTACTGTCGTAAATCCTCCCACTCCGGTGACGCTGTCGATATTGGTA<br>GAGCTGCCTTCAGAACCTCTTTGAACTTGTTGTCGAACACTATTTTTT |

| | | | |
|---|---|---|---|
| | | | CAAAGGACTTGACTGACCCATACCAAGATTCTGCTAAAGAATTCAAG<br>GAGTTGGTCGGTAACATTATGGTTGAAGCCGGTAAGCCAAACTTAGT<br>TGACTTCTTCCCATCTTTGAAGAAGATTGATCCTCAAGGTATCCGTC<br>ACAGAATGACTATCCATTTCGGTAAGGTTTTGGAATTATTCGGTGGT<br>TTGATCAACGAAAGATTGGAACTTAAATCTTCCCAAAAGTCTGACGA<br>AAAGAACGACGTCCTAGATGTCTGCTTGTCCATCTCCAAGGAAAACC<br>CAGATGAAATTGACAGAACTCATATTGAACGTATGTGTTTGGACTTG<br>TTCGTTGCTGGTACTGATACCACATCCTCCACTTTGGAATGGGCTAT<br>GGCCGAAGTTTTGAGAAACCCAGAAACCTTGGTCAAGGCCAAGGCT<br>GAATTGGAAGAAGTCATCGGTAAGGGTAAAATCTTGGCTGAAGCTG<br>ATGTCTCCAATTTGCCTTATTTGAGATGTATCGTCAAGGAAACTTTGA<br>GAATTCATCCACCTGTTCCTTTCTTGATTCCAAGAAAGGTTGAAATG<br>GACGTTGAAGCTTGTGGCTACACTGTTCCAAAGAATTCTCAAGTTTT<br>CGTCAATGTCTGGGCTATTGGCAGAGATCCAACTTTGTGGGAAGAC<br>CCATTGTCCTTCAAGCCAGAAAGATTCATGGCTTCCGAATTGGACGT<br>TAGAGGTCGTGACTTCGAATTGTTGCCATTCGGTGCTGGTAGAAGAA<br>TCTGTCCAGGTCTGCCATTGGCCATCAGAATGGTTCCAGTTATGTTG<br>GGTTCTTTGGTTAACTCTTTCAACTGGAAACTAGATGGTGGTATTGC<br>TCCAAAGGAATTGGACATGGGTGAAAAATTTGGTATCACCTTAGCAA<br>AGGCTCAATCTCTAAGAGCCGTCCCATCTCCATTGGTGTGA |
| tr24LjG8H | CDS from *Lonicera japonica* codon optimized for *S. cerevisiae* (truncation first 24 aminoacids). | (Wang et al., 2023) | **ATG**AAGTCTGGTAAGAACAAGAACTTGCCACCAGGTCCAGCTCCATT<br>GCCAATTATTGGTTCCTTGCACAAGTTGGGTGACCAACCACACCAAA<br>GTCTAGCCAAGTTGGCTAAGATCTACGGTCCAATCATGTCTTTGAAG<br>TTAGGTAGAATCACCACCGTCGTTATTAGCTCTTCTGATGCTGCTAA<br>GGAAGTCTTGCAAAAGAAGGATCTCGCTTTCTCTTCTAGACACGTTC<br>CAGACGCTTTGCACGCTCACAACCAATTCAAGTTCAGTGTTGTTTGG<br>TTACCAGTCGCTCCACAATGGAGATCCTTGAGGAAGATTCTGAACTC<br>CAACATCTTCTCTGGTAACAGATTGGATGCTAACCAACACTTAAGAT<br>GTAGAAAGGTCGAAGAACTAATCACTTACTGTCGTAAATCCTCCCAC<br>TCCGGTGACGCTGTCGATATTGGTAGAGCTGCCTTCAGAACCTCTTT<br>GAACTTGTTGTCGAACACTATTTTTCAAAGGACTTGACTGACCCATA<br>CCAAGATTCTGCTAAAGAATTCAAGGAGTTGGTCGGTAACATTATGG<br>TTGAAGCCGGTAAGCCAAACTTAGTTGACTTCTTCCCATCTTTGAAG<br>AAGATTGATCCTCAAGGTATCCGTCACAGAATGACTATCCATTTCGG<br>TAAGGTTTTGGAATTATTCGGTGGTTTGATCAACGAAAGATTGGAAC<br>TTAAATCTTCCCAAAAGTCTGACGAAAAGAACGACGTCCTAGATGTC<br>TGCTTGTCCATCTCCAAGGAAAACCCAGATGAAATTGACAGAACTCA<br>TATTGAACGTATGTGTTTGGACTTGTTCGTTGCTGGTACTGATACCA<br>CATCCTCCACTTTGGAATGGGCTATGGCCGAAGTTTTGAGAAACCCA<br>GAAACCTTGGTCAAGGCCAAGGCTGAATTGGAAGAAGTCATCGGTA<br>AGGGTAAAATCTTGGCTGAAGCTGATGTCTCCAATTTGCCTTATTTG<br>AGATGTATCGTCAAGGAAACTTTGAGAATTCATCCACCTGTTCCTTTC<br>TTGATTCCAAGAAAGGTTGAAATGGACGTTGAAGCTTGTGGCTACAC<br>TGTTCCAAAGAATTCTCAAGTTTTCGTCAATGTCTGGGCTATTGGCA<br>GAGATCCAACTTTGTGGGAAGACCCATTGTCCTTCAAGCCAGAAAGA<br>TTCATGGCTTCCGAATTGGACGTTAGAGGTCGTGACTTCGAATTGTT<br>GCCATTCGGTGCTGGTAGAAGAATCTGTCCAGGTCTGCCATTGGCC<br>ATCAGAATGGTTCCAGTTATGTTGGGTTCTTTGGTTAACTCTTTCAAC<br>TGGAAACTAGATGGTGGTATTGCTCCAAAGGAATTGGACATGGGTG<br>AAAAATTTGGTATCACCTTAGCAAAGGCTCAATCTCTAAGAGCCGTC<br>CCATCTCCATTGGTGTGA |
| pCCW12-1X_RamO | Yeast RamR chimeric promoter. | This study. | CACCCATGAACCACACGGTTAGTCCAAAAGGGGCAGTTCAGATTCC<br>AGATGCGGGAATTAGCTTGCTGCCACCCTCACCTCACTAACGCTGC<br>GGTGTGCGGATACTTCATGCTATTTATAGACGCGCGTGTCGGAATCA<br>GCACGCGCAAGAACCAAATGGGAAAATCGGAATGGGTCCAGAACTG<br>CTTTGAGTGCTGGCTATTGGCGTCTGATTTCCGTTTTGGGAATCCTT<br>TGCCGCGCGCCCCTCTCAAAACTCCGCACAAGTCCCAGAAAGCGG<br>GAAAGAAATAAAACGCCACCAAAAAAAAAAAAATAAAAGCCAATCCT<br>CGAAGCGTGGGTGGTAGGCCCTGGATTATCCCGTACAAGTATTTCT |

| | | | |
|---|---|---|---|
| | | | CAGGAGTAAAAAAACCGTTTGTTTTGGAATTTCCCATTTCGCGGCCA<br>CCTACGCCGCTATCTTTGCAACAACTATCTGCGATAACTCAGCAAAT<br>TTTGCATATTCGTGTTGCAGTATTGCGATAATGGGAGTCTTACTTCCA<br>ACATAACGGCAGAAAGAAATGTGAGAAAATTTTGCATCCTTTGCCTC<br>CGTTCAAGTATATAAAGTCGGCATGCATGAGTGCTTACTCACTCATT<br>TGATAATCTTTCTTTCCATCCTACATTGTTCTAATTATTCTTATTCTCC<br>TTTATTCTTTCCTAACATACCAAGAAATTAATCTTCTGTCATTCGCTTA<br>AACACTATATCAATAAAGATCT |
| pCCW12-2X_RamO | Yeast RamR chimeric promoter. | This study. | CACCCATGAACCACACGGTTAGTCCAAAAGGGGCAGTTCAGATTCC<br>AGATGCGGGAATTAGCTTGCTGCCACCCTCACCTCACTAACGCTGC<br>GGTGTGCGGATACTTCATGCTATTTATAGACGCGCGTGTCGGAATCA<br>GCACGCGCAAGAACCAAATGGGAAAATCGGAATGGGTCCAGAACTG<br>CTTTGAGTGCTGGCTATTGGCGTCTGATTTCCGTTTTGGGAATCCTT<br>TGCCGCGCGCCCCTCTCAAAACTCCGCACAAGTCCCAGAAAGCGG<br>GAAAGAAATAAAACGCCACCAAAAAAAAAAAAATAAAAGCCAATCCT<br>CGAAGCGTGGGTGGTAGGCCCTGGATTATCCCGTACAAGTATTTCT<br>CAGGAGTAAAAAAACCGTTTGTTTTGGAATTTCCCATTTCGCGGCCA<br>CCTACGCCGCTATCTTTGCAACAACTATCTGCGATAACTCAGCAAAT<br>TTTGCATATTCGTGTTGCAGTATTGCGATAATGGGAGTCTTACTTCCA<br>ACATAACGGCAGAAAGAAATGTGAGAAAATTTTGCATCCTTTGCCTC<br>CGTTCAAGTATATAAAGTCGGCATGCATGAGTGCTTACTCACTCATA<br>TGAGTGCTTACTCACTCATTTGATAATCTTTCTTTCCATCCTACATTG<br>TTCTAATTATTCTTATTCTCCTTTATTCTTTCCTAACATACCAAGAAAT<br>TAATCTTCTGTCATTCGCTTAAACACTATATCAATAAAGATCT |
| pCCW12-mutRamO1 | Yeast RamR chimeric promoter. | This study. | CACCCATGAACCACACGGTTAGTCCAAAAGGGGCAGTTCAGATTCC<br>AGATGCGGGAATTAGCTTGCTGCCACCCTCACCTCACTAACGCTGC<br>GGTGTGCGGATACTTCATGCTATTTATAGACGCGCGTGTCGGAATCA<br>GCACGCGCAAGAACCAAATGGGAAAATCGGAATGGGTCCAGAACTG<br>CTTTGAGTGCTGGCTATTGGCGTCTGATTTCCGTTTTGGGAATCCTT<br>TGCCGCGCGCCCCTCTCAAAACTCCGCACAAGTCCCAGAAAGCGG<br>GAAAGAAATAAAACGCCACCAAAAAAAAAAAAATAAAAGCCAATCCT<br>CGAAGCGTGGGTGGTAGGCCCTGGATTATCCCGTACAAGTATTTCT<br>CAGGAGTAAAAAAACCGTTTGTTTTGGAATTTCCCATTTCGCGGCCA<br>CCTACGCCGCTATCTTTGCAACAACTATCTGCGATAACTCAGCAAAT<br>TTTGCATATTCGTGTTGCAGTATTGCGATAATGGGAGTCTTACTTCCA<br>ACATAACGGCAGAAAGAAATGTGAGAAAATTTTGCATCCTTTGCCTC<br>CGTTCAAGTATATAAAGTCGGCATGCTATAATGAGTGAGTACGCACT<br>CATTATTTGATAATCTTTCTTTCCATCCTACATTGTTCTAATTATTCTT<br>ATTCTCCTTTATTCTTTCCTAACATACCAAGAAATTAATCTTCTGTCAT<br>TCGCTTAAACACTATATCAATAAAGATCT |
| pCCW12-mutRamO2 | Yeast RamR chimeric promoter. | This study. | CACCCATGAACCACACGGTTAGTCCAAAAGGGGCAGTTCAGATTCC<br>AGATGCGGGAATTAGCTTGCTGCCACCCTCACCTCACTAACGCTGC<br>GGTGTGCGGATACTTCATGCTATTTATAGACGCGCGTGTCGGAATCA<br>GCACGCGCAAGAACCAAATGGGAAAATCGGAATGGGTCCAGAACTG<br>CTTTGAGTGCTGGCTATTGGCGTCTGATTTCCGTTTTGGGAATCCTT<br>TGCCGCGCGCCCCTCTCAAAACTCCGCACAAGTCCCAGAAAGCGG<br>GAAAGAAATAAAACGCCACCAAAAAAAAAAAAATAAAAGCCAATCCT<br>CGAAGCGTGGGTGGTAGGCCCTGGATTATCCCGTACAAGTATTTCT<br>CAGGAGTAAAAAAACCGTTTGTTTTGGAATTTCCCATTTCGCGGCCA<br>CCTACGCCGCTATCTTTGCAACAACTATCTGCGATAACTCAGCAAAT<br>TTTGCATATTCGTGTTGCAGTATTGCGATAATGGGAGTCTTACTTCCA<br>ACATAACGGCAGAAAGAAATGTGAGAAAATTTTGCATCCTTTGCCTC<br>CGTTCAAGTATATAAAGTCGGCATGCTTGATAATGAGTGAGTACGCA<br>CTCATTATAGATTGATAATCTTTCTTTCCATCCTACATTGTTCTAATTA<br>TTCTTATTCTCCTTTATTCTTTCCTAACATACCAAGAAATTAATCTTCT<br>GTCATTCGCTTAAACACTATATCAATAAAGATCT |

| | | | |
|---|---|---|---|
| pTDH3-1X_RamO | Yeast RamR chimeric promoter. | This study. | CTATTTTCGAGGACCTTGTCACCTTGAGCCCAAGAGAGCCAAGATTT<br>AAATTTTCCTATGACTTGATGCAAATTCCCAAAGCTAATAACATGCAA<br>GACACGTACGGTCAAGAAGACATATTTGACCTCTTAACAGGTTCAGA<br>CGCGACTGCCTCATCAGTAAGACCCGTTGAAAAGAACTTACCTGAAA<br>AAAACGAATATATACTAGCGTTGAATGTTAGCGTCAACAACAAGAAG<br>TTTAATGACGCGGAGGCCAAGGCAAAAAGATTCCTTGATTACGTAAG<br>GGAGTTAGAATCATTTTGAATAAAAAACACGCTTTTTCAGTTCGAGTT<br>TATCATTATCAATACTGCCATTTCAAAGAATACGTAAATAATTAATAGT<br>AGTGATTTTCCTAACTTTATTTAGTCAAAAAATTAGCCTTTTAATTCTG<br>CTGTAACCCGTACATGCCCAAAATAGGGGGCGGGTTACACAGAATA<br>TATAACATCGTAGGTGTCTGGGTGAACAGTTTATTCCTGGCATCCAC<br>TAAATATAATGGAGCCCGCTTTTTAAGCTGGCATCCAGAAAAAAAAA<br>GAATCCCAGCACCAAAATATTGTTTTCTTCACCAACCATCAGTTCATA<br>GGTCCATTCTCTTAGCGCAACTACAGAGAACAGGGGCACAAACAGG<br>CAAAAAACGGGCACAACCTCAATGGAGTGATGCAACCTGCCTGGAG<br>TAAATGATGACACAAGGCAATTGACCCACGCATGTATCTATCTCATTT<br>TCTTACACCTTCTATTACCTTCTGCTCTCTCTGATTTGGAAAAAGCTG<br>AAAAAAAAGGTTGAAACCAGTTCCCTGAAATTATTCCCCTACTTGACT<br>AATAAGTATATAAAGACGGTAATGAGTGCTTACTCACTCATGGTATTG<br>ATTGTAATTCTGTAAATCTATTTCTTAAACTTCTTAAATTCTACTTTTAT<br>AGTTAGTCTTTTTTTTAGTTTTAAAACACCAAGAACTTAGTTTCGAATA<br>AACACACATAAACAAACAAAACCTGCACTAAAACA |
| pTDH3-2X_RamO | Yeast RamR chimeric promoter. | This study. | CTATTTTCGAGGACCTTGTCACCTTGAGCCCAAGAGAGCCAAGATTT<br>AAATTTTCCTATGACTTGATGCAAATTCCCAAAGCTAATAACATGCAA<br>GACACGTACGGTCAAGAAGACATATTTGACCTCTTAACAGGTTCAGA<br>CGCGACTGCCTCATCAGTAAGACCCGTTGAAAAGAACTTACCTGAAA<br>AAAACGAATATATACTAGCGTTGAATGTTAGCGTCAACAACAAGAAG<br>TTTAATGACGCGGAGGCCAAGGCAAAAAGATTCCTTGATTACGTAAG<br>GGAGTTAGAATCATTTTGAATAAAAAACACGCTTTTTCAGTTCGAGTT<br>TATCATTATCAATACTGCCATTTCAAAGAATACGTAAATAATTAATAGT<br>AGTGATTTTCCTAACTTTATTTAGTCAAAAAATTAGCCTTTTAATTCTG<br>CTGTAACCCGTACATGCCCAAAATAGGGGGCGGGTTACACAGAATA<br>TATAACATCGTAGGTGTCTGGGTGAACAGTTTATTCCTGGCATCCAC<br>TAAATATAATGGAGCCCGCTTTTTAAGCTGGCATCCAGAAAAAAAAA<br>GAATCCCAGCACCAAAATATTGTTTTCTTCACCAACCATCAGTTCATA<br>GGTCCATTCTCTTAGCGCAACTACAGAGAACAGGGGCACAAACAGG<br>CAAAAAACGGGCACAACCTCAATGGAGTGATGCAACCTGCCTGGAG<br>TAAATGATGACACAAGGCAATTGACCCACGCATGTATCTATCTCATTT<br>TCTTACACCTTCTATTACCTTCTGCTCTCTCTGATTTGGAAAAAGCTG<br>AAAAAAAAGGTTGAAACCAGTTCCCTGAAATTATTCCCCTACTTGACT<br>AATAAGTATATAAAGACGGTAATGAGTGCTTACTCACTCATATGAGT<br>GCTTACTCACTCATGGTATTGATTGTAATTCTGTAAATCTATTTCTTAA<br>ACTTCTTAAATTCTACTTTTATAGTTAGTCTTTTTTTTAGTTTTAAAACA<br>CCAAGAACTTAGTTTCGAATAAACACACATAAACAAACAAAACCTGC<br>ACTAAAACA |
| pTDH3-mutRamO1 | Yeast RamR chimeric promoter. | This study. | CTATTTTCGAGGACCTTGTCACCTTGAGCCCAAGAGAGCCAAGATTT<br>AAATTTTCCTATGACTTGATGCAAATTCCCAAAGCTAATAACATGCAA<br>GACACGTACGGTCAAGAAGACATATTTGACCTCTTAACAGGTTCAGA<br>CGCGACTGCCTCATCAGTAAGACCCGTTGAAAAGAACTTACCTGAAA<br>AAAACGAATATATACTAGCGTTGAATGTTAGCGTCAACAACAAGAAG<br>TTTAATGACGCGGAGGCCAAGGCAAAAAGATTCCTTGATTACGTAAG<br>GGAGTTAGAATCATTTTGAATAAAAAACACGCTTTTTCAGTTCGAGTT<br>TATCATTATCAATACTGCCATTTCAAAGAATACGTAAATAATTAATAGT<br>AGTGATTTTCCTAACTTTATTTAGTCAAAAAATTAGCCTTTTAATTCTG<br>CTGTAACCCGTACATGCCCAAAATAGGGGGCGGGTTACACAGAATA<br>TATAACATCGTAGGTGTCTGGGTGAACAGTTTATTCCTGGCATCCAC<br>TAAATATAATGGAGCCCGCTTTTTAAGCTGGCATCCAGAAAAAAAAA<br>GAATCCCAGCACCAAAATATTGTTTTCTTCACCAACCATCAGTTCATA<br>GGTCCATTCTCTTAGCGCAACTACAGAGAACAGGGGCACAAACAGG |

| | | | |
|---|---|---|---|
| | | | CAAAAAACGGGCACAACCTCAATGGAGTGATGCAACCTGCCTGGAG<br>TAAATGATGACACAAGGCAATTGACCCACGCATGTATCTATCTCATTT<br>TCTTACACCTTCTATTACCTTCTGCTCTCTCTGATTTGGAAAAAGCTG<br>AAAAAAAAAGGTTGAAACCAGTTCCCTGAAATTATTCCCCTACTTGACT<br>AATAAGTATATAAAGACGGTATAATGAGTGAGTACGCACTCATTATG<br>GTATTGATTGTAATTCTGTAAATCTATTTCTTAAACTTCTTAAATTCTA<br>CTTTTATAGTTAGTCTTTTTTTTAGTTTTAAAACACCAAGAACTTAGTT<br>TCGAATAAACACACATAAACAAACAAAACCTGCACTAAAACA |
| pTDH3-mutRamO2 | Yeast RamR chimeric promoter. | This study. | CTATTTTCGAGGACCTTGTCACCTTGAGCCCAAGAGAGCCAAGATTT<br>AAATTTTCCTATGACTTGATGCAAATTCCCAAAGCTAATAACATGCAA<br>GACACGTACGGTCAAGAAGACATATTTGACCTCTTAACAGGTTCAGA<br>CGCGACTGCCTCATCAGTAAGACCCGTTGAAAAGAACTTACCTGAAA<br>AAAACGAATATATACTAGCGTTGAATGTTAGCGTCAACAACAAGAAG<br>TTTAATGACGCGGAGGCCAAGGCAAAAAGATTCCTTGATTACGTAAG<br>GGAGTTAGAATCATTTTGAATAAAAAACACGCTTTTTCAGTTCGAGTT<br>TATCATTATCAATACTGCCATTTCAAAGAATACGTAAATAATTAATAGT<br>AGTGATTTTCCTAACTTTATTTAGTCAAAAAATTAGCCTTTTAATTCTG<br>CTGTAACCCGTACATGCCCAAAATAGGGGGCGGGTTACACAGAATA<br>TATAACATCGTAGGTGTCTGGGTGAACAGTTTATTCCTGGCATCCAC<br>TAAATATAATGGAGCCCGCTTTTTAAGCTGGCATCCAGAAAAAAAAA<br>GAATCCCAGCACCAAAATATTGTTTTCTTCACCAACCATCAGTTCATA<br>GGTCCATTCTCTTAGCGCAACTACAGAGAACAGGGGCACAAACAGG<br>CAAAAAACGGGCACAACCTCAATGGAGTGATGCAACCTGCCTGGAG<br>TAAATGATGACACAAGGCAATTGACCCACGCATGTATCTATCTCATTT<br>TCTTACACCTTCTATTACCTTCTGCTCTCTCTGATTTGGAAAAAGCTG<br>AAAAAAAAAGGTTGAAACCAGTTCCCTGAAATTATTCCCCTACTTGACT<br>AATAAGTATATAAAGACGGTATTGATAATGAGTGAGTACGCACTCATT<br>ATAGAGGTATTGATTGTAATTCTGTAAATCTATTTCTTAAACTTCTTAA<br>ATTCTACTTTTATAGTTAGTCTTTTTTTTAGTTTTAAAACACCAAGAAC<br>TTAGTTTCGAATAAACACACATAAACAAACAAAACCTGCACTAAAACA |

**Supplementary Table 2. Plasmids cloned and used in this study**

| Name | Description | Reference |
|---|---|---|
| pReg-RamR | https://www.addgene.org/190610/ | (d'Oelsnitz et al., 2022) |
| Pramr-GFP | https://www.addgene.org/194148/ | (d'Oelsnitz et al., 2022) |
| pSELIS-RamR | https://www.addgene.org/202631/ | (d'Oelsnitz et al., 2022) |
| pReg-CAT1 | RamR repressor expression plasmid for CAT1 variant in *E. coli*. | This study. |
| pReg-CAT2 | RamR repressor expression plasmid for CAT2 variant in *E. coli*. | This study. |
| pReg-MIT1 | RamR repressor expression plasmid for MIT1 variant in *E. coli*. | This study. |
| pReg-MIT2 | RamR repressor expression plasmid for MIT2 variant in *E. coli*. | This study. |
| pReg-QUI1 | RamR repressor expression plasmid for QUI1 variant in *E. coli*. | This study. |
| pReg-RAU1 | RamR repressor expression plasmid for RAU1 variant in *E. coli*. | This study. |
| pReg-RAU2 | RamR repressor expression plasmid for RAU2 variant in *E. coli*. | This study. |
| pReg-STR1 | RamR repressor expression plasmid for STR1 variant in *E. coli*. | This study. |
| pET28-CAT2 | Expression plasmid for CAT2 variant with N-terminal His-tag for protein purification. | This study. |
| pET28-MIT2 | Expression plasmid for MIT2 variant with N-terminal His-tag for protein purification. | This study. |
| pET28-QUI1 | Expression plasmid for QUI1 variant with N-terminal His-tag for protein purification. | This study. |
| pET28-RAU2 | Expression plasmid for RAU2 variant with N-terminal His-tag for protein purification. | This study. |
| MoClo Yeast Toolkit (YTK) | https://www.addgene.org/kits/moclo-ytk/ | (Lee et al., 2015) |
| pCfB9340 | Guide RNA plasmid pgRNA_VIII-1_NatMX. | (Babaei et al., 2021) |
| pCfB9359 | VIII-1_Markerfree_BackBone for overexpression library integration. | (Babaei et al., 2021) |
| pCFB3045 | Guide RNA plasmid pgRNA_XI-3_NatMX. | (Jessop-Fabre et al., 2016) |
| pMHGG0_RamR_Sc_opt | Level-0 yeast MoClo plasmid containing yeast codon optimized RamR CDS part. | This study. |
| pMHGG0_RamR_nat | Level-0 yeast MoClo plasmid containing native codon usage RamR CDS part. | This study. |
| pMHGG1_RamR_Sc_opt | Level-1 yeast MoClo plasmid containing codon optimized RamR transcription unit (pCCW12-RamR-tADH1). Includes CEN-ARS yeast ORI and URA yeast selection marker. | This study. |
| pMHGG1_RamR_nat | Level-1 yeast MoClo plasmid containing native codon usage RamR transcription unit (pCCW12-RamR-tADH1). Includes CEN-ARS yeast ORI and URA yeast selection marker. | This study. |
| pMAD15 | Yeast genome integration plasmid of construct pTDH3-yeGFP-tCPS1 at site XI-3. | (Deichmann et al., 2025) |
| pMHO11 | Yeast genome integration plasmid of construct pTDH3-1X_RamO-yeGFP-tCPS1 at site XI-3. | This study. |
| pMHO12 | Yeast genome integration plasmid of construct pTDH3-2X_RamO-yeGFP-tCPS1 at site XI-3. | This study. |
| pMHO14 | Yeast genome integration plasmid of construct pTDH3-mutRamO1-yeGFP-tCPS1 at site XI-3. | This study. |

| | | |
|---|---|---|
| pMHO15 | Yeast genome integration plasmid of construct pTDH3-mutRamO2-yeGFP-tCPS1 at site XI-3. | This study. |
| pCCD2 | Yeast genome integration plasmid of construct pCCW12-yeGFP-tCPS1 at site XI-3. | This study. |
| pCCD3 | Yeast genome integration plasmid of construct pCCW12-1X_RamO-yeGFP-tCPS1 at site XI-3. | This study. |
| pCCD4 | Yeast genome integration plasmid of construct pCCW12-2X_RamO-yeGFP-tCPS1 at site XI-3. | This study. |
| pCCD5 | Yeast genome integration plasmid of construct pCCW12-mutRamO1-yeGFP-tCPS1 at site XI-3. | This study. |
| pCCD6 | Yeast genome integration plasmid of construct pCCW12-mutRamO2-yeGFP-tCPS1 at site XI-3. | This study. |
| pMHGG1_pCCW12-CAT2 | Level-1 yeast MoClo plasmid containing native codon usage RamR CAT2 transcription unit (pCCW12-CAT2-tADH1). Includes CEN-ARS yeast ORI and URA yeast selection marker. | This study. |
| pMHGG1_pHHF2-CAT2 | Level-1 yeast MoClo plasmid containing native codon usage RamR CAT2 transcription unit (pHHF2-CAT2-tADH1). Includes CEN-ARS yeast ORI and URA yeast selection marker. | This study. |
| pMHGG1_pRPL18B-CAT2 | Level-1 yeast MoClo plasmid containing native codon usage RamR CAT2 transcription unit (pRPL18B-CAT2-tADH1). Includes CEN-ARS yeast ORI and URA yeast selection marker. | This study. |
| pMHGG1_pCCW12-MIT2 | Level-1 yeast MoClo plasmid containing native codon usage RamR MIT2 transcription unit (pCCW12-MIT2-tADH1). Includes CEN-ARS yeast ORI and URA yeast selection marker. | This study. |
| pMHGG1_ pCCW12-QUI1 | Level-1 yeast MoClo plasmid containing native codon usage RamR QUI1 transcription unit (pCCW12-QUI1-tADH1). Includes CEN-ARS yeast ORI and URA yeast selection marker. | This study. |
| pMHGG1_pHHF2-QUI1 | Level-1 yeast MoClo plasmid containing native codon usage RamR MIT2 transcription unit (pHHF2-QUI1-tADH1). Includes CEN-ARS yeast ORI and URA yeast selection marker. | This study. |
| pMHGG1_pRPL18B-QUI1 | Level-1 yeast MoClo plasmid containing native codon usage RamR MIT2 transcription unit (pRPL18B-QUI1-tADH1). Includes CEN-ARS yeast ORI and URA yeast selection marker. | This study. |
| pMHGG1_pCCW12-RAU2 | Level-1 yeast MoClo plasmid containing native codon usage RamR RAU2 transcription unit (pCCW12-RAU2-tADH1). Includes CEN-ARS yeast ORI and URA yeast selection marker. | This study. |
| pMHGG1_pHHF2-RAU2 | Level-1 yeast MoClo plasmid containing native codon usage RamR RAU2 transcription unit (pHHF2-RAU2-tADH1). Includes CEN-ARS yeast ORI and URA yeast selection marker. | This study. |
| pMHGG1_pRPL18B-RAU2 | Level-1 yeast MoClo plasmid containing native codon usage RamR RAU2 transcription unit (pRPL18B-RAU2-tADH1). Includes CEN-ARS yeast ORI and URA yeast selection marker. | This study. |
| pMHGG1_pCCW12-STR1 | Level-1 yeast MoClo plasmid containing native codon usage RamR STR1 transcription unit (pCCW12-STR1-tADH1). Includes CEN-ARS yeast ORI and URA yeast selection marker. | This study. |

| | | |
|---|---|---|
| pMHO32 | Yeast genome integration plasmid of reporter and repressor construct (pCCW12-1X_RamO-yeGFP-tCPS1 + pCCW12-RAU2-tADH1) at site VIII-1. | This study. |
| pMHO33 | Yeast genome integration plasmid of reporter and repressor construct (pCCW12-1X_RamO-yeGFP-tCPS1 + pCCW12-CAT2-tADH1) at site VIII-1. | This study. |
| pMHO34 | Yeast genome integration plasmid of reporter and repressor construct (pCCW12-1X_RamO-yeGFP-tCPS1 + pCCW12-STR1-tADH1) at site VIII-1. | This study. |
| pMHO37 | Yeast genome integration plasmid of reporter and repressor construct (pCCW12-1X_RamO-yeGFP-tCPS1 + pCCW12-RAU2-NLS-tADH1) at site VIII-1. | This study. |
| pMHO38 | Yeast genome integration plasmid of reporter and repressor construct (pCCW12-1X_RamO-yeGFP-tCPS1 + pCCW12-CAT2-NLS-tADH1) at site VIII-1. | This study. |
| pMHO39 | Yeast genome integration plasmid of reporter and repressor construct (pCCW12-1X_RamO-yeGFP-tCPS1 + pCCW12-STR1-NLS-tADH1) at site VIII-1. | This study. |

**Supplementary Table 3. Yeast strains used in this study.** Bold font in the genotype indicates the genetic modification made to the parent strain.

| Name | Parent | Genotype | Reference |
|---|---|---|---|
| MIA-B0 | CEN.PK2-1C | MATa; his3D1; leu2-3_112; ura3-52; trp1-289; PTEF1-SpCas9-TCYC1. | (J. Zhang et al., 2022) |
| yMHO20 | MIA-B0 | MATa; his3D1; leu2-3_112; ura3-52; trp1-289; PTEF1-SpCas9-TCYC1, **PTDH3-yeGFP-TCPS1**. | This study. |
| yMHO21 | MIA-B0 | MATa; his3D1; leu2-3_112; ura3-52; trp1-289; PTEF1-SpCas9-TCYC1, **PTDH3_1XRamO-yeGFP-TCPS1**. | This study. |
| yMHO22 | MIA-B0 | MATa; his3D1; leu2-3_112; ura3-52; trp1-289; PTEF1-SpCas9-TCYC1, **PTDH3_2XRamO-yeGFP-TCPS1**. | This study. |
| yMHO32 | MIA-B0 | MATa; his3D1; leu2-3_112; ura3-52; trp1-289; PTEF1-SpCas9-TCYC1, **PTDH3_mutRamO1-yeGFP-TCPS1**. | This study. |
| yMHO33 | MIA-B0 | MATa; his3D1; leu2-3_112; ura3-52; trp1-289; PTEF1-SpCas9-TCYC1, **PCCW12_mutRamO2-yeGFP-TCPS1**. | This study. |
| yCCD6 | MIA-B0 | MATa; his3D1; leu2-3_112; ura3-52; trp1-289; PTEF1-SpCas9-TCYC1, **PCCW12-yeGFP-TCPS1**. | This study. |
| yCCD7 | MIA-B0 | MATa; his3D1; leu2-3_112; ura3-52; trp1-289; PTEF1-SpCas9-TCYC1, **PCCW12_1XRamO-yeGFP-TCPS1**. | This study. |
| yCCD8 | MIA-B0 | MATa; his3D1; leu2-3_112; ura3-52; trp1-289; PTEF1-SpCas9-TCYC1, **PCCW12_2XRamO-yeGFP-TCPS1**. | This study. |
| yCCD9 | MIA-B0 | MATa; his3D1; leu2-3_112; ura3-52; trp1-289; PTEF1-SpCas9-TCYC1, **PCCW12_mutRamO1-yeGFP-TCPS1**. | This study. |
| yCCD10 | MIA-B0 | MATa; his3D1; leu2-3_112; ura3-52; trp1-289; PTEF1-SpCas9-TCYC1, **PCCW12_mutRamO2-yeGFP-TCPS1**. | This study. |
| yCCD1 | yMHO21 | MATa; his3D1; leu2-3_112; ura3-52; trp1-289; PTEF1-SpCas9-TCYC1, PTDH3_1XRamO-yeGFP-TCPS1, **pMHGG1_RamR_nat**. | This study. |
| yCCD2 | yMHO22 | MATa; his3D1; leu2-3_112; ura3-52; trp1-289; PTEF1-SpCas9-TCYC1, PTDH3_2XRamO-yeGFP-TCPS1, **pMHGG1_RamR_nat**. | This study. |
| yCCD3 | yMHO32 | MATa; his3D1; leu2-3_112; ura3-52; trp1-289; PTEF1-SpCas9-TCYC1, PTDH3_mutRamO1-yeGFP-TCPS1, **pMHGG1_RamR_nat**. | This study. |
| yCCD4 | yMHO33 | MATa; his3D1; leu2-3_112; ura3-52; trp1-289; PTEF1-SpCas9-TCYC1, PCCW12_mutRamO2-yeGFP-TCPS1, **pMHGG1_RamR_nat**. | This study. |
| yCCD12 | yCCD7 | MATa; his3D1; leu2-3_112; ura3-52; trp1-289; PTEF1-SpCas9-TCYC1, PCCW12_1XRamO-yeGFP-TCPS1, **pMHGG1_RamR_nat**. | This study. |
| yCCD13 | yCCD8 | MATa; his3D1; leu2-3_112; ura3-52; trp1-289; PTEF1-SpCas9-TCYC1, PCCW12_2XRamO-yeGFP-TCPS1, **pMHGG1_RamR_nat**. | This study. |
| yCCD14 | yCCD9 | MATa; his3D1; leu2-3_112; ura3-52; trp1-289; PTEF1-SpCas9-TCYC1, PCCW12_mutRamO1-yeGFP-TCPS1, **pMHGG1_RamR_nat**. | This study. |
| yCCD15 | yCCD10 | MATa; his3D1; leu2-3_112; ura3-52; trp1-289; PTEF1-SpCas9-TCYC1, PCCW12_mutRamO2-yeGFP-TCPS1, **pMHGG1_RamR_nat**. | This study. |
| yCCD23 | MIA-B0 | MATa; his3D1; leu2-3_112; ura3-52; trp1-289; PTEF1-SpCas9-TCYC1, **PCCW12_1XRamO-yeGFP-TCPS1**. | This study. |
| yCCD20 | yCCD7 | MATa; his3D1; leu2-3_112; ura3-52; trp1-289; PTEF1-SpCas9-TCYC1, PCCW12_1XRamO-yeGFP-TCPS1, **pMHGG1_ pCCW12-QUI1**. | This study. |
| yCCD28 | yCCD7 | MATa; his3D1; leu2-3_112; ura3-52; trp1-289; PTEF1-SpCas9-TCYC1, PCCW12_1XRamO-yeGFP-TCPS1, **pMHGG1_ pCCW12-STR1**. | This study. |

| | | | |
|---|---|---|---|
| yCCD46 | yCCD7 | MATa; his3D1; leu2-3_112; ura3-52; trp1-289; PTEF1-SpCas9-TCYC1, PCCW12_1XRamO-yeGFP-TCPS1, **pMHGG1_ pCCW12-MIT2**. | This study. |
| yCCD49 | yCCD7 | MATa; his3D1; leu2-3_112; ura3-52; trp1-289; PTEF1-SpCas9-TCYC1, PCCW12_1XRamO-yeGFP-TCPS1, **pMHGG1_ pCCW12-CAT2**. | This study. |
| yCCD50 | yCCD7 | MATa; his3D1; leu2-3_112; ura3-52; trp1-289; PTEF1-SpCas9-TCYC1, PCCW12_1XRamO-yeGFP-TCPS1, **pMHGG1_ pCCW12-RAU2**. | This study. |
| yCCD51 | yCCD7 | MATa; his3D1; leu2-3_112; ura3-52; trp1-289; PTEF1-SpCas9-TCYC1, PCCW12_1XRamO-yeGFP-TCPS1, **pMHGG1_ pHHF2-QUI1**. | This study. |
| yCCD52 | yCCD7 | MATa; his3D1; leu2-3_112; ura3-52; trp1-289; PTEF1-SpCas9-TCYC1, PCCW12_1XRamO-yeGFP-TCPS1, **pMHGG1_ pRPL18B-QUI1**. | This study. |
| yMHO75 | yCCD7 | MATa; his3D1; leu2-3_112; ura3-52; trp1-289; PTEF1-SpCas9-TCYC1, PCCW12_1XRamO-yeGFP-TCPS1, **pMHGG1_ pHHF2-RAU2**. | This study. |
| yMHO76 | yCCD7 | MATa; his3D1; leu2-3_112; ura3-52; trp1-289; PTEF1-SpCas9-TCYC1, PCCW12_1XRamO-yeGFP-TCPS1, **pMHGG1_ pRPL18B-RAU2**. | This study. |
| yMHO78 | yCCD7 | MATa; his3D1; leu2-3_112; ura3-52; trp1-289; PTEF1-SpCas9-TCYC1, PCCW12_1XRamO-yeGFP-TCPS1, **pMHGG1_ pHHF2-CAT2**. | This study. |
| yMHO79 | yCCD7 | MATa; his3D1; leu2-3_112; ura3-52; trp1-289; PTEF1-SpCas9-TCYC1, PCCW12_1XRamO-yeGFP-TCPS1, **pMHGG1_ pRPL18B-CAT2**. | This study. |
| yMHO87 | MIA-B0 | MATa; his3D1; leu2-3_112; ura3-52; trp1-289; PTEF1-SpCas9-TCYC1, **PCCW12_1XRamO-yeGFP-TCPS1, PCCW12-RAU2-TADH1**. | This study. |
| yMHO88 | MIA-B0 | MATa; his3D1; leu2-3_112; ura3-52; trp1-289; PTEF1-SpCas9-TCYC1, **PCCW12_1XRamO-yeGFP-TCPS1, PCCW12-CAT2-TADH1**. | This study. |
| yMHO89 | MIA-B0 | MATa; his3D1; leu2-3_112; ura3-52; trp1-289; PTEF1-SpCas9-TCYC1, **PCCW12_1XRamO-yeGFP-TCPS1, PCCW12-STR1-TADH1**. | This study. |
| yMHO92 | MIA-B0 | MATa; his3D1; leu2-3_112; ura3-52; trp1-289; PTEF1-SpCas9-TCYC1, **PCCW12_1XRamO-yeGFP-TCPS1, PCCW12-RAU2_NLS-TADH1**. | This study. |
| yMHO93 | MIA-B0 | MATa; his3D1; leu2-3_112; ura3-52; trp1-289; PTEF1-SpCas9-TCYC1, **PCCW12_1XRamO-yeGFP-TCPS1, PCCW12-CAT2_NLS-TADH1**. | This study. |
| yMHO94 | MIA-B0 | MATa; his3D1; leu2-3_112; ura3-52; trp1-289; PTEF1-SpCas9-TCYC1, **PCCW12_1XRamO-yeGFP-TCPS1, PCCW12-STR1_NLS-TADH1.** | This study. |
| MIA-CZ1 | MIA-CM-5 | MATa, his3D1, leu2-3_112, ura3-52, trp1-289, atf1Δ oye2Δ adh6Δ oye3Δ ari1Δ, rox1Δ, PTEF1-SpyCas9-TCYC1, PTEF1-CroCPR-TPRM9, PPGK1-CroCYB5-TIDP1, PTEF2-AgrGPPS2-TVPS13, PFBA1-GgaFPSN144W-TIDP1, PPGK1-IDI1-TPRM9, PTDH3-tHMG1-TADH1, PCCW12-ERG20F96W, N127WtCroGES-TCYC1, PPGK1-CroTDC-TPRM9, PTDH3-CroG8H-TADH1, PPGK1-Vmi8HGO-A -TADH1,PFBA1-NcaISY-TCYC1, PTEF1-NcaMLPLA-TADH1, PTEF2-CroIO-TCYC1, PFBA1-CroADH2-TCPS1,PPGK1-Cro7DLGT-TVPS13, PTEF2-Cro7DLH-TCYC1, PTDH3-CroLAMT-TADH1, PTPI1-CroSLS-TIDP1,PFBA1-CroSTR-TPRM9, PCCW12-ZWF1-TADH1, PPGK1-INO2-TCYC1. | (Holtz et al., 2024) |
| yMHO130 | MIA-CZ-1 | MATa, his3D1, leu2-3_112, ura3-52, trp1-289, atf1Δ oye2Δ adh6Δ oye3Δ ari1Δ, rox1Δ, PTEF1-SpyCas9-TCYC1, PTEF1-CroCPR- | This study. |

| | | | |
|---|---|---|---|
| | | TPRM9, PPGK1-CroCYB5-TIDP1, PTEF2-AgrGPPS2-TVPS13, PFBA1-GgaFPSN144W-TIDP1, PPGK1-IDI1-TPRM9, PTDH3-tHMG1-TADH1, PCCW12-ERG20F96W, N127WtCroGES-TCYC1, PPGK1-CroTDC-TPRM9, **PTDH3-TeG8H-TADH1**, PPGK1-Vmi8HGO-A -TADH1,PFBA1-NcaISY-TCYC1, PTEF1-NcaMLPLA-TADH1, PTEF2-CroIO-TCYC1, PFBA1-CroADH2-TCPS1,PPGK1-Cro7DLGT-TVPS13, PTEF2-Cro7DLH-TCYC1, PTDH3-CroLAMT-TADH1, PTPI1-CroSLS-TIDP1,PFBA1-CroSTR-TPRM9, PCCW12-ZWF1-TADH1, PPGK1-INO2-TCYC1. | |
| yMHO131 | MIA-CZ-1 | MATa, his3D1, leu2-3_112, ura3-52, trp1-289, atf1Δ oye2Δ adh6Δ oye3Δ ari1Δ, rox1Δ, PTEF1-SpyCas9-TCYC1, PTEF1-CroCPR-TPRM9, PPGK1-CroCYB5-TIDP1, PTEF2-AgrGPPS2-TVPS13, PFBA1-GgaFPSN144W-TIDP1, PPGK1-IDI1-TPRM9, PTDH3-tHMG1-TADH1, PCCW12-ERG20F96W, N127WtCroGES-TCYC1, PPGK1-CroTDC-TPRM9, **PTDH3-RsG8H-TADH1**, PPGK1-Vmi8HGO-A -TADH1,PFBA1-NcaISY-TCYC1, PTEF1-NcaMLPLA-TADH1, PTEF2-CroIO-TCYC1, PFBA1-CroADH2-TCPS1,PPGK1-Cro7DLGT-TVPS13, PTEF2-Cro7DLH-TCYC1, PTDH3-CroLAMT-TADH1, PTPI1-CroSLS-TIDP1,PFBA1-CroSTR-TPRM9, PCCW12-ZWF1-TADH1, PPGK1-INO2-TCYC1. | This study. |
| yMHO132 | MIA-CZ-1 | MATa, his3D1, leu2-3_112, ura3-52, trp1-289, atf1Δ oye2Δ adh6Δ oye3Δ ari1Δ, rox1Δ, PTEF1-SpyCas9-TCYC1, PTEF1-CroCPR-TPRM9, PPGK1-CroCYB5-TIDP1, PTEF2-AgrGPPS2-TVPS13, PFBA1-GgaFPSN144W-TIDP1, PPGK1-IDI1-TPRM9, PTDH3-tHMG1-TADH1, PCCW12-ERG20F96W, N127WtCroGES-TCYC1, PPGK1-CroTDC-TPRM9, **PTDH3-LjG8H-TADH1**, PPGK1-Vmi8HGO-A -TADH1,PFBA1-NcaISY-TCYC1, PTEF1-NcaMLPLA-TADH1, PTEF2-CroIO-TCYC1, PFBA1-CroADH2-TCPS1,PPGK1-Cro7DLGT-TVPS13, PTEF2-Cro7DLH-TCYC1, PTDH3-CroLAMT-TADH1, PTPI1-CroSLS-TIDP1,PFBA1-CroSTR-TPRM9, PCCW12-ZWF1-TADH1, PPGK1-INO2-TCYC1. | This study. |
| yMHO133 | MIA-CZ-1 | MATa, his3D1, leu2-3_112, ura3-52, trp1-289, atf1Δ oye2Δ adh6Δ oye3Δ ari1Δ, rox1Δ, PTEF1-SpyCas9-TCYC1, PTEF1-CroCPR-TPRM9, PPGK1-CroCYB5-TIDP1, PTEF2-AgrGPPS2-TVPS13, PFBA1-GgaFPSN144W-TIDP1, PPGK1-IDI1-TPRM9, PTDH3-tHMG1-TADH1, PCCW12-ERG20F96W, N127WtCroGES-TCYC1, PPGK1-CroTDC-TPRM9, **PTDH3-tr24LjG8H-TADH1**, PPGK1-Vmi8HGO-A -TADH1,PFBA1-NcaISY-TCYC1, PTEF1-NcaMLPLA-TADH1, PTEF2-CroIO-TCYC1, PFBA1-CroADH2-TCPS1,PPGK1-Cro7DLGT-TVPS13, PTEF2-Cro7DLH-TCYC1, PTDH3-CroLAMT-TADH1, PTPI1-CroSLS-TIDP1,PFBA1-CroSTR-TPRM9, PCCW12-ZWF1-TADH1, PPGK1-INO2-TCYC1. | This study. |
| yMHO145 | yMHO130 | MATa, his3D1, leu2-3_112, ura3-52, trp1-289, atf1Δ oye2Δ adh6Δ oye3Δ ari1Δ, rox1Δ, PTEF1-SpyCas9-TCYC1, PTEF1-CroCPR-TPRM9, PPGK1-CroCYB5-TIDP1, PTEF2-AgrGPPS2-TVPS13, PFBA1-GgaFPSN144W-TIDP1, PPGK1-IDI1-TPRM9, PTDH3-tHMG1-TADH1, PCCW12-ERG20F96W, N127WtCroGES-TCYC1, PPGK1-CroTDC-TPRM9, PTDH3-TeG8H-TADH1, PPGK1-Vmi8HGO-A -TADH1,PFBA1-NcaISY-TCYC1, PTEF1-NcaMLPLA-TADH1, PTEF2-CroIO-TCYC1, PFBA1-CroADH2-TCPS1,PPGK1-Cro7DLGT-TVPS13, PTEF2-Cro7DLH-TCYC1, PTDH3-CroLAMT-TADH1, PTPI1-CroSLS-TIDP1,PFBA1-CroSTR-TPRM9, PCCW12-ZWF1-TADH1, PPGK1-INO2-TCYC1, **PCCW12_1XRamO-yeGFP-TCPS1, PCCW12-STR1_NLS-TADH1.** | This study. |
| yMHO147 | yMHO145 | MATa, his3D1, leu2-3_112, ura3-52, trp1-289, atf1Δ oye2Δ adh6Δ oye3Δ ari1Δ, rox1Δ, PTEF1-SpyCas9-TCYC1, PTEF1-CroCPR-TPRM9, PPGK1-CroCYB5-TIDP1, PTEF2-AgrGPPS2-TVPS13, PFBA1-GgaFPSN144W-TIDP1, PPGK1-IDI1-TPRM9, PTDH3-tHMG1-TADH1, PCCW12-ERG20F96W, N127WtCroGES-TCYC1, | This study. |

| | | | |
|---|---|---|---|
| | | PPGK1-CroTDC-TPRM9, PTDH3-TeG8H-TADH1, PPGK1-Vmi8HGO-A -TADH1,PFBA1-NcaISY-TCYC1, PTEF1-NcaMLPLA-TADH1, PTEF2-CroIO-TCYC1, PFBA1-CroADH2-TCPS1,PPGK1-Cro7DLGT-TVPS13, PTEF2-Cro7DLH-TCYC1, PTDH3-CroLAMT-TADH1, PFBA1-CroSTR-TPRM9, PCCW12-ZWF1-TADH1, PPGK1-INO2-TCYC1, PCCW12_1XRamO-yeGFP-TCPS1, PCCW12-STR1_NLS-TADH1 (**ΔCroSLS**). | |
| yMHO212 | yMHO130 | MATa, his3D1, leu2-3_112, ura3-52, trp1-289, atf1Δ oye2Δ adh6Δ oye3Δ ari1Δ, rox1Δ, PTEF1-SpyCas9-TCYC1, PTEF1-CroCPR-TPRM9, PPGK1-CroCYB5-TIDP1, PTEF2-AgrGPPS2-TVPS13, PFBA1-GgaFPSN144W-TIDP1, PPGK1-IDI1-TPRM9, PTDH3-tHMG1-TADH1, PCCW12-ERG20F96W, N127WtCroGES-TCYC1, PPGK1-CroTDC-TPRM9, PTDH3-TeG8H-TADH1, PPGK1-Vmi8HGO-A -TADH1,PFBA1-NcaISY-TCYC1, PTEF1-NcaMLPLA-TADH1, PTEF2-CroIO-TCYC1, PFBA1-CroADH2-TCPS1,PPGK1-Cro7DLGT-TVPS13, PTEF2-Cro7DLH-TCYC1, PTDH3-CroLAMT-TADH1, PFBA1-CroSTR-TPRM9, PCCW12-ZWF1-TADH1, PPGK1-INO2-TCYC1 (**ΔCroSLS**). | This study. |
| yMHO242 | yMHO130 | MATa, his3D1, leu2-3_112, ura3-52, trp1-289, atf1Δ oye2Δ adh6Δ oye3Δ ari1Δ, rox1Δ, PTEF1-SpyCas9-TCYC1, PTEF1-CroCPR-TPRM9, PPGK1-CroCYB5-TIDP1, PTEF2-AgrGPPS2-TVPS13, PFBA1-GgaFPSN144W-TIDP1, PPGK1-IDI1-TPRM9, PTDH3-tHMG1-TADH1, PCCW12-ERG20F96W, N127WtCroGES-TCYC1, PPGK1-CroTDC-TPRM9, PTDH3-TeG8H-TADH1, PPGK1-Vmi8HGO-A -TADH1,PFBA1-NcaISY-TCYC1, PTEF1-NcaMLPLA-TADH1, PTEF2-CroIO-TCYC1, PFBA1-CroADH2-TCPS1,PPGK1-Cro7DLGT-TVPS13, PTEF2-Cro7DLH-TCYC1, PTDH3-CroLAMT-TADH1, PTPI1-CroSLS-TIDP1,PFBA1-CroSTR-TPRM9, PCCW12-ZWF1-TADH1, PPGK1-INO2-TCYC1, **PTDH3-PHB1-TENO2, PCCW12-PHB2-TTDH1**. | This study. |
| yMHO243 | yMHO130 | MATa, his3D1, leu2-3_112, ura3-52, trp1-289, atf1Δ oye2Δ adh6Δ oye3Δ ari1Δ, rox1Δ, PTEF1-SpyCas9-TCYC1, PTEF1-CroCPR-TPRM9, PPGK1-CroCYB5-TIDP1, PTEF2-AgrGPPS2-TVPS13, PFBA1-GgaFPSN144W-TIDP1, PPGK1-IDI1-TPRM9, PTDH3-tHMG1-TADH1, PCCW12-ERG20F96W, N127WtCroGES-TCYC1, PPGK1-CroTDC-TPRM9, PTDH3-TeG8H-TADH1, PPGK1-Vmi8HGO-A -TADH1,PFBA1-NcaISY-TCYC1, PTEF1-NcaMLPLA-TADH1, PTEF2-CroIO-TCYC1, PFBA1-CroADH2-TCPS1,PPGK1-Cro7DLGT-TVPS13, PTEF2-Cro7DLH-TCYC1, PTDH3-CroLAMT-TADH1, PTPI1-CroSLS-TIDP1,PFBA1-CroSTR-TPRM9, PCCW12-ZWF1-TADH1, PPGK1-INO2-TCYC1, **PTDH3-PHB1-TENO2, PCCW12-PHB2-TTDH1, PTEF1-PSD1-TPGK1.** | This study. |

**Supplementary Table 4. Primers used for RamR library construction.** Degenerate NNS codons are shown in bold.

| Name | Sequence |
|---|---|
| Blue_lib_Fw | AACACCCTTTACTTACATTTG**NNS**CAGGAC**NNS**TGCCAATCA**NNS**ATCATGGAATTGGATCGTTCTATTACTGACGC |
| Blue_lib_Rv | CAAATGTAAGTAAAGGGTGTTGATCAACTCATCTTTCG |
| Yellow_lib_Fw | TTCTATTACTGACGCTAAGATGATG**NNS**CGTTTT**NNS**TGGAACAGT**NNS**ATTAGCTGGGGATTGAACCACCCAG |
| Yellow_lib_Rv | CATCATCTTAGCGTCAGTAATAGAACGATCCAATTC |
| Red_lib_Fw | GTGTTTATGTCCGACGAGTAC**NNS**GCCTTCGGC**NNS**GGGTTGTTC**NNS**GCGCTTGCTGAGACGACTATG |
| Red_lib_Rv | GTACTCGTCGGACATAAACACCATAAGAAC |
| Pink_lib_Fw | GATATGTTCCCGGAGTTACGCGAC |
| Pink_lib_Rv | CGTAACTCCGGGAACATATC**SNNSNN**GCGTTG**SNN**GGTTTCCTTCGTCAACTTTTCAGAA |
| Purple_lib_Fw | GTTCTTATGGTGTTTATGTCCGACGAGTAC |
| Purple_lib_Rv | TCGGACATAAACACCATAAGAAC**SNN**ACGGTG**SNNSNN**GTCGCGTAACTCCGGGAAC |
| Green_lib_Fw | TTGCGTTGGGCTTCGAGGCT**NNS**TGGCGCGCA**NNSNNS**CGCGAAGAGCAGTAATAATCCTAAG |
| Green_lib_Rv | AGCCTCGAAGCCCAACGCAATG |
| Lilac_lib_Fw | CCACCGTAGTGTTCTTATGGTG**NNSNNS**TCCGACGAGTACCGCGCCTTC**NNS**GACGGGTTGTTCTTGGCG |
| Lilac_lib_Rv | CACCATAAGAACACTACGGTGGCACAAGT |
| Silver_lib_Fw | GAACCACCCAGCTCGCCATCGTGCC**NNSNNS**CAGTTG**NNS**GTTTCTGAAAAGTTGACGAAGGAAAC |
| Silver_lib_Rv | GATGGCGAGCTGGGTGGTTCAATC |
| Orange_lib_Fw | TCCCGGAGTTACGCGACTTG**NNS**CACCGT**NNSNNS**CTTATGGTGTTTATGTCCGACGAG |

| | |
|---|---|
| Orange_lib_Rv | CAAGTCGCGTAACTCCGGGAACATATC |
| epPCR_insert_Fw | CGGAACCACGTATCAGAAGGAGGTTAGTATATG |
| epPCR_insert_Rv | CGAGCCTGCACTCCTCGAATTCTTAGGATTATTA |
| epPCR_backbone_Fw | TAATAATCCTAAGAATTCGAGGAGTGCAGGCTCG |
| epPCR_backbone_Rv | CATATACTAACCTCCTTCTGATACGTGGTTCCG |

**Supplementary Table 5. Overview of genes included in the overexpression library tested in yMHO130 to demonstrate an application of the STR1 sensor.** Endogenous genes from *S. cerevisiae* are colored in blue, their putative homologs from *Y. lipolytica* are colored in pink while plant genes are colored in green. The full FASTA file containing the DNA sequences of each gene of the library is available in **Supp. File 1**.

| Gene | Function | Category |
|---|---|---|
| DAP1 | Heme-binding protein regulating sterol synthesis | Lipid metabolism |
| INO2 | Transcription factor for phospholipid genes | Transcriptional regulation |
| ICE2 | ER membrane protein in lipid homeostasis | Lipid metabolism |
| ERO1 | Disulfide bond formation in ER | Protein folding (ER oxidative folding) |
| CPR5 | Peptidyl-prolyl isomerase | Chaperone |
| HSP26 | Small heat shock protein | Chaperone |
| SSA1 | Cytosolic Hsp70 | Chaperone |
| HSP104 | Protein disaggregase | Chaperone |
| HAC1 | UPR transcription factor | Transcriptional regulation |
| UPC2 | Sterol-regulated transcription factor | Transcriptional regulation |
| POS5 | Mitochondrial NADH kinase (NADPH) | Cofactor & redox metabolism |
| ZWF1 | Glucose-6-phosphate dehydrogenase | Cofactor & redox metabolism |
| STB5 | Regulates NADPH-related genes | Transcriptional regulation |
| ALD6 | Aldehyde dehydrogenase (NADPH production) | Cofactor & redox metabolism |
| TPI1 | Glycolysis enzyme | Central carbon metabolism |
| OLE1 | Fatty acid desaturase | Lipid metabolism |
| HSP78 | Mitochondrial disaggregase | Chaperone |
| HEM13 | Coproporphyrinogen oxidase | Heme biosynthesis |
| HEM14 | Protoporphyrinogen oxidase | Heme biosynthesis |
| HEM2 | ALA dehydratase | Heme biosynthesis |
| HEM15 | Ferrochelatase | Heme biosynthesis |
| HEM3 | Porphobilinogen deaminase | Heme biosynthesis |
| HEM12 | Uroporphyrinogen decarboxylase | Heme biosynthesis |
| FET4 | Iron transporter | Transport |
| DGK1 | Diacylglycerol kinase | Lipid metabolism |
| CDS1 | CDP-diacylglycerol synthase | Lipid metabolism |
| CHO1 | Phosphatidylserine synthase | Lipid metabolism |
| PSD1 | PS decarboxylase (mitochondrial) | Lipid metabolism |

| | | |
|---|---|---|
| PSD2 | PS decarboxylase (Golgi/vacuole) | Lipid metabolism |
| CHO2 | Phospholipid methyltransferase | Lipid metabolism |
| OPI3 | Phospholipid methyltransferase | Lipid metabolism |
| SCT1 | Glycerol-3-phosphate acyltransferase | Lipid metabolism |
| GPT2 | Glycerol-3-phosphate acyltransferase | Lipid metabolism |
| AYR1 | Acyl-DHAP reductase | Lipid metabolism |
| SLC1 | Acyltransferase | Lipid metabolism |
| ALE1 | Lysophospholipid acyltransferase | Lipid metabolism |
| URA7 | CTP synthase | Nucleotide metabolism |
| URA8 | CTP synthase | Nucleotide metabolism |
| CKI1 | Choline kinase | Lipid metabolism |
| PCT1 | Cytidylyltransferase | Lipid metabolism |
| CPT1 | Cholinephosphotransferase | Lipid metabolism |
| CPR1 | Cyclophilin | Chaperone |
| YDR154C | Uncharacterized protein | Unknown |
| CPR3 | Mitochondrial cyclophilin | Chaperone |
| CPR2 | ER cyclophilin | Chaperone |
| CPR6 | Hsp90 co-chaperone | Chaperone |
| CPR4 | Secreted cyclophilin | Chaperone |
| SSA2 | Cytosolic Hsp70 | Chaperone |
| SSA4 | Stress-induced Hsp70 | Chaperone |
| SSA3 | Stress-induced Hsp70 | Chaperone |
| SSB2 | Ribosome-associated Hsp70 | Chaperone |
| SSB1 | Ribosome-associated Hsp70 | Chaperone |
| KAR2 | ER Hsp70 (BiP) | Chaperone |
| SSC1 | Mitochondrial Hsp70 | Chaperone |
| ECM10 | Hsp70-like protein | Chaperone |
| SSQ1 | Fe-S cluster assembly Hsp70 | Fe-S cluster biogenesis |
| ERG10 | Acetyl-CoA acetyltransferase | Mevalonate pathway |
| ERG13 | HMG-CoA synthase | Mevalonate pathway |
| HMG1 | HMG-CoA reductase | Mevalonate pathway |
| ERG12 | Mevalonate kinase | Mevalonate pathway |
| ERG8 | Phosphomevalonate kinase | Mevalonate pathway |
| MVD1 | Mevalonate diphosphate decarboxylase | Mevalonate pathway |
| IDI1 | IPP isomerase | Mevalonate pathway |
| ERG20 | Farnesyl diphosphate synthase | Mevalonate pathway |
| ATP1 | α-subunit of mitochondrial F1F0 ATP synthase | Energy metabolism (mitochondrial) |

| | | |
|---|---|---|
| VMA2 | Subunit of vacuolar H+-ATPase (proton pump) | Transport |
| MGR1 | Mitochondrial inner membrane protein, quality control (i-AAA protease regulator) | Protein quality control |
| VMA1 | Catalytic subunit of vacuolar ATPase | Transport |
| RTN1 | ER membrane protein shaping tubular ER | Lipid metabolism |
| YSP2 | Sterol-binding protein involved in sterol transport | Lipid metabolism |
| VAC8 | Vacuolar membrane protein involved in inheritance, autophagy | Vesicle trafficking |
| AFG1 | Mitochondrial ATPase involved in ribosome assembly | Protein quality control |
| LAM5 | Sterol transporter at membrane contact sites | Lipid metabolism |
| PHB1 | Prohibitin, mitochondrial membrane scaffold protein | Protein quality control |
| PHB2 | Prohibitin complex subunit | Protein quality control |
| LAM4 | Sterol transporter (ER–PM contact sites) | Lipid metabolism |
| TOM71 | Mitochondrial outer membrane import receptor | Protein targeting/import |
| ATP2 | β-subunit of mitochondrial ATP synthase | Energy metabolism (mitochondrial) |
| MMM1 | ER–mitochondria encounter structure (ERMES) component | Organelle organization |
| LAM6 | Sterol transporter at multiple contact sites | Lipid metabolism |
| ATG26 | Sterol glucosyltransferase in autophagy | Lipid metabolism |
| MGR3 | Regulator of mitochondrial protease (i-AAA complex) | Protein quality control |
| TOM70 | Mitochondrial protein import receptor | Protein targeting/import |
| YlACL1 | ATP-citrate lyase (acetyl-CoA production, Y. lipolytica) | Central carbon metabolism |
| YlACS_YALI1_F08845g | Acetyl-CoA synthetase (Y. lipolytica) | Central carbon metabolism |
| ACS1 | Acetyl-CoA synthetase (non-fermentative growth) | Central carbon metabolism |
| CAB5 | Pantothenate kinase (CoA biosynthesis) | Cofactor & redox metabolism |

| | | |
|---|---|---|
| CAB4 | Phosphopantetheine adenylyltransferase (CoA biosynthesis) | Cofactor & redox metabolism |
| ACS2 | Acetyl-CoA synthetase (fermentative growth) | Central carbon metabolism |
| ERG20_F96W_N127W-trunCrGES | Engineered FPP synthase fused to geraniol synthase (geraniol production) | Strictosidine biosynthesis pathway |
| GgaFPS_N144W | Mutant farnesyl diphosphate synthase favoring GPP formation | Strictosidine biosynthesis pathway |
| AgrGPPS2 | Geranyl diphosphate synthase (GPP production) | Strictosidine biosynthesis pathway |
| CrTDC | Tryptophan decarboxylase (tryptamine formation) | Strictosidine biosynthesis pathway |
| TeG8H | Geraniol 8-hydroxylase (P450) | Strictosidine biosynthesis pathway |
| CrCPR | Cytochrome P450 reductase | Strictosidine biosynthesis pathway |
| CrCYB5 | Cytochrome b5 (electron transfer to P450s) | Strictosidine biosynthesis pathway |
| Cr8HGO | 8-hydroxygeraniol oxidoreductase | Strictosidine biosynthesis pathway |
| NcaMLPLA | Major latex protein-like enzyme (cyclization step) | Strictosidine biosynthesis pathway |
| NmISY2 | Iridoid synthase (nepetalactol formation) | Strictosidine biosynthesis pathway |
| NmISY2_mutH+ | Engineered iridoid synthase (improved activity) | Strictosidine biosynthesis pathway |
| Cr8HGO-NmISY2 | Fusion protein (improved flux between steps) | Strictosidine biosynthesis pathway |
| Cr8HGO-NmISY2_mutH+ | Optimized fusion enzyme | Strictosidine biosynthesis pathway |
| CrIO | Iridoid oxidase (P450) | Strictosidine biosynthesis pathway |
| CrADH2 | Alcohol dehydrogenase | Strictosidine biosynthesis pathway |
| Cr7DLGT | 7-deoxyloganetic acid glucosyltransferase | Strictosidine biosynthesis pathway |
| Cr7DLH | 7-deoxyloganic acid hydroxylase (P450) | Strictosidine biosynthesis pathway |
| CrLAMT | Loganic acid methyltransferase | Strictosidine biosynthesis pathway |
| CrSLS | Secologanin synthase (P450) | Strictosidine biosynthesis pathway |
| CrSTR | Strictosidine synthase (key MIA entry step) | Strictosidine biosynthesis pathway |

| | | |
|---|---|---|
| HsPCIF1 | mRNA cap methyltransferase | Central carbon metabolism regulation |
| CrNPF2.6 | Transporter for MIA intermediates | MIA transporter |
| CrNPF2.9 | Transporter for MIA intermediates | MIA transporter |
| YALI1_A21420p | *Y. lipolytica* homolog from *S. cerevisiae* DAP1 | Unknown |
| YALI1_E05309p | *Y. lipolytica* homolog from *S. cerevisiae* DAP1 | Unknown |
| YALI1_E28648p | *Y. lipolytica* homolog from *S. cerevisiae* DAP1 | Unknown |
| YALI1_E00412p | *Y. lipolytica* homolog from *S. cerevisiae* DAP1 | Unknown |
| YALI1_D04978p | *Y. lipolytica* homolog from *S. cerevisiae* ICE2 | Unknown |
| YALI1_D12147p | *Y. lipolytica* homolog from *S. cerevisiae* ERO1 | Unknown |
| YALI1_E27249p | *Y. lipolytica* homolog from *S. cerevisiae* CPR5 | Unknown |
| YALI1_C14450p | *Y. lipolytica* homolog from *S. cerevisiae* CPR5 | Unknown |
| YALI1_C13890p | *Y. lipolytica* homolog from *S. cerevisiae* CPR5 | Unknown |
| YALI1_C23926p | *Y. lipolytica* homolog from *S. cerevisiae* CPR5 | Unknown |
| YALI1_C09390p | *Y. lipolytica* homolog from *S. cerevisiae* CPR5 | Unknown |
| YALI1_A17450p | *Y. lipolytica* homolog from *S. cerevisiae* CPR5 | Unknown |
| YALI1_C22592p | *Y. lipolytica* homolog from *S. cerevisiae* CPR5 | Unknown |
| YALI1_C21626p | *Y. lipolytica* homolog from *S. cerevisiae* CPR5 | Unknown |
| YALI1_C04553p | *Y. lipolytica* homolog from *S. cerevisiae* HSP26 | Unknown |
| YALI1_D10458p | *Y. lipolytica* homolog from *S. cerevisiae* SSA1 | Unknown |
| YALI1_F32597p | *Y. lipolytica* homolog from *S. cerevisiae* SSA1 | Unknown |
| YALI1_D28273p | *Y. lipolytica* homolog from *S. cerevisiae* SSA1 | Unknown |
| YALI1_A22539p | *Y. lipolytica* homolog from *S. cerevisiae* SSA1 | Unknown |
| YALI1_E00115p | *Y. lipolytica* homolog from *S. cerevisiae* SSA1 | Unknown |

| | | |
|---|---|---|
| YALI1_D36255p | *Y. lipolytica* homolog from *S. cerevisiae* SSA1 | Unknown |
| YALI1_E41647p | *Y. lipolytica* homolog from *S. cerevisiae* SSA1 | Unknown |
| YALI1_E16731p | *Y. lipolytica* homolog from *S. cerevisiae* SSA1 | Unknown |
| YALI1_A00102p | *Y. lipolytica* homolog from *S. cerevisiae* SSA1 | Unknown |
| YALI1_C24707p | *Y. lipolytica* homolog from *S. cerevisiae* SSA1 | Unknown |
| YALI1_D27274p | *Y. lipolytica* homolog from *S. cerevisiae* SSA1 | Unknown |
| YALI1_E33056p | *Y. lipolytica* homolog from *S. cerevisiae* HSP104 | Unknown |
| YALI1_F16656p | *Y. lipolytica* homolog from *S. cerevisiae* HSP104 | Unknown |
| YALI1_B20777p | *Y. lipolytica* homolog from *S. cerevisiae* HSP104 | Unknown |
| YALI1_B16808p | *Y. lipolytica* homolog from *S. cerevisiae* HAC1 | Unknown |
| YALI1_B13069p | *Y. lipolytica* homolog from *S. cerevisiae* UPC2 | Unknown |
| YALI1_E21312p | *Y. lipolytica* homolog from *S. cerevisiae* POS5 | Unknown |
| YALI1_E32908p | *Y. lipolytica* homolog from *S. cerevisiae* POS5 | Unknown |
| YALI1_E28545p | *Y. lipolytica* homolog from *S. cerevisiae* POS5 | Unknown |
| YALI1_E26811p | *Y. lipolytica* homolog from *S. cerevisiae* ZWF1 | Unknown |
| YALI1_D34353p | *Y. lipolytica* homolog from *S. cerevisiae* ALD6 | Unknown |
| YALI1_C04093p | *Y. lipolytica* homolog from *S. cerevisiae* ALD6 | Unknown |
| YALI1_E00588p | *Y. lipolytica* homolog from *S. cerevisiae* ALD6 | Unknown |
| YALI1_F06858p | *Y. lipolytica* homolog from *S. cerevisiae* ALD6 | Unknown |
| YALI1_D10197p | *Y. lipolytica* homolog from *S. cerevisiae* ALD6 | Unknown |
| YALI1_F07847p | *Y. lipolytica* homolog from *S. cerevisiae* TPI1 | Unknown |
| YALI1_F02481p | *Y. lipolytica* homolog from *S. cerevisiae* TPI1 | Unknown |
| YALI1_E20159p | *Y. lipolytica* homolog from *S. cerevisiae* TPI1 | Unknown |

| | | |
|---|---|---|
| YALI1_E35410p | *Y. lipolytica* homolog from *S. cerevisiae* HEM13 | Unknown |
| YALI1_D30196p | *Y. lipolytica* homolog from *S. cerevisiae* HEM14 | Unknown |
| YALI1_F27686p | *Y. lipolytica* homolog from *S. cerevisiae* HEM2 | Unknown |
| YALI1_F25924p | *Y. lipolytica* homolog from *S. cerevisiae* HEM15 | Unknown |
| YALI1_F34051p | *Y. lipolytica* homolog from *S. cerevisiae* HEM3 | Unknown |
| YALI1_C02391p | *Y. lipolytica* homolog from *S. cerevisiae* HEM12 | Unknown |
| YALI1_F25444p | *Y. lipolytica* homolog from *S. cerevisiae* DGK1 | Unknown |
| YALI1_E17431p | *Y. lipolytica* homolog from *S. cerevisiae* CDS1 | Unknown |
| YALI1_D10930p | *Y. lipolytica* homolog from *S. cerevisiae* CHO1 | Unknown |
| YALI1_D04432p | *Y. lipolytica* homolog from *S. cerevisiae* PSD1 | Unknown |
| YALI1_E07347p | *Y. lipolytica* homolog from *S. cerevisiae* CHO2 | Unknown |
| YALI1_E15432p | *Y. lipolytica* homolog from *S. cerevisiae* OPI3 | Unknown |
| YALI1_C00230p | *Y. lipolytica* homolog from *S. cerevisiae* SCT1 | Unknown |
| YALI1_C09773p | *Y. lipolytica* homolog from *S. cerevisiae* AYR1 | Unknown |
| YALI1_A20484p | *Y. lipolytica* homolog from *S. cerevisiae* AYR1 | Unknown |
| YALI1_E22736p | *Y. lipolytica* homolog from *S. cerevisiae* SLC1 | Unknown |
| YALI1_F25951p | *Y. lipolytica* homolog from *S. cerevisiae* ALE1 | Unknown |
| YALI1_B07366p | *Y. lipolytica* homolog from *S. cerevisiae* URA7 | Unknown |
| YALI1_B12742p | *Y. lipolytica* homolog from *S. cerevisiae* CKI1 | Unknown |
| YALI1_D22834p | *Y. lipolytica* homolog from *S. cerevisiae* PCT1 | Unknown |
| YALI1_C08197p | *Y. lipolytica* homolog from *S. cerevisiae* PCT1 | Unknown |
| YALI1_C15420p | *Y. lipolytica* homolog from *S. cerevisiae* CPT1 | Unknown |
| YALI1_E31498p | *Y. lipolytica* homolog from *S. cerevisiae* CPT1 | Unknown |

| | | |
|---|---|---|
| YALI1_B11532p | *Y. lipolytica* homolog from *S. cerevisiae* ERG10 | Unknown |
| YALI1_E13899p | *Y. lipolytica* homolog from *S. cerevisiae* ERG10 | Unknown |
| YALI1_E22238p | *Y. lipolytica* homolog from *S. cerevisiae* ERG10 | Unknown |
| YALI1_F38116p | *Y. lipolytica* homolog from *S. cerevisiae* ERG13 | Unknown |
| YALI1_E05731p | *Y. lipolytica* homolog from *S. cerevisiae* HMG1 | Unknown |
| YALI1_B21004p | *Y. lipolytica* homolog from *S. cerevisiae* ERG12 | Unknown |
| YALI1_E07506p | *Y. lipolytica* homolog from *S. cerevisiae* ERG8 | Unknown |
| YALI1_F08363p | *Y. lipolytica* homolog from *S. cerevisiae* MVD1 | Unknown |
| YALI1_F06018p | *Y. lipolytica* homolog from *S. cerevisiae* IDI1 | Unknown |
| YALI1_E06759p | *Y. lipolytica* homolog from *S. cerevisiae* ERG20 | Unknown |
| YALI1_D10930p | *Y. lipolytica* homolog from *S. cerevisiae* CHO1 | Unknown |
| YALI1_D04432p | *Y. lipolytica* homolog from *S. cerevisiae* PSD1 | Unknown |
| YALI1_E07347p | *Y. lipolytica* homolog from *S. cerevisiae* CHO2 | Unknown |
| YALI1_E15432p | *Y. lipolytica* homolog from *S. cerevisiae* OPI3 | Unknown |
| YALI1_C00230p | *Y. lipolytica* homolog from *S. cerevisiae* SCT1 | Unknown |
| YALI1_C09773p | *Y. lipolytica* homolog from *S. cerevisiae* AYR1 | Unknown |
| YALI1_A20484p | *Y. lipolytica* homolog from *S. cerevisiae* AYR1 | Unknown |
| YALI1_E22736p | *Y. lipolytica* homolog from *S. cerevisiae* SLC1 | Unknown |
| YALI1_F25951p | *Y. lipolytica* homolog from *S. cerevisiae* ALE1 | Unknown |
| YALI1_B07366p | *Y. lipolytica* homolog from *S. cerevisiae* URA7 | Unknown |
| YALI1_B12742p | *Y. lipolytica* homolog from *S. cerevisiae* CKI1 | Unknown |
| YALI1_D22834p | *Y. lipolytica* homolog from *S. cerevisiae* PCT1 | Unknown |
| YALI1_C08197p | *Y. lipolytica* homolog from *S. cerevisiae* PCT1 | Unknown |

| | | |
|---|---|---|
| YALI1_C15420p | *Y. lipolytica* homolog from *S. cerevisiae* CPT1 | Unknown |
| YALI1_E31498p | *Y. lipolytica* homolog from *S. cerevisiae* CPT1 | Unknown |
| YALI1_B11532p | *Y. lipolytica* homolog from *S. cerevisiae* ERG10 | Unknown |
| YALI1_E13899p | *Y. lipolytica* homolog from *S. cerevisiae* ERG10 | Unknown |
| YALI1_E22238p | *Y. lipolytica* homolog from *S. cerevisiae* ERG10 | Unknown |
| YALI1_F38116p | *Y. lipolytica* homolog from *S. cerevisiae* ERG13 | Unknown |
| YALI1_E05731p | *Y. lipolytica* homolog from *S. cerevisiae* HMG1 | Unknown |
| YALI1_B21004p | *Y. lipolytica* homolog from *S. cerevisiae* ERG12 | Unknown |
| YALI1_E07506p | *Y. lipolytica* homolog from *S. cerevisiae* ERG8 | Unknown |
| YALI1_F08363p | *Y. lipolytica* homolog from *S. cerevisiae* MVD1 | Unknown |
| YALI1_F06018p | *Y. lipolytica* homolog from *S. cerevisiae* IDI1 | Unknown |
| YALI1_E06759p | *Y. lipolytica* homolog from *S. cerevisiae* ERG20 | Unknown |

#### Supplementary Table 6. X-ray crystallography data collection and refinement statistics.

| **Protein name**<br>**PDB Entry** | QUI1-quinine complex<br>**36BZ** | CAT2<br>**36CD** |
|---|---|---|
| **Data collection** | | |
| Space group | C 1 2 1 | P 21 21 21 |
| Cell dimensions | | |
| a, b, c (Å) | 94.74, 39.73, 105.62 | 40.78 53.92 173.58 |
| α, β, γ (°) | 90, 113.1, 90 | 90, 90, 90 |
| Resolution (Å) | 48.56 - 1.77 (1.81 - 1.77)* | 45.80 - 2.43 (2.52 - 2.43) |
| $R_{pim}$ | 0.076 (0.472) | 0.069 (0.548) |
| CC ½ $^{Y}$ | 0.987 (0.619) | 0.998 (0.677) |
| I / σ | 11.9 (1.5) | 13.9 (2.2) |
| Completeness (%) | 98.84 (98.29) | 99.9 (99.9) |
| Redundancy | 4.2 (4.2) | 6.3 (6.6) |
| **Refinement** | | |
| Resolution (Å) | 48.56 - 1.77 (1.81 - 1.77) | 43.4 - 2.43 (2.51 - 2.43) |
| No. reflections | 35223 (2470) | 15123 (1354) |
| $R_{work}$ | 0.2244 (0.2753) | 0.2328 (0.2692) |
| $R_{free}^{\pm}$ | 0.2715 (0.3300) | 0.2861 (0.3380) |
| **No. atoms** | 3184 | 2745 |
| Protein | 2898 | 2745 |
| Ligand/ion | 48 | 0 |
| Water | 238 | 0 |
| **B-factors (Å²)** | | |
| Protein | 26.22 | 57.41 |
| Ligand/ion | 32.15 | |
| Water | 30.25 | |
| **R.m.s. deviations** | | |
| Bond lengths (Å) | 0.007 | 0.008 |
| Bond angles (°) | 0.79 | 0.99 |
| **Ramachandran plot** | | |
| Favored | 99.17% | 98.85% |
| Allowed | 0.83% | 1.15% |
| Outliers | 0.00 % | 0.00 % |
| **Molprobity score^** | 1.48 | 1.8 |

*Values for the corresponding parameters in the outermost shell in parenthesis.

$^{Y}CC_{1/2}$ is the Pearson correlation coefficient for a random half of the data, the two numbers represent the lowest and highest resolution shell, respectively.

$^{\pm}R_{free}$ is the $R_{work}$ calculated for about 10% of the reflections randomly selected and omitted from refinement.

^MolProbity score is calculated by combining clashscore with rotamer and Ramachandran percentage and scaled based on X-ray resolution. The percentage is calculated with 100th percentile as the best and 0th percentile as the worst among structures of comparable resolution.

### Supplementary Text

To enable a discussion of secondary structural elements and their relation to the ligand binding pocket, we provided a canonical reference based on the apo wild-type structure (PDB: 3VVX) using DSSP, which reveals eleven helices (α1-α11) including a single anti-parallel β-bridge (T190–V139) and a short $3_{10}$ segment (**Supp. Fig. 4**).

- **CAT2**

For CAT2 (T85C I88C Y92A M143R), we obtained a high-resolution (2.4 Å) apo crystal structure (**Fig. 3B**). Comparison with wild-type RamR revealed that the mutations alone (without ligand) induce a substantial conformational rearrangement: the homodimer C-α RMSD exceeds 4.5 Å, and even single-chain alignment shows significant backbone deviation (**Supp. Fig. 5A**). This indicates that the evolved mutations have remodeled the apo state itself, potentially pre-organizing the pocket for CAT binding.

Because no holo structure was obtained, we cross-referenced Boltz-2 co-folding predictions with Vina docking poses to estimate a binding pose with moderate shape complementarity and meaningful intermolecular interactions (**Supp. Fig. 5B**). Mapping wild-type residues onto this pose rationalizes the key substitutions: I88C and Y92A relieve steric incompatibility with CAT (**Supp. Fig. 5C**), while M143R repositions the sidechain. The arginine swings toward the solvent-exposed surface, vacating space for the ligand (**Supp. Fig. 5D**). PLIP analysis of the immediate contact shell identifies a hydrophobic sandwich formed by L130 and F155, with D152 contributing a salt bridge to CAT (**Supp. Fig. 5E**).

- **MIT2**

For the MIT2 variant (K63I L66M M70V M184T L188I T189C), a high-occupancy ligand density could not be obtained; we therefore used a structural model to rationalize mutations (prepared with Boltz-2).

In the modelled pose K63I, the wild-type lysine sidechain would sterically preclude ligand binding entirely (**Supp. Fig. 6AB**). Additionally, the modelled pose shows stronger shape complementarity and sits directly adjacent to the mutated cluster K63I/L66M/M70V (**Supp. Fig. 6ABC**). M184T seems to alter the long-range hydrophobic interaction with F155 and Y92, potentially loosening the binding pocket for readily ligand access. The modelled pose engages a rich interaction network including a Y59 hydrogen bond to the MIT carbonyl, a π–π stack with F155, and additional hydrophobic contacts (**Supp. Fig. 6C**). Together, these analyses suggest the alternative pose may represent the productive binding mode, although experimental confirmation is needed.

- **QUI1**

QUI1 (K63R L66H M70C) yielded the most informative holo structure: a high-resolution (1.8 Å) holo structure with bound QUI. In contrast to CAT2, the QUI1 backbone closely matches wild-type RamR at both the monomer and homodimer level (C-α RMSD <2 Å, **Supp. Fig. 7A**), indicating that QUI binding is achieved through sidechain-level rearrangements rather than larger conformational change.

The structure reveals a resolved structural-water network in which K63R, D152, R148, and D124 each engage QUI through bridging waters (**Supp. Fig. 7BD**). The K63R guanidinium is itself pre-organized by an adjacent packing arrangement: C70 makes a van der Waals contact with H66, and a water-mediated hydrogen bond from H66 constrains the ε-NH of the R63 sidechain; reducing its conformational entropy and stabilizing the productive rotamer for QUI binding. L66H also leads to a new hydrogen bond (2.9 Å distance) with S91 in the neighbor helix

(**Supp. Fig. 7F**), potentially stabilizing local helix-helix arrangement to position K63R which forms a cation-$\pi$ interaction with the bicyclic aromatic ring of QUI.

This hydrophilic face is opposed by hydrophobic residues on the other side of the pocket, producing an amphiphilic binding site complementary to QUI's amphiphilic nature. The combination of strong shape complementarity and extensive direct vdW contact (**Supp. Fig. 7C**) is consistent with the low-micromolar affinity measured for QUI. PLIP analysis additionally identifies F155 and Y92 as π-stacking partners with QUI along with V138, L188, T189, and M143 forming the hydrophobic face (**Supp. Fig. 7E)**.

- **RAU2**

No experimental structure was obtained for RAU2, so binding poses were estimated by cross-referencing Boltz-2 co-folding and Vina docking poses. Two viable poses emerged, both consistent with the accumulated mutations in RAU2 (T85N I88A Y92V F142M M143S) (**Fig. 3B**, **Supp. Fig. 8AB**). In both poses, T85N is positioned to mediate a water-bridged hydrogen bond to the indoloquinolizidine tertiary amine in pose 1 and to the pyrrole-type indole nitrogen in pose 2. Y92V relieves a steric clash that would otherwise prevent RAU binding in either pose, providing a structural rationale for the low affinity of RAU in WT RamR (**Fig. 3B**).

PLIP analysis of pose 1 highlights a cation–π contact between K63 and RAU, with D152 and R148 engaging the RAU carboxylic acid through hydrogen bonding and/or salt-bridge interactions (**Supp. Fig. 8C**). Pose 2 instead places K62 in contact with the carboxylic acid, with F155 contributing to the canonical π-interaction and D152 hydrogen-bonding to the tertiary amine (**Supp. Fig. 8D**).

To probe how RAU2 discriminates among closely related (hetero)yohimbane alkaloids, we compared RAU with its structural analogs yohimbine (YOH), corynanthine (COR), ajmalicine (AJM), and tetrahydroalstonine (THA). RAU, YOH, and COR share the pentacyclic yohimbane skeleton and differ only at single stereocenters. RAU is the C-20 epimer of YOH and COR is the C-16 epimer of YOH (**Supp. Fig. 9A**), while AJM and THA are heteroyohimbanes in which the saturated E-ring is replaced by a vinyl ether (**Supp. Fig. 9B**).

Superimposing each analog onto the modelled RAU2 pose (aligned on the conserved indole skeleton) identifies V138 and the D152 backbone in helix α10 as the principal steric gates: only RAU and THA are accommodated (**Supp. Fig. 9C**), consistent with the moderate fold-induction observed for THA experimentally, while COR, AJM, and YOH all clash at these positions (**Supp. Fig. 9D**). Visualizing each analog individually against RAU (**Supp. Fig. 10**) isolates the per-ligand clashes. In YOH, the C-17 hydroxyl clashes with V138 and the methyl ester with the D152 backbone, illustrating how a single stereocenter inversion is sufficient to disrupt pocket complementarity. L130, T85N, and C134 emerge as secondary candidates for further investigation of (hetero)yohimbane discrimination for future directed evolution experiments (**Supp. Fig. 9D**).

- **STR1**

STR presents a distinct opportunity among our ligands: the glucose moiety adds many potential hydrogen-bonding donors and acceptors. As with RAU2, no experimental structure was obtained and pose estimation by Boltz-2/Vina cross-referencing revealed a single viable pose (**Fig. 3B**, **Supp. Fig. 11A**).

The modelled pose rationalizes each STR1 mutation through a coherent mechanism centered on helix α8. C134K introduces a long, flexible side chain that reaches toward the glucose hydroxyl while making hydrophobic contacts with the STR indole (**Supp. Fig. 11B**). S137P prematurely terminates α8, perturbing the distal β-bridge involving V138 (**Supp. Fig. 4A**); this in turn may reposition V138 toward STR, contributing to a more complementary hydrophobic pocket (Figure x9B). L133V (also in α8) likely allows the helix to pack inward, tightening the pocket (**Supp. Fig. 11C**). PLIP analysis confirms the expected involvement of D152 and R148 in hydrogen-bonding the STR tertiary amine, while V138, K134, T85, F155, and Y93 form a hydrophobic surface complementary to the indole moiety (**Supp. Fig. 11D**). A rotated view (**Supp. Fig. 11E**) shows complementary interactions with the glucose moiety.

**Supplementary Material References**

Babaei, M., Sartori, L., Karpukhin, A., Abashkin, D., Matrosova, E., & Borodina, I. (2021). Expansion of EasyClone-MarkerFree toolkit for Saccharomyces cerevisiae genome with new integration sites. *FEMS Yeast Research*, *21*(4), foab027. https://doi.org/10.1093/femsyr/foab027

d'Oelsnitz, S., Kim, W., Burkholder, N. T., Javanmardi, K., Thyer, R., Zhang, Y., Alper, H. S., & Ellington, A. D. (2022). Using fungible biosensors to evolve improved alkaloid biosyntheses. *Nature Chemical Biology*, *18*(9), Article 9. https://doi.org/10.1038/s41589-022-01072-w

Deichmann, M., Schiesaro, G., Ramanathan, K., Zeeberg, K., Koefoed, N. M. T., Ormhøj, M., Friis, R. U. W., Gill, R. T., Hadrup, S. R., Jensen, E. D., & Jensen, M. K. (2025). A yeast surface display platform for characterizing CAR T cell responses to cancer antigens. *Nature Communications*, *16*(1), 10306. https://doi.org/10.1038/s41467-025-65236-7

Holtz, M., Rago, D., Nedermark, I., Hansson, F. G., Lehka, B. J., Hansen, L. G., Marcussen, N. E. J., Veneman, W. J., Ahonen, L., Wungsintaweekul, J., Acevedo-Rocha, C. G., Dirks, R. P., Zhang, J., Keasling, J. D., & Jensen, M. K. (2024). Metabolic engineering of yeast for *de novo* production of kratom monoterpene indole alkaloids. *Metabolic Engineering*, *86*, 135–146. https://doi.org/10.1016/j.ymben.2024.09.011

Jessop-Fabre, M. M., Jakočiūnas, T., Stovicek, V., Dai, Z., Jensen, M. K., Keasling, J. D., & Borodina, I. (2016). EasyClone-MarkerFree: A vector toolkit for marker-less integration of genes into Saccharomyces cerevisiae via CRISPR-Cas9. *Biotechnology Journal*, *11*(8), 1110–1117. https://doi.org/10.1002/biot.201600147

Lee, M. E., DeLoache, W. C., Cervantes, B., & Dueber, J. E. (2015). A Highly Characterized Yeast Toolkit for Modular, Multipart Assembly. *ACS Synthetic Biology*, *4*(9), 975–986. https://doi.org/10.1021/sb500366v

Li, S., Si, T., Wang, M., & Zhao, H. (2015). Development of a Synthetic Malonyl-CoA Sensor in Saccharomyces cerevisiae for Intracellular Metabolite Monitoring and Genetic Screening. *ACS Synthetic Biology*, *4*(12), 1308–1315. https://doi.org/10.1021/acssynbio.5b00069

Spinner, A., Sreenivasan, S., McLellan, J. R., Ikonomova, S., Cortade, D., d'Oelsnitz, S., Sheldon, K., Vasilyeva, O., Alperovich, N. Y., Chadha, A., Nematollahi, L., Dhroso, A., Sisson, Z., Hudson, C. M., DeBenedictis, E., Kelly, P. J., Apel, A. R., Ross, D., & Baranowski, C. (2026). *GROQ-seq Datasets*

*Across Transcription Factors (LacI, RamR, VanR), T7 RNA Polymerase and TEV Protease* (p. 2026.04.15.718744). bioRxiv. https://doi.org/10.64898/2026.04.15.718744

Wang, H., Jiang, G., Liang, N., Dong, T., Shan, M., Yao, M., Wang, Y., Xiao, W., & Yuan, Y. (2023). Systematic Engineering to Enhance 8-Hydroxygeraniol Production in Yeast. *Journal of Agricultural and Food Chemistry*, *71*(10), 4319–4327. https://doi.org/10.1021/acs.jafc.2c09028

Zhang, J., Hansen, L. G., Gudich, O., Viehrig, K., Lassen, L. M. M., Schrübbers, L., Adhikari, K. B., Rubaszka, P., Carrasquer-Alvarez, E., Chen, L., D'Ambrosio, V., Lehka, B., Haidar, A. K., Nallapareddy, S., Giannakou, K., Laloux, M., Arsovska, D., Jørgensen, M. A. K., Chan, L. J. G., … Keasling, J. D. (2022). A microbial supply chain for production of the anti-cancer drug vinblastine. *Nature*, *609*(7926), Article 7926. https://doi.org/10.1038/s41586-022-05157-3

Zhang, Y., Yuan, M., Wu, X., Zhang, Q., Wang, Y., Zheng, L., Chiu, T., Zhang, H., Lan, L., Wang, F., Liao, Y., Gong, X., Yan, S., Wang, Y., Shen, Y., & Fu, X. (2023). The construction and optimization of engineered yeast chassis for efficient biosynthesis of 8-hydroxygeraniol. *mLife*, *2*(4), 438–449. https://doi.org/10.1002/mlf2.12099
